# Microbiota from young mice counteracts susceptibility to age-related gout through modulating butyric acid levels in aged mice

**DOI:** 10.1101/2024.04.22.590443

**Authors:** Ning Song, Hang Gao, Jianhao Li, Yi Liu, Mingze Wang, Zhiming Ma, Naisheng Zhang, Wenlong Zhang

## Abstract

Gout is a prevalent form of inflammatory arthritis that occurs due to high levels of uric acid in the blood leading to the formation of urate crystals in and around the joints, particularly affecting the elderly. Recent research has provided evidence of distinct differences in the gut microbiota of patients with gout and hyperuricemia when compared to healthy individuals. However, the link between gut microbiota and age-related gout remained underexplored. Our study found that gut microbiota plays a crucial role in determining susceptibility to age-related gout. Specifically, we observed that age-related gut microbiota regulated the activation of the NLRP3 inflammasome pathway and modulated uric acid metabolism. More scrutiny highlighted the positive impact of "younger" microbiota on the gut microbiota structure of old or aged mice, enhancing butanoate metabolism and butyric acid content. Experimentation with butyrate supplementation indicated that butyric acid exerts a dual effect, inhibiting inflammation in acute gout and reducing serum uric acid levels. These insights emphasize the potential of gut microbiome rejuvenation in mitigating senile gout, unraveling the intricate dynamics between microbiota, aging, and gout. It potentially serves as a therapeutic target for senile gout-related conditions.

## Introduction

Gout, the most common inflammatory arthritis in elderly individuals, results from the deposition of monosodium urate crystals in articular and nonarticular structures(1), particularly among individuals aged 75-84 years, and the occurrence rate in this population can reach 4%(2). The development of gout is primarily attributed to the significant risk posed by a high serum urate concentration(2). Pathological hyperuricemia is defined by a serum urate concentration exceeding 408 μmol/L [6.8 mg/dL], which forms monosodium urate crystals in vitro at physiological pH and temperature(3). The reason why the elderly population commonly experiences gout is complicated by the fact that the population is also ageing, and the treatment of this disease is often intricate due to the presence of comorbidities and medications prescribed for concurrent conditions(2). Although the basic principles for the prevention and treatment for gout remain the same across different age groups, elderly individuals often exhibit lower tolerance for medication dosages, types, side effects, and surgical procedures due to their physiological factors. Moreover, despite the increasing severity of gout among elderly individuals, research on this issue remains scarce.

The gut microbiota plays essential roles in regulating energy and metabolism, as revealed by recent studies(4, 5), which have shown that individuals with hyperuricemia and gout exhibit dysbiosis of the gut microbiota(6, 7). Moreover, because the gut microbiota can directly participate in the metabolism of purines and uric acid(8), it may play a crucial role in gout and hyperuricemia development. Although numerous studies have examined the relationship between the microbiome and early life stages, the impact of the microbiome’s on ageing and frailty in later life needs to be explored. Furthermore, studies have shown that the structure of the gut microbiota undergoes a gradual "ageing" process with advancing age, and this process is characterized by a decrease in the microbial diversity(9, 10). In addition, transplantation of a young microbiota can improve central nervous system inflammation and retinal inflammation in aged mice(11), and counteract age-related behavioral deficits(12). We hypothesize that the high prevalence of gout in the elderly population may be closely related to its “ageing” gut microbiota. The association between the "ageing" gut microbiota and gout in elderly individuals has not been reported. Hence, we conducted a study to elucidate the influence of the ageing gut microbiota on the occurrence and progression of gout in elderly individuals.

In this study, we first tested the sensitivity to monosodium urate (MSU) in different age groups and conducted a microbiota clearance (Abx) on mice of different age groups to assess their sensitivity to MSU again. At the same time, we also tested their serum uric acid levels. We found that sensitivity to MSU increased with age, but changed after clearing the gut microbiota in terms of sensitivity to MSU and serum uric acid levels. Then we performed cross-age fecal microbiota transplantation (FMT) and subsequently stimulated the mice with MSU with the aim of how mice belonging to different age groups exhibit sensitivity to MSU after undergoing FMT. Because hyperuricemia is a necessary physiological factor for gout, we also evaluated the expression levels of uric acid-producing enzymes and uric acid transport proteins in the mice. Surprisingly, transplantation of the gut microbiota from aged mice into young mice, significantly increased their sensitivity to MSU. Conversely, the transplantation of the gut microbiota from young mice into aged mice, significantly decreased their sensitivity to MSU. These findings suggest that the gut microbiota of older individuals plays a promoting role in the occurrence and progression of gout. Moreover, an analysis of the serum uric acid levels of the mice after cross-age FMT yielded similar results. To investigate the underlying mechanisms of how gut microbiota influences gout and hyperuricemia, we performed 16S rDNA sequencing and untargeted metabolomics analysis of fecal samples. We then, observed a significant increase in the abundance of Bifidobacterium and Akkermansia in the gut microbiota of young mice and old or aged mice after transplantation of the gut microbiota from young mice. To further investigate the potential biological pathways affected by the gut microbiota, we performed functional metagenomic analysis using Tax4FUN software and found that butanoate metabolism was more robust in young mice than in aged mice. Furthermore, we also observed that transplantation of the gut microbiota from young mice to aged mice enhanced the butanoate metabolism of the recipient mice. Due to the limitations of untargeted metabolomics, we did not observe any differences in the levels of butyric acid among the different groups. However, the pathway data obtained by fecal untargeted metabolomics also yielded similar results. Based on the above-described results, we hypothesize that butyrate may play a significant role in these processes. Excitingly, the results from a short-chain fatty acid (SCFA) analysis support our hypothesis. Furthermore, a supplementation experiment using butyrate revealed, that the results aligned well with the cross-age FMT findings, which suggests that transplantation of the gut microbiota from young mice into aged mice can effectively prevent gout and hyperuricemia, and that butyrate is likely the critical factor playing a crucial role. Our research findings demonstrate the potential of young gut microbiota in preventing gout in elderly individuals and provide new insights and perspectives for the prevention and treatment of gout in elderly individuals.

## Results

### Gout susceptibility increases with age, related to gut microbiota

In this study, we used the male C57BL/6 mice belonging to three age groups: young (∼3 months), old (∼18 months) and aged (∼24-months) mice (Fig. 1a). To investigate the impact of age on gout, we used a mouse model in which subcutaneous injections of MSU crystals were administered to the dorsal aspect of the hind paws (13–15). The result showed that the old group exhibited a significant increase in footpad swelling compared with the young group, and the levels of IL-1β was significantly elevated in the Old and Aged groups (Fig. 1b, c). The levels of IL-6 was significantly elevated in the Old and Aged groups, after MSU stimulated. While no significant difference in foot tissue TNF-α concentration was found between Aged and Young groups, an significant upward trend was observed in Old group(supplementary material 4 part one). A prominent correlation was found between the occurrence of gout and the concentration of serum uric acid, which exhibits an upwards trend with advancing age(2). Thus, we also measured the serum uric acid level, we found uric acid levels significantly increase with age (Fig. 1d). There are reports indicating a close relationship between gut microbiota and gout(6). Therefore, we simultaneously tested the gut microbiota of mice at different ages. The results showed that as age increases, the ASVs (Amplicon Sequence Variants) showed a declining trend (Fig. 1e). Moreover, the principal coordinates analysis (PCoA) results revealed distinctive differences in the phylogenetic community structures between these groups (Fig. 1f). Then we carried out antibiotics (ABX) cocktail on mice of different age groups, found changes in the footpad swelling, IL-1β and discovered no differences in the level of serum uric acid (Fig. 1g-i). There was no significant difference in the content of IL-1β in the foot tissues of mice of all ages without MSU stimulation, whether treated with antibiotics or untreated (supplementary material 4 part two). These data highlight indicates that aging-associated changes in the gut microbiota exacerbate gout attacks.

**Fig 1.**
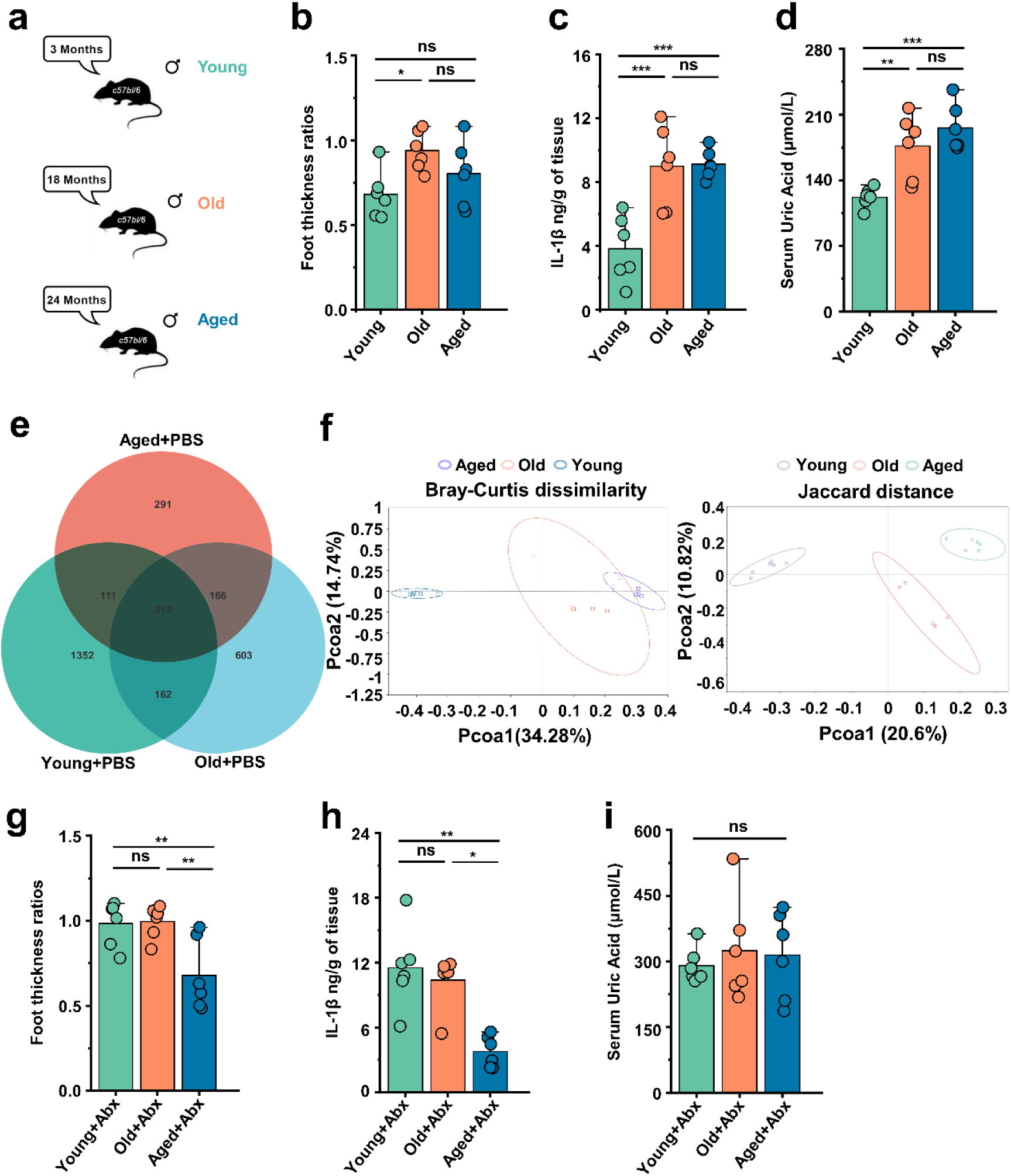
Gout susceptibility increases with age, related to gut microbiota. (a) Mice of different ages. (b-d) The foot thickness ratios (b), foot tissue’s IL-1β concentrations (c) and serum concentrations of uric acid of three different age ranges groups (d) were tested after MSU administration (n=6). (e-f) The three different age ranges groups’ ASVs (Amplicon Sequence Variants) (e) and PcoA analysis (using Bray-Curtis dissimilarity and Jaccard distance) (f). (g-i) The foot thickness ratios (g), foot tissue’s IL-1β concentrations (h) and serum concentrations of uric acid of three different age ranges groups (i) (treated with antibiotics (ABX) cocktail) were tested after MSU administration (n=6). Values are presented as the mean ± SEM. Differences were assessed by One-Way ANOVA and denoted as follows: *p < 0.05, **p < 0.01, and ***p < 0.001, “ns” indicates no significant difference between groups.

### Aged-to-young FMT worsens acute gout, whereas young-to-aged FMT reduced this disease

Based on the above results, we conducted fecal microbiota exchanges between male C57BL/6 mice also belonging to three age groups: young (∼3 months), old (∼18 months), and aged (∼24-months) mice. The experimental design and timeline are presented in Figure 2a. To investigate the impact of cross-age FMT on gout, we used a gout mouse model as aforementioned. The results of supplementary figure 1a showed that young mice transplanted with fecal microbiota from mice in the old or aged group (Young+Old or Young+Aged) exhibited a significant increase in footpad swelling compared with the control group (Young+PBS). However, no significant difference in footpad swelling was found between old or aged mice transplanted with fecal microbiota from mice in the young groups (Old+Young and Aged+Young) and the control groups (Old+PBS and Aged+PBS) (supplementary figure 1a). Subsequently, we conducted a comparative analysis of haematoxylin & eosin (H&E)-stained tissue sections (scar, 1000μm), derived from mice with cross-age FMT and MSU-induced acute gout. We found that the Young+Old and Young+Aged group showed more inflammatory cell infiltration in the subcutaneous soft tissues than the Young+PBS group, whereas the Old+Young and Aged+Young groups showed less inflammatory cell infiltration (Fig. 2b). The inflammatory factors in the foot tissue of the mice with cross-age FMT were then examined. The levels of IL-1β, IL-6, and TNFα were significantly elevated in the Young+Aged group and significantly lower in the Aged+Young group compared with the Aged+PBS group (Fig. 2c-e). Meanwhile, to investigate the impact of cross-age FMT on MSU-induced inflammation, we utilized an animal model of C57BL/6 mice administered an intraperitoneal injection of 2.5 mg of MSU (16). Six hours after MSU intraperitoneal injection, we washed the mouse peritoneal cavity with sterile PBS and collected the peritoneal fluid and peritoneal cells separately. Simultaneously, we also examined the levels of inflammatory cytokines in mouse serum. Consistent with the acute gout model, the Young+Old or Young+Aged group exhibited more pronounced activation of IL- 1β, IL-6, and TNF-α in peritoneal fluid (Fig. 2f-h) and serum (Supplementary figure 1b-c). Although no differences in the IL-6 levels of serum and peritoneal fluid was found between the Old+PBS and Old+Young groups or between the Aged+Young and Aged+PBS groups (supplementary figure 1c and Fig. 2g), a significant reduction in IL-1β and TNF-α was observed (supplementary figure 1b, d and Fig. 2f, h). These findings suggest that the increased susceptibility to gout in the elderly may closely related to “aging” gut microbiota.

**Fig 2.**
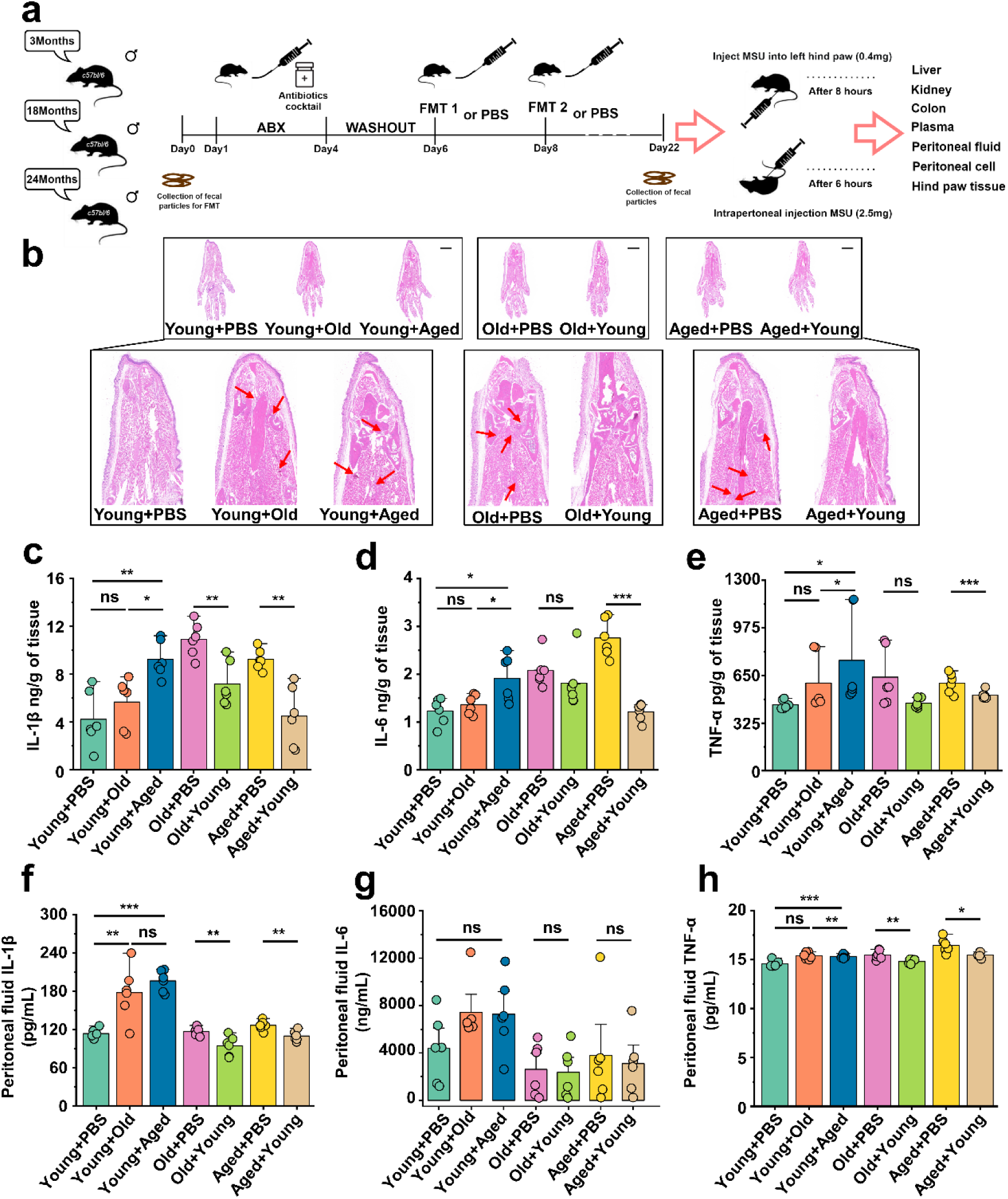
Aged-to-young FMT worsens acute gout, whereas young-to-aged FMT reduced this disease. (a) Overall experimental design and timeline for experiments. (b) Representative H&E-stained images of left foot tissues. Scale bars 1000 μm and 3x magnification. (c-e) Foot tissue inflammatory parameters, including IL-1β (c), IL-6 (d) and TNF-α (e) concentrations, from the indicated mice are shown (n=6). (f-h) The peritoneal fluid concentrations of IL-1β (f), IL-6 (g) and TNF-α (h) inflammatory parameters were measured in the indicated mice(n=6). Values are presented as the mean ± SEM. Differences were assessed by t-test or One-Way ANOVA and denoted as follows: *p < 0.05, **p < 0.01, and ***p < 0.001, “ns” indicates no significant difference between groups.

### “Younger” gut microbiota suppresses NLRP3 inflammasome pathway, “aging” gut microbiota promotes

The pathogenesis of acute gout has been primarily linked to the activation of proinflammatory pathways, notably NLRP3, and this activation instigates a surge in the production of the inflammatory cytokines, which further underscores their pivotal roles in the progression of the disease’s (17). We primarily examined proteins of foot tissue and peritoneal cells associated with the NLRP3 inflammasome pathway. Interestingly, we discovered that the Young+Aged group demonstrated more pronounced activation of NLRP3, Pro-Caspase-1, Caspase-1, and IL-1β compared with the control group (Fig. 3a-b). However, the Old+Young and Aged+Young groups showed lower protein levels of Caspase-1 and IL-1β compared with the control group (Fig. 3c-f), which suggests that the gut microbiota of elderly mice may make them more sensitive to MSU stimulation, whereas the gut microbiota of young mice can effectively resist the MSU stimulation, and inhibit Caspase-1 cleavage, and IL-1β secretion. Meanwhile, the NLRP3 inflammasome pathway also was investigated in peritoneal cells, and similar results were obtained in the gout model. The fecal microbiota from mice in the old or aged group exacerbated the activation of Pro-Caspase -1, Caspase-1, Pro-IL-1β, and IL-1β in peritoneal cells, increasing the production of IL-1β (Supplementary figure 2a-b). In contrast, the fecal microbiota from mice in the young group inhibited the cleavage of Caspase-1 and the secretion of IL-1β (Supplementary figure 2c-f).

**Fig 3.**
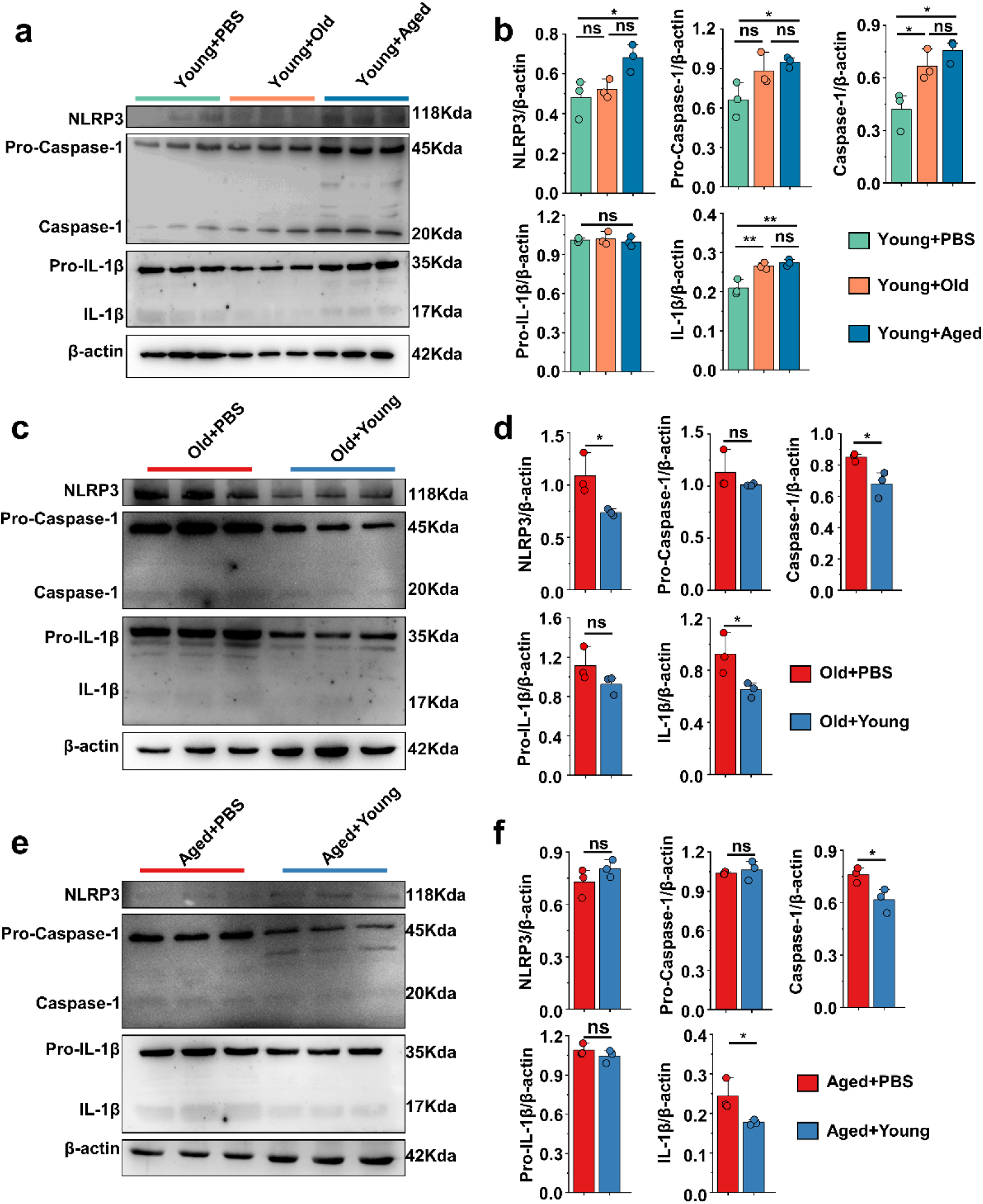
“Younger” gut microbiota suppresses NLRP3 inflammasome pathway, “aging” gut microbiota promotes. (a-b) Representative western blot images and band density (Young+PBS, Young+Old and Young+Aged) of foot tissue NLRP3 pathways proteins (n=3). (c-d) Representative western blot images and band density (Old+PBS and Old+Young) of foot tissue NLRP3 pathways proteins (n=3). (e-f) Representative western blot images and band density (Aged+PBS and Aged+Young) of foot tissue NLRP3 pathways proteins (n=3). Values are presented as the mean ± SEM. Differences were assessed by t-test or One-Way ANOVA and denoted as follows: *p < 0.05, **p < 0.01, and ***p < 0.001, “ns” indicates no significant difference between groups.

### Beneficial effects of FMT from young to aged mice on uric acid metabolism

Serum samples were collected from the the different groups, and their serum uric acid levels were initially assessed. Surprisingly, we observed an elevation in the average serum uric acid levels in the Young+Old or Young+Aged group, whereas the Old+Young or Aged+Young group exhibited decreases in the serum uric acid levels compared with the same-age control group (Fig. 4a). Our study, showed that compared with those of the same-age control group, the serum levels of AST and ALT were elevated in the Young+Old and Young+Aged groups, although the differences were not significant, whereas the Aged+Young groups exhibited modest decreases in these levels (Supplementary figure 3a-b). Furthermore, similar trends were found for indicators (Crea and BUN) of renal function (Supplementary figure 3c-d). Hyperuricemia is attributed to increased uric acid synthesis and decreased uric acid excretion in the body. We first measured the activity of enzymes involved in uric acid synthesis, namely adenosine deaminase (ADA), guanine deaminase (GDA), and xanthine dehydrogenase (XOD), in the liver and that of (XOD) in the kidney. The activity of ADA, GDA, and XOD in the liver, and the activity of XOD in the kidney of the Young+Old and Young+Aged groups did not significantly differ from those of the same-age control groups (Fig. 4b-e). The fecal microbiota from mice in the young group induced a notable decrease in the activity of enzymes related to uric acid synthesis, and the most prominent reductions were found for ADA and XOD activity in the liver and XOD activity in the kidney (Fig. 4b, d and e). We then examined the mRNA levels of relevant proteins involved in uric acid transport. We initially measured the renal injury marker KIM-1, and observed that the fecal microbiota from mice in the young group contributed to attenuating the mRNA expression levels of this marker compared with the same-age control group (Fig. 4f). The urate transporter URAT1, which is responsible for uric acid reabsorption, exhibited a similar trend (Fig. 4g). However, cross-age FMT did not significantly impact the mRNA expression of another uric acid reabsorption protein, GLUT9 (Fig. 4h). The Young+Aged group showed lower mRNA expression levels of the uric acid excretion proteins OAT1 and OAT3 compared with the Young+PBS group (Fig. 4i-j). Although the mRNA expression levels of OAT1 and OAT3 did not show significant differences, a significant increasing trend was found for the other uric acid excretion protein, ABCG2, in aged mice transplanted with fecal microbiota from mice in the young group exhibited a significant increasing trend (Fig. 4k).

**Fig 4.**
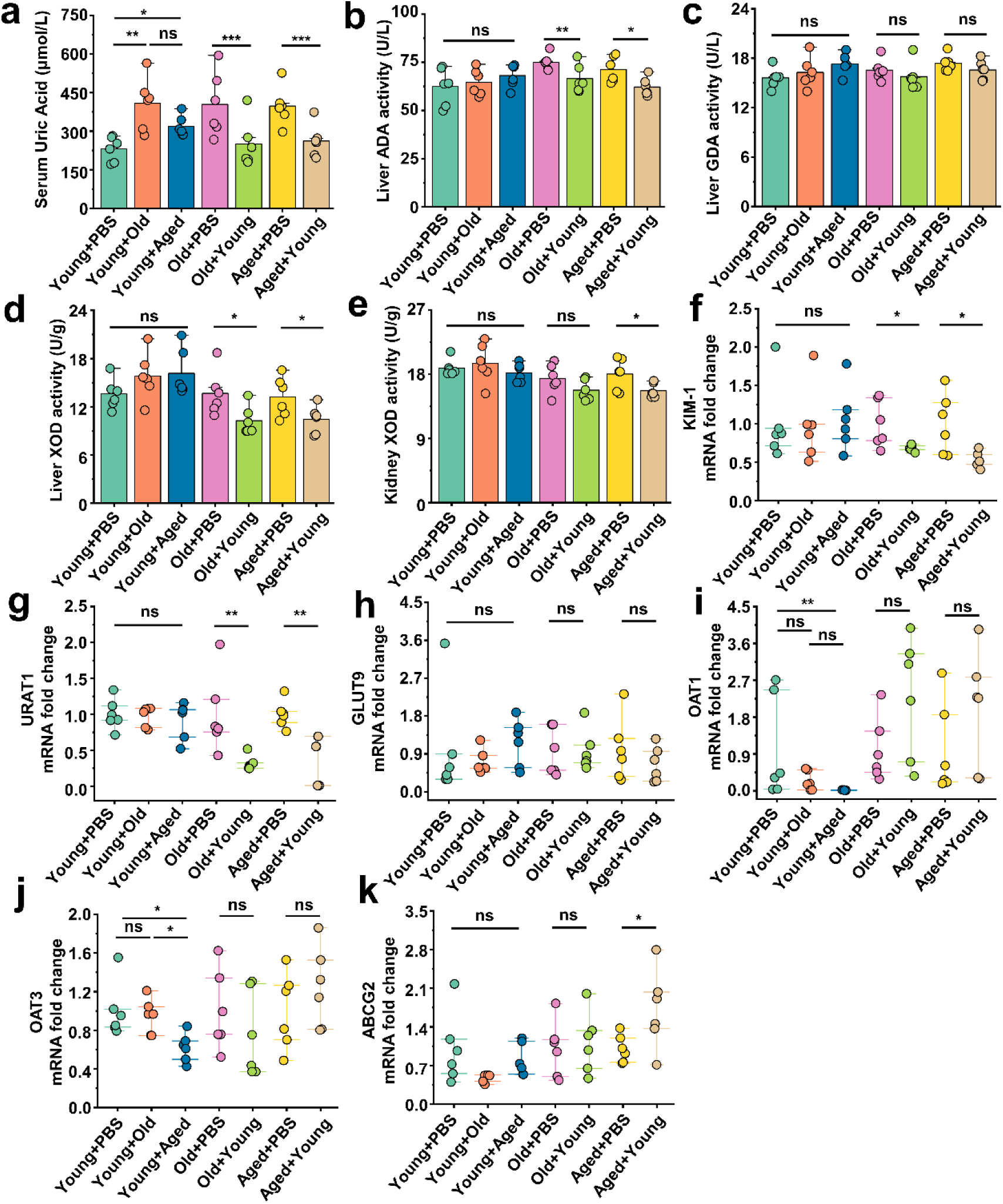
Beneficial effects of FMT from young to aged mice on uric acid metabolism. (a) All groups’ serum concentrations of uric acid (n=6). (b-d) The activity of uric acid-producing enzymes of liver in the cross-age fecal microbiota transplantation group and its control group (n=6), including ADA (b), GDA (c) and XOD (d). (e) The activity of XOD of kidney in the cross-age fecal microbiota transplantation group and its control group (n=6). (f) Relative kidney injury molecule-1 (KIM-1) expression in the indicated groups by qPCR (n = 6). (g and h) Relative renal genes for uric acid reabsorption expression in the indicated groups by qPCR (n = 6), including URAT1 (g) and GLUT9 (h). (i-k) Relative renal genes for uric acid excretion expression in the indicated groups by qPCR (n = 6), including OAT1 (i), OAT3 (j) and ABCG2 (k). Values are presented as the mean ± SEM. Differences were assessed by t-test or One-Way ANOVA and denoted as follows: *p < 0.05, **p < 0.01, and ***p < 0.001, “ns” indicates no significant difference between groups.

Because ABCG2 is also expressed in the intestine, we examined its mRNA expression levels in the colon and found that the fecal microbiota from mice in the aged group could inhibit its expression (supplementary figure 3e). Similar trends were found for the mRNA expression levels of mice in ZO-1 and JAMA in the colon of the Young+Aged group (Supplementary figure 3g-h). No significant alterations in the colonic Occludin mRNA expression levels were observed among all the groups (supplementary figure 3f). However, the mRNA expression levels of JAMA in the Old+Young and Aged+Young groups were significantly different from those in the same-age control groups (supplementary figure 3h). Our study, showed that compared with those of the same-age control group, the colon levels of ZO-1 and Occludin protein were significantly decreases in the Young+Old and Young+Aged groups, whereas the Old+Young and Aged+Young groups exhibited modest elevated in these levels (supplementary material 4 part three).These results suggests that the fecal microbiota from mice in the old or aged group leads to insufficient uric acid excretion, whereas the fecal microbiota from mice in the young group promotes uric acid elimination, inhibits reabsorption, and may contribute to the integrity of the intestinal barrier structure and the maintenance of normal physiological function.

### Modifications in the gut microbiota composition following cross-age FMT

To characterize the age-related alterations in the gut microbiota and assess changes following transplantation, we conducted 16S rDNA amplicon sequencing of all the mice at the endpoint. We focused on investigating the gut microbiota of mice after cross-age FMT. We detected the relative abundance of the top 10 phylum levels in each group (Fig. 5a) and found no significant differences in the abundance of Bacteroidetes and Firmicutes in the transplanted groups compared with that of their respective control groups (Supplementary figure 4a-b). No significant differences were found in the the ratio of Firmicutes to Bacteroidetes (supplementary figure 4c). The species richness, evenness, and rarity are fundamental components of biodiversity and are commonly quantified using indices such as Chao1, observed otus, Shannon, and Simpson indices. Although no significant differences in the Simpson index were found among these groups (supplementary figure 4e), and the indices of the Old+Young and Aged+Young groups were not significantly different from those of the Old+PBS and Aged+PBS groups, respectively, the Chao 1 (Fig. 5b), observed_otus (Fig. 5c), and Shannon (supplementary figure 4d) indices of the Young+Aged group were significantly lower than those of the Young+PBS group. These findings are in line with other studies(18), indicating declines in the richness and diversity of the gut microbiota during aging. Moreover, the principal coordinates analysis (PCoA) results revealed distinctive differences in the phylogenetic community structures between these groups. We showed that the Young+Old and Young+Aged groups tended to be closer to the Old+PBS and Aged+PBS groups, and the Old+Young and Aged+Young groups tended to be closer to the Young+PBS group (Fig. 5d). We then compared the abundance of the top 15 genus level among all the groups (supplementary figure 4f). A Metastats analysis, found that the abundance of Bifidobacterium significantly differed between the Young+PBS group and the Young+Old and Young+Aged groups, and that the abundance of Lachnoclostridium showed an increasing trend in the Old+Young and Aged+Young groups (Fig. 5e). The ternary plot showed that Akkermansia appears to be the dominant species in the Young+PBS, Old+Young and Aged+Young group (Fig. 5f). According to previous reports(19–22) on Bifidobacterium and Akkermansia, we hypothesize that these genera or their metabolites may play a key role in resistance to gout and hyperuricemia. Based on recent research findings(23) and a KEGG analysis of bacterial community functions, we discovered that butanoate metabolism was more robust in the Young+PBS group than in the Old+PBS and Aged+PBS groups (Table 2a-b). However, no similar phenomenon was observed in the Young+Old and Young+Aged groups compared with the Young+PBS group (Table 2c-d). More interestingly, the Old+Young and Aged+Young groups showed stronger butanoate metabolism than the Old+PBS and Aged+PBS groups, respectively (Table 2e-f). Considering the results from the analysis modifications in the gut microbiota composition after cross-age FMT, both Bifidobacterium and Akkermansia metabolites include SCFAs. These findings indicate that butyric acid derived from the young microbiome may be the critical element responsible for controlling gout. Numerous studies have reported the beneficial effects of butyric acid, but further research on the relationship between butyric acid and gout is needed.

**Fig 5.**
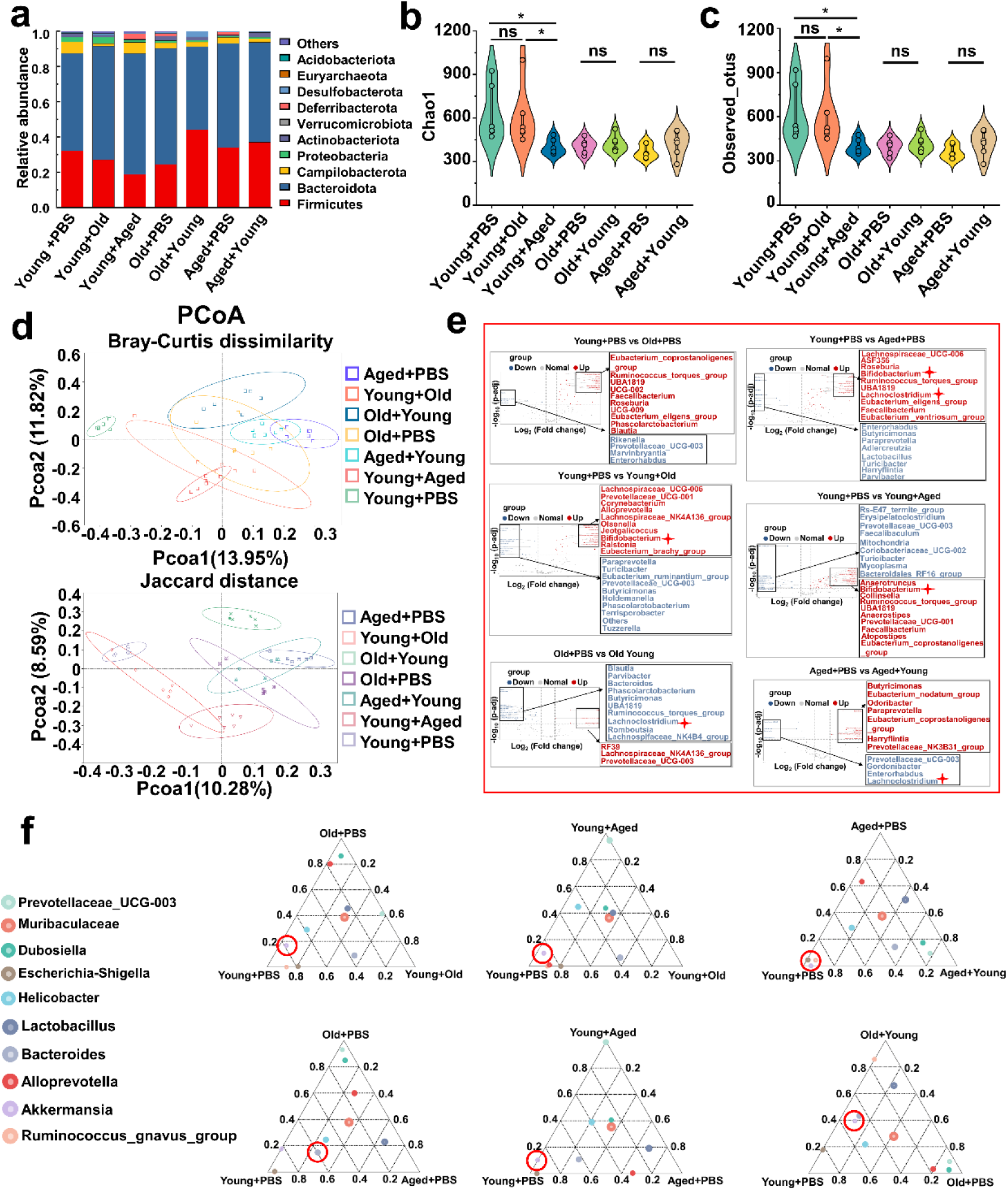
Modifications in the gut microbiota composition following cross-age FMT. (a) Bacterial composition at the phylum levels (top 10) of the indicated groups (n=6). (b-c) Alpha diversity indices including Chao1 (b) and observed_otus (c) index in the indicated groups (n=6). (d) β-diversity difference among the seven groups analyzed by the PcoA using Bray-Curtis dissimilarity and Jaccard distance. (e) Volcano Plot of inter-group significance analysis using metastat (t-test, p < 0.05). (f) The ternary plot of three different groups among the seven groups at genus levels (top 10).

### Changes in fecal microbiota metabolism and pathways after cross-age FMT

We also performed an untargeted metabolomics analysis of samples collected after cross-age FMT to investigate the research question. Chemical classification of the metabolites identified in this study was performed, and a pie chart was generated to reflect the distribution and numbers of metabolites in each category. The pie chart for Class I metabolite classification and KEGG pathway annotation is shown in supplementary figure 5a. Among the 42 samples analysed in this study, a total of 1009 metabolites were identified in the positive ion mode, and 522 metabolites were identified in the negative ion mode among the 42 samples analyzed in this project. The volcano plot provides an overview of the distribution of differentially expressed metabolites (supplementary figure 5b). We also used the KEGG database for metabolic analysis and network research of the identified biological entities. The enrichment results are based on KEGG pathway units, and hypergeometric testing was performed to identify pathways enriched in differentially abundant metabolites compared with all specified metabolite backgrounds. Through pathway enrichment, we discerned and elucidated the principal biochemical metabolic pathways and signal transduction pathways in which the differentialy abundant metabolites actively participate. The metabolites showing differential abundance between the Young+PBS and Old+PBS groups were enriched in the secondary bile acid biosynthesis pathway (supplementary figure 6a). Interestingly, the metabolites showing differential abundance between the Young+PBS and Aged+PBS groups were also enriched in the butanoate metabolism pathway (supplementary figure 6b). The same phenomenon was found from the comparisons of the Young+PBS and Young+Old groups, Young+PBS and Young+Aged groups, Old+PBS and Old+Young groups, and Aged+Young and Aged+PBS groups: the metabolites identified as showing differential abundance between the groups metabolites were enriched in the butanoate metabolism pathway (Fig. 6a-d). Furthermore, we found that the metabolites showing differential abundance between the Old+PBS and Old+Young groups and between the Aged+Young and Aged+PBS groups were enriched in the secondary bile acid biosynthesis pathway (Fig. 6c-d). Due to the limitations of untargeted metabolomics, we did not observe any differences in butyric acid. However, the results were consistent with the predicted functional capabilities of microbial communities and corresponded to butanoate metabolism. Based on the abovementioned results, butyrate may play an important role.

**Fig 6.**
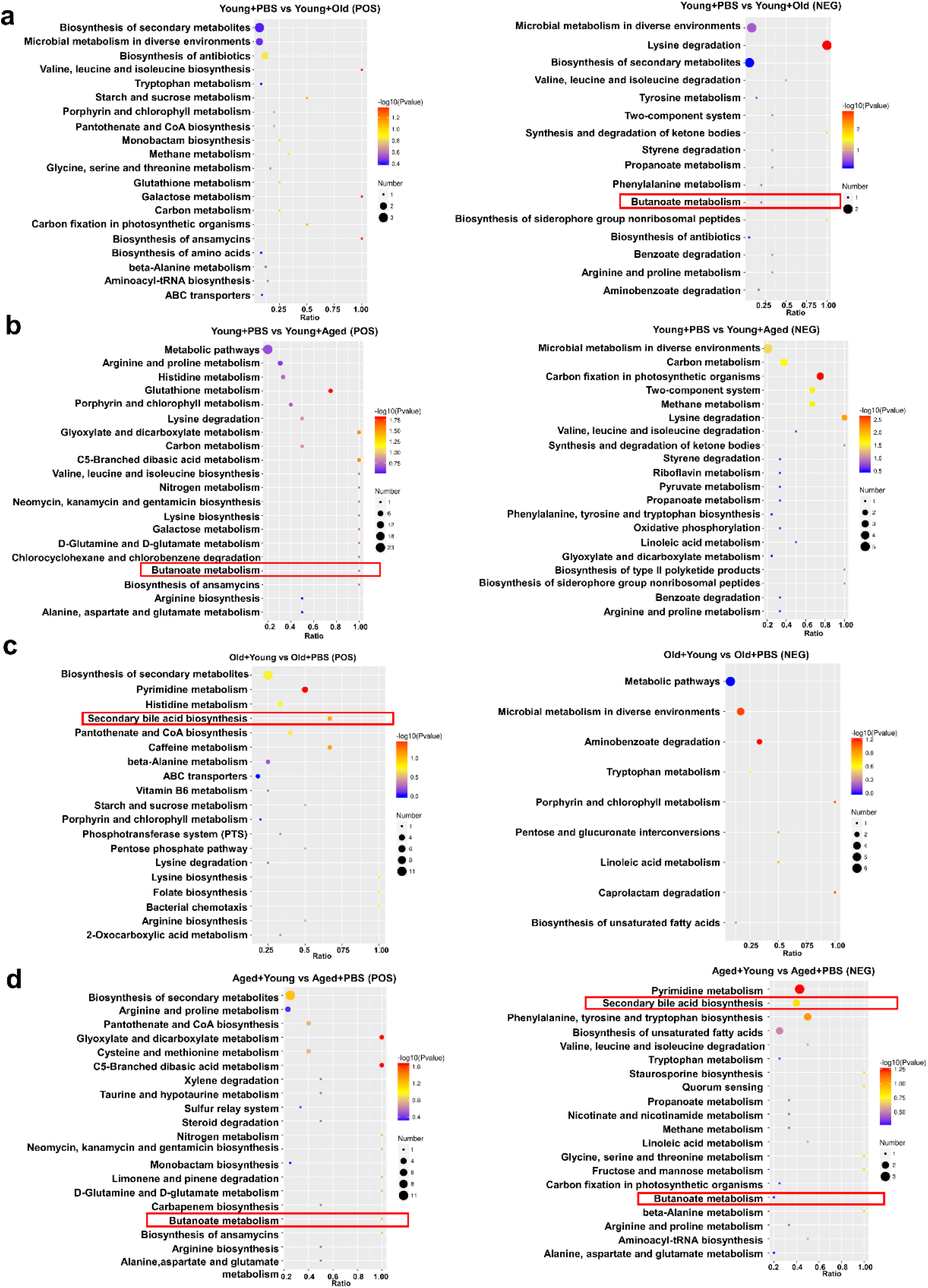
Changes in fecal microbiota metabolism and pathways after cross-age FMT. (a) Comparison of differential KEGG enrichment bubble plots between Young+PBS and Young+Old. (b) Comparison of differential KEGG enrichment bubble plots between Young+PBS and Young+Aged. (c) Comparison of differential KEGG enrichment bubble plots between Old+PBS and Old+Young. (d) Comparison of differential KEGG enrichment bubble plots between Aged+PBS and Aged+Young. The enrichment analysis was performed at the KEGG Pathway level using a hypergeometric test, as shown in the figure below. The pathways that were significantly enriched in the differential metabolites compared to the background of all identified metabolites. Pathway enrichment analysis enables us to determine the major biochemical metabolic pathways and signaling transduction pathways that are implicated by the differential metabolites.

### Butyrate supplementation inhibits gout and MSU-induced peritonitis

Based on the predictions of the functional capabilities of microbial communities and untargeted metabolomics results, we hypothesized that butyrate might play a key role. Because butyric acid is a short-chain fatty acid, we measured the levels of SCFAs (nk/g) in the Young+PBS, Aged+PBS, and Aged+Young groups. Subsequently, we observed that the butyric acid content in the faeces of Young+PBS group was significantly higher than that in the Aged+PBS group. Simultaneously, the butyric acid content in the faeces of mice in the Aged+Young was also significantly higher than that in the Aged+PBS (Fig. 7a). 2-Methylvalerate and 3-Methylvalerate were not detected, and 4- Methylvaleric was detected in only a subset of samples, thus, these results are not presented. The difference analysis of the detected short chain fatty acids is shown in Supplementary figure 7a-h. Therefore, mice were supplemented with butyrate for 14 days and then subjected to intraperitoneal injections of MSU into the dorsal aspect of the hind paws to observe its anti-inflammatory effects in acute gout. The experimental results showed that butyrate exerted significant anti-inflammatory effects in the acute gout model with inflammation induced by intraperitoneal injection of MSU. Mice with gout model, supplemented with butyrate showed significant reductions in the foot thickness ratio (Fig. 7b) and the levels of inflammatory factors (IL-1β, IL-6, and TNF-α) in their foot tissue (Fig. 7c-e). Pathological sections revealed that the foot tissue of mice supplemented with butyrate exhibited less inflammatory cell infiltration than that of the control group (Fig. 7f). Moreover, after supplementation with butyrate, mice that received an intraperitoneal injection of MSU showed significant decreases in the levels of inflammatory factors (IL-1β, IL-6, and TNF-α) in both serum (Supplementary figure 7i-k) and peritoneal fluid (Fig. 7g-i) compared with those in the control group. Subsequently, we also analyzed the expression of proteins associated with the NLRP3 inflammasome pathway. Although no significant change in the expression of NLRP3 after supplementation with butyrate was observed in both the gout and peritonitis models, supplementation with butyrate effectively suppressed the production of Caspase-1 and IL-1β (Fig. 7j and supplementary figure 7l). The anti-inflammatory effects of butyrate have been widely reported, but limited studies have investigated its specific on gout. Here we demonstrate that butyrate effectively inhibits inflammation induced by MSU stimulation.

**Fig 7.**
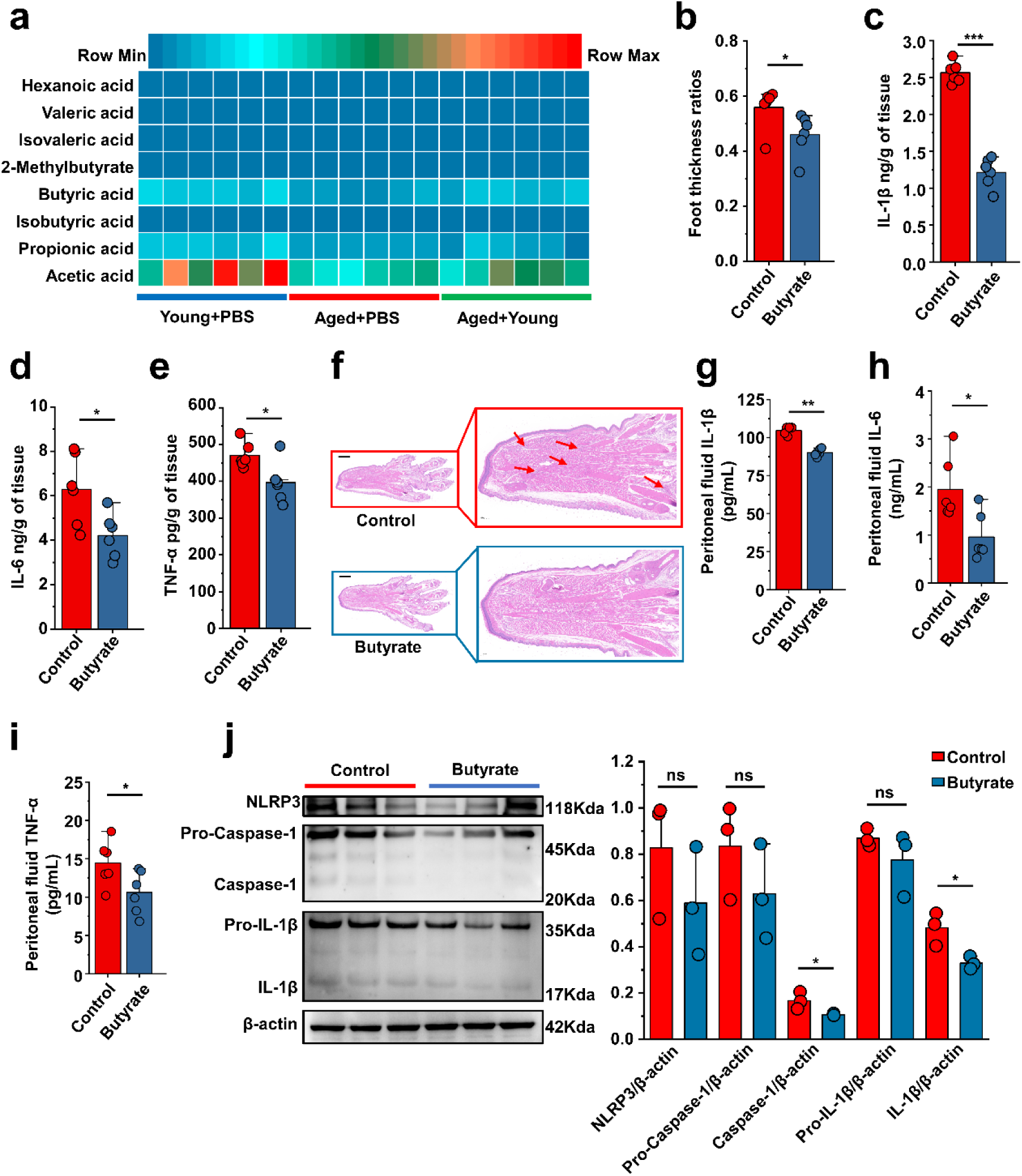
Butyrate supplementation inhibits gout and MSU-induced peritonitis. (a) The concentration of primary short-chain fatty acids (SCFAs) in fecal samples from both young (3 months), aged (24 months) mice and Aged+Young (FMT from young to aged). Graphs were generated to illustrate the changes in individual SCFAs, with a sample size of 6 in the Young+PBS, Aged+PBS and Aged+Young group(n=6). (b-e) The foot thickness ratios (b) were tested after MSU administration, foot tissue inflammatory parameters, including IL-1β (c), IL-6 (d) and TNF-α (e) concentrations, from the indicated mice are shown(n=6). (f) Representative H&E-stained images of left foot tissues. Scale bars 1000 μm and 3x magnification. (g-i) The peritoneal fluid concentrations of IL-1β (g), IL-6 (h) and TNF-α (i) inflammatory parameters were measured in the indicated mice (n=6). (j) Representative western blot images of foot tissue NLRP3 pathways proteins and band density (n=3). Values are presented as the mean ± SEM. Differences were assessed by t-test or One-Way ANOVA and denoted as follows: *p < 0.05, **p < 0.01, and ***p < 0.001, “ns” indicates no significant difference between groups.

### Serum uric acid-lowering effect of butyrate in old or aged mice

This has been proved in figure 1d, that the serum uric acid levels in mice of different age groups exhibited increase in their serum uric acid levels with increase age. There is scarce research on the impact of butyrate on serum uric acid levels, and the effects of butyrate on hepatic uric acid production or renal uric acid excretion have not been investigated. Additionally, the impact of butyrate on the serum uric acid levels in aged mice remains unexplored. Aged mice were supplemented with butyrate for 14 days, and we found that butyrate supplementation significantly reduced the serum uric acid levels in 18-month-old and 24-month-old mice (Fig. 8a). In this regard, we also assessed the activity of enzymes implicated in uric acid synthesis, namely, ADA, GDA, and XOD in the liver, as well as that of XOD in the kidney. In addition to identifying that butyrate can reduce the activity of ADA in 24-month-old mice (Aged) (Fig. 8b), and supplementation with butyrate did not have significant effects on the activities of enzymes involved in uric acid production in both 18-month-old and 24-month-old mice (Fig. 8c-e). Butyrate may not necessarily lower the serum uric acid levels by inhibiting uric acid production. Because butyrate might exert its influence on serum uric acid levels by affecting uric acid transport, we subsequently examined the expression of relevant transporter mRNA in mouse kidneys and the expression of the colonic ABCG2 transporter that facilitates uric acid excretion. The expression of the renal injury marker KIM-1 was significantly decreased in mice supplemented with butyrate (Fig. 8f). In contrast to the results obtained cross-age FMT, no significant difference in the expression of URAT1 was observed (Fig. 8g). However, the expression of GLUT9 was significantly decreased in 24-month-old mice after butyrate supplementation, whereas no significant difference was observed in the 18- month-old mice (Fig. 8h). We also found that the supplementation of old or aged mice with butyrate resulted in an increased mRNA expression level of uric acid transporters involved in uric acid excretion and significantly increased the expression of OAT1 and OAT3 (Fig. 8i-j). Although no significant difference in the expression of ABCG2 was detected in old mice, a significant increased in its expression was observed in aged mice after supplementation with butyrate (Fig. 8k). Meanwhile, the mRNA expression of ABCG2 in the colons was significantly increased in old or aged mice after supplementation with butyrate (Fig. 8l). Moreover, supplementation with butyrate significantly increased the mRNA expression of ZO-1, Occludin, and JAMA in the colons of mice (Supplementary figure 8a-c). Based on the abovementioned results, we found that supplementation with butyrate promotes uric acid excretion and improves the intestinal tight junction integrity.

**Fig 8.**
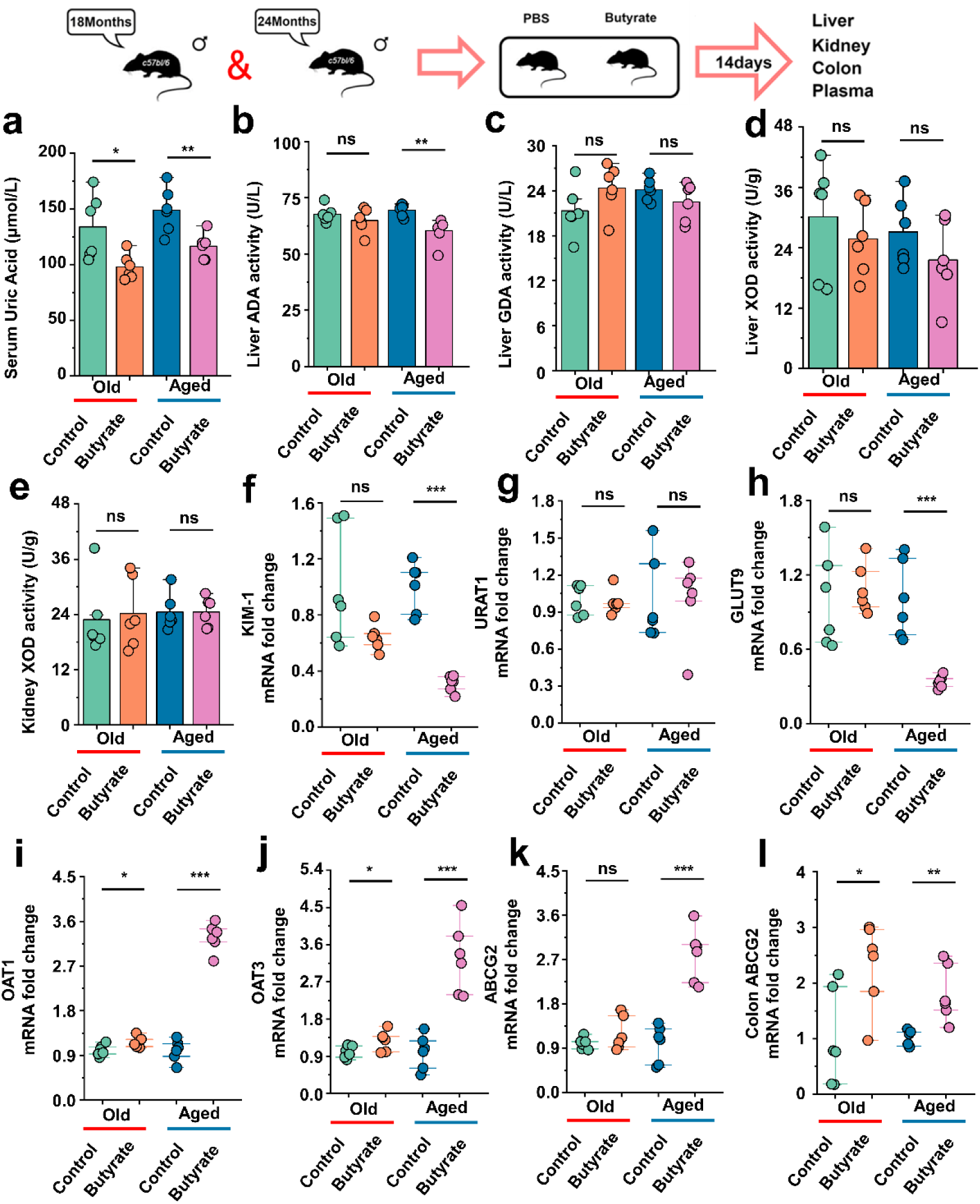
Serum uric acid-lowering effect of butyrate in old or aged mice. (a) The serum uric acid concentrations in Old+PBS and Old+Butyrate, Aged+PBS and Aged+Butyrate (n=6). (b-d) The activity of uric acid-producing enzymes of liver in the Old or Aged control group and the old or aged group supplemented with butyrate (n=6), including ADA (b), GDA (c) and XOD (d). (e) The activity of XOD of kidney in the Old or Aged control group and the old or aged group supplemented with butyrate (n=6). (f) Relative kidney injury molecule-1 (KIM-1) expression in the indicated groups by qPCR (n = 6). (g and h) Relative renal genes for uric acid reabsorption expression in the indicated groups by qPCR (n = 6), including URAT1 (g) and GLUT9 (h). (i-k) Relative renal genes for uric acid excretion expression in the indicated groups by qPCR (n = 6), including OAT1 (i), OAT3 (j) and ABCG2 (k). (l) Relative colonic genes for uric acid expression in the indicated groups by qPCR, including ABCG2 (n = 6). Values are presented as the mean ± SEM. Differences were assessed by t-test or One-Way ANOVA and denoted as follows: *p < 0.05, **p < 0.01, and ***p < 0.001, “ns” indicates no significant difference between groups.

## Discussion

The incidence of gout and hyperuricemia continually to increases as individuals. However, research on gout among the elderly population is relatively limited. Previous studies have confirmed a close relationship between gout and the gut microbiota(24, 25), but the specific connection between the gut microbiota of older individuals and gout remains unexplored. FMT from young to old mice also reportedly alleviates age-related stroke(26)and other benefits(11, 27). And there are also studies indicating Older patients demonstrate heightened inflammatory reactions during gout attacks(28). Based on this finding, we hypothesize that the gut microbiota of different age groups may exert varying effects on gout. Therefore, we conducted cross-age FMT to investigate the interaction between the gut microbiota in different age groups and gout.

This study demonstrated that FMT from young to aged mice effectively alleviated the inflammatory response caused by MSU and improved uric acid metabolism in elderly mice, reducing the symptoms of gout. In contrast, FMT from old or aged to young mice exacerbated gout, indicating the importance of the gut microbiota composition in gout development. Activation of the NLRP3 inflammasome has been implicated in the pathogenesis of gout, leading to the production of inflammatory cytokines such as IL-1β, IL-6, and TNF-α. The beneficial effects of FMT from young to aged mice could be attributed to inhibition of the NLRP3 inflammasome pathway. A “younger” gut microbiota could suppress the activation of NLRP3, Caspase-1 and IL-1β, thereby reduce the inflammatory response in gout. Furthermore, the modulation of uric acid metabolism in old or aged mice played a role in the effects of young gut microbiota transplantation. Additionally, the improvement in the intestinal tight junction integrity observed after transplantation of the gut microbiota from young to old or aged mice may contribute to enhanced elimination of uric acid. Recent studies have also shown a direct relationship between hyperuricemia and gut microbiota. Certain anaerobic microbial communities are capable of effectively degrading purines and uric acid and thereby regulate the abundance of purines in the body to maintain the body’s uric acid balance(29–31). Our study also showed that the gut microbiota plays a role in maintaining the balance of serum uric acid in the body. The “ageing” gut microbiota is more likely to elevate the levels of serum uric acid in the body by suppressing the expression of uric acid excretion transporter (OAT1, OAT3). The “younger” gut microbiota helps to restrain the increase of serum uric acid levels in old or aged mice by suppressing the expression of uric acid reabsorption transporters (URAT1) and the activity of uric acid-producing enzymes (mainly XOD) and also promotes the expression of uric acid excretion transporters (ABCG2) in aged mice. We also found that old or aged mice exhibited stronger purine metabolism (Table 2a, b). These results may explain the reason for the high levels of uric acid detected in elderly individuals.

Moreover, through 16S rDNA sequencing analysis, we found that the abundance of Bifidobacterium and Akkermansia was higher in young mice. Additionally, FMT from young to old or aged mice increased the abundance of the Akkermansia genus. There are existing reports regarding the therapeutic effects of Bifidobacterium(32) and Akkermansia(33) on gout and hyperuricemia. Therefore, we performed further investigation, and using microbiome-predicted functional and untargeted metabolomics data, we observed a significant enhancement in butanoate metabolism in young mice. Surprisingly, using untargeted metabolomics data, comparisons of the results after FMT from young to old or aged mice with those of old or aged mice also identified the secondary bile acid biosynthesis pathways. Studies have indicated that transplantation of the gut microbiota from healthy mice to progeroid mice can enhance the enrichment of secondary bile acid biosynthesis pathways, and the restoration of secondary bile acids may potentially contribute to extension of the healthspan and lifespan in progeroid mice(34). The current investigation into the impact of secondary bile acids on gout and hyperuricemia is still in its incipient phase, with the intrinsic mechanisms remaining largely unelucidated, thereby rendering this domain a pivotal subject of profound scientific inquiry.

Moreover, old or aged mice transplanted with gut microbiota from young mice also exhibited an increase in butanoate metabolism. Upon comparison, we discovered other bacteria genera that produce butyrate, such as Lachnoclostridium. Additionally, literature(35, 36) reports have indicated that Bifidobacteria combined with other genera can enhance the production of butyrate. Meanwhile, Akkermansia, particularly the species Akkermansia muciniphila, has been found to confer several beneficial traits, as evidenced by preclinical studies. These traits include promoting the growth of butyrate-producing bacteria through the production of acetate, which leads to a decrease in the loss of the colonic bilayer and subsequent reduction in inflammation(37). These findings led us to hypothesize that butyric acid might play a role in this phenomenon. Although, Figure 7 does show a similar trend for acetic and propionic acids as for butyric acid. However, considering the predictive data of microbial function and the non-targeted metabolomic data, there is an enhancement of Butanoate_metabolism in both young mice and elderly mice receiving young mouse intestinal microbiota transplants. Therefore, we prioritized butyrate as the subject of our study. Due to the scarcity of elderly mice, we are unable to conduct subsequent experiments with acetic and propionic acids, which is one of the limitations of this study. This work will be addressed in our follow-up research. Subsequently, we selected the Young+PBS, Aged+PBS, and Aged+Young groups for analysis of the short-chain fatty acid levels and discovered that the fecal butyric acid content in young mice was higher than that in aged mice, and that FMT from young mice led to an increase in fecal butyric acid content in aged mice. Consistent with previous reports, the content of SCFAs in the faeces of elderly mice was lower than that in the faeces of young mice(38). This result shifted our focus to butyric acid. Butyric acid, a SCFA derived from the gut microbiota, has been found to play a crucial role in host health(39). Emerging evidence indicates that among SCFAs, butyrate plays a pivotal role as a regulator in mediating metabolic control of the microbiota(40). The anti-inflammatory effects of butyric acid have been extensively reported,(41, 42) but only a few studies have investigated its impact on gout. Similarly, research on the anti-hyperuricemia effects of butyric acid is relatively scarce. We then conducted experiments using gout and peritonitis models, and found that mice supplemented with butyrate exhibited relief in symptoms of gout and inflammation. Butyrate could also reduce the serum uric acid levels in old or aged mice by promoting the expression of uric acid excretion transporters.

The findings of this study demonstrate the positive effects of gut microbiota transplantation from young to aged mice in mitigating gout symptoms. Inhibition of the NLRP3 inflammasome pathway was found to modulate uric acid metabolism, and increase butyric acid levels. The gut microbiota of young mice effectively alleviated the inflammatory response and improved uric acid metabolism in aged mice. We enhanced Butanoate metabolism in the gut microbiota of young mice, while the opposite was observed in aged mice. Furthermore, we observed higher levels of butyrate in the feces of young mice. Moreover, butyrate has excellent anti-inflammatory and uric acid-lowering effects. These findings highlight the potential of targeting the gut microbiota as a therapeutic approach for the prevention and treatment of senile gout-related conditions.

## Materials and Methods

### Preparation of MSU crystals

A solution consisting of 10 mM uric acid and 154 mM NaCl (both from Sigma) was adjusted to pH 7.2 and agitated for 7 days to produce MSU crystals. For sterilization the needle-shaped crystals were washed with ethanol, dried under sterile conditions and heated at 180°C for 2 h. The crystals were stored at 20 mg/mL in sterile PBS for the experimental model.

### Animals

All the mice in the different age groups were purchased from Longde Biotechnology Co., LTD (Changchun, Jilin, China). The mice were housed in a controlled environment with a temperature of 24±2°C and a 12-hour light/12- hour dark cycle, in a pathogen-free facility. These mice were provided ad libitum access to food and water. All animal experiments were conducted in compliance with the regulations set forth by the Administration of Affairs Concerning Experimental Animals in China. The Institutional Animal Care and Use Committee at Jilin University approved the entire protocol, including the mouse autopsy and sample collection (20170318).

### Animal and FMT

Young (3 months), old (18 months) and aged (24 months) male mice were randomly reassigned to experimental cages one week prior to the experiment. The mice in the FMT groups were subjected to antibiotic interventions and depletion of the host gut microbiota by via oral gavage of a 3-day broad-spectrum antibiotic cocktail regimen (vancomycin, 100 mg/kg; metronidazole, 200 mg/kg; ampicillin 200 mg/kg, neomycin 200 mg/kg) to maximize potential engraftment of the donor microbiota. Those in the PBS groups were orally gavaged with isopycnic PBS.

Before antibiotic interventions, fecal slurries from the donor mice were prepared by pooling fecal pellet material from 12-15 mice per cage (chosen randomly) for each age group. At the time of pellet collection, at least three pellets from each mouse were collected into sterile centrifuge tubes, and the mice were then returned to their cage. The fecal pellets from each age group were mixed, and sterile PBS (five times the weight of the fecal pellets) was added. The homogenate was centrifuged at 500 × g and 4°C for 10 min and fecal slurry was collected. After addition of 10% glycerol (v/v) and the slurry was stored at –80°C until use. After antibiotic washout, recipient mice were reassigned to different groups and received heterochronic donor microbiota twice 3 days apart by oral gavage of the fecal slurry preparation. Each mouse received 200 μL the fecal slurry preparation during each gavage.

Control groups for 3-month-old, 18-month-old and 24-month-old receiving heterochronic transfer were gavaged with PBS only, revealed antibiotic treatment/PBS gavage only and were administered control microbiota (obtained from a donor pool of the same age group).

Before MSU injection, fecal pellets were collected for sequencing and untargeted metabolomics analysis on day 22, and were then stored at −80°C. Blood and tissues were collected at the end of the experimental period. All FMT interventions were performed at the same time of day across all the cages to control for the circadian rhythm variabilities in feeding and the microbiota/metabolite composition.

### Experimental model details

#### Gout model

To establish this model, we administered 0.4 mg of preformed MSU crystals, resuspended in 20 μL of sterile PBS via subcutaneous injection into the left hind paws of the animals. We measured the hind paw thickness using calipers (the minimum accuracy of which was 0.01 mm) before and 8 hours after MSU injection. Eight hours after injection, mouse dorsal foot tissues were collected for histopathological observation and protein extraction.

#### Peritonitis model

Peritonitis was induced in the mice by intraperitoneal injection of MSU crystals at a dose of 2.5 mg in PBS. Six hours later, the peritoneal cells were collected by lavage (3 mL of PBS), and the cell count was determined using a hemocytometer to ensure consistent cell numbers. The lavage fluid was used to detect cytokines, and proteins were cells extracted from the cells.

#### Butyrate supplementation experiment

In this experiment, 3-month-old mice received a supplemental dose of butyrate (administered via gavage at a concentration of 200 mg/kg) for 14 days. Subsequently, these mice were subjected to monosodium urate (MSU) stimulation. The investigation was conducted utilizing both gout and peritonitis models to assess the effects of butyrate supplementation. Simultaneously, mice aged 18 months and 24 months were given the same dosage of supplemental butyrate as the 3-month-old mice for 14 days. Subsequently, their liver, kidneys, intestines, and blood samples were collected for analysis.

### Histological analysis

The left hind paws were collected and immediately fixed with 4% paraformaldehyde for more than 48 h. The fixed tissues were prepared after paraffin embedding. At least three slices from each tissue were prepared. The prepared sections were then dewaxed with xylene and hydrated via different gradients of alcohol. Furthermore, all hydrated sections were stained with hematoxylin &eosin (H&E), detected under an optical microscope (Olympus, Tokyo, Japan) and analyzed with Caseviewer2.0 software.

### Western blotting

The total proteins from the peritoneal cells and foot tissue samples were extracted using a protein extraction kit (Thermo Fisher Scientific, USA). The targeted proteins were separated by 10% or 15% SDS‒PAGE based on the molecular weight and then bonded to 0.22 μm PVDF membranes. After blocking with 5% skim milk for three hours at RT, specific antibodies, including NLRP3, Caspace-1, IL-1β and β-actin, were used to detected target proteins at an appropriate final concentration according to the manufacturer’s instructions. The PVDF membranes were then incubated with goat anti-rabbit IgG (1:20,000), wash with TBST, and determined using the ECL plus western blotting detection system (Tanon, China). The grey values of the western blotting bands were procured using Image-Pro Plus 6.0 software.

### RNA extraction and qPCR

Tissue samples were collected and total RNA was extracted using Trizol (Takara, CA, JPN) as previously described, cDNA was reverse transcribed using a TransScript Uni All-in-One (TransGen Biotech, Beijing, China) and reacted with specific primers using a FastStart Universal SYBR Green Master Mix (ROX) (Roche, Switzerland, Basel) and Bio-Rad CFX96 (Bio-Rad Laboratories, CA, USA). The specific primers used in this study are shown in Table 1. The 2^−ΔΔCt^ method was used to calculate the relative expression levels of genes using the control group as the calibrator.

**Table 1.**
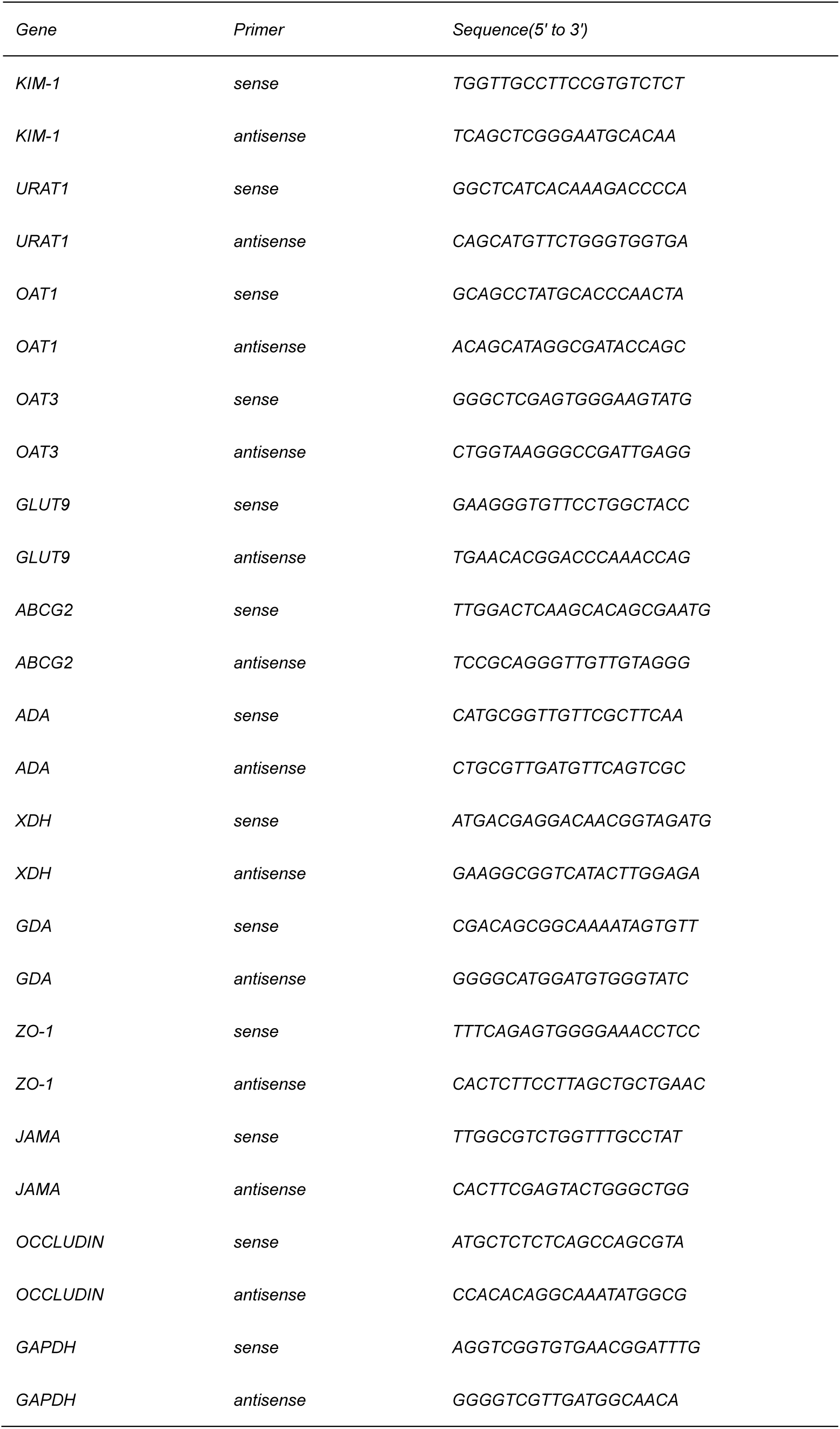
Oligonucleotides.

**Table 2.**
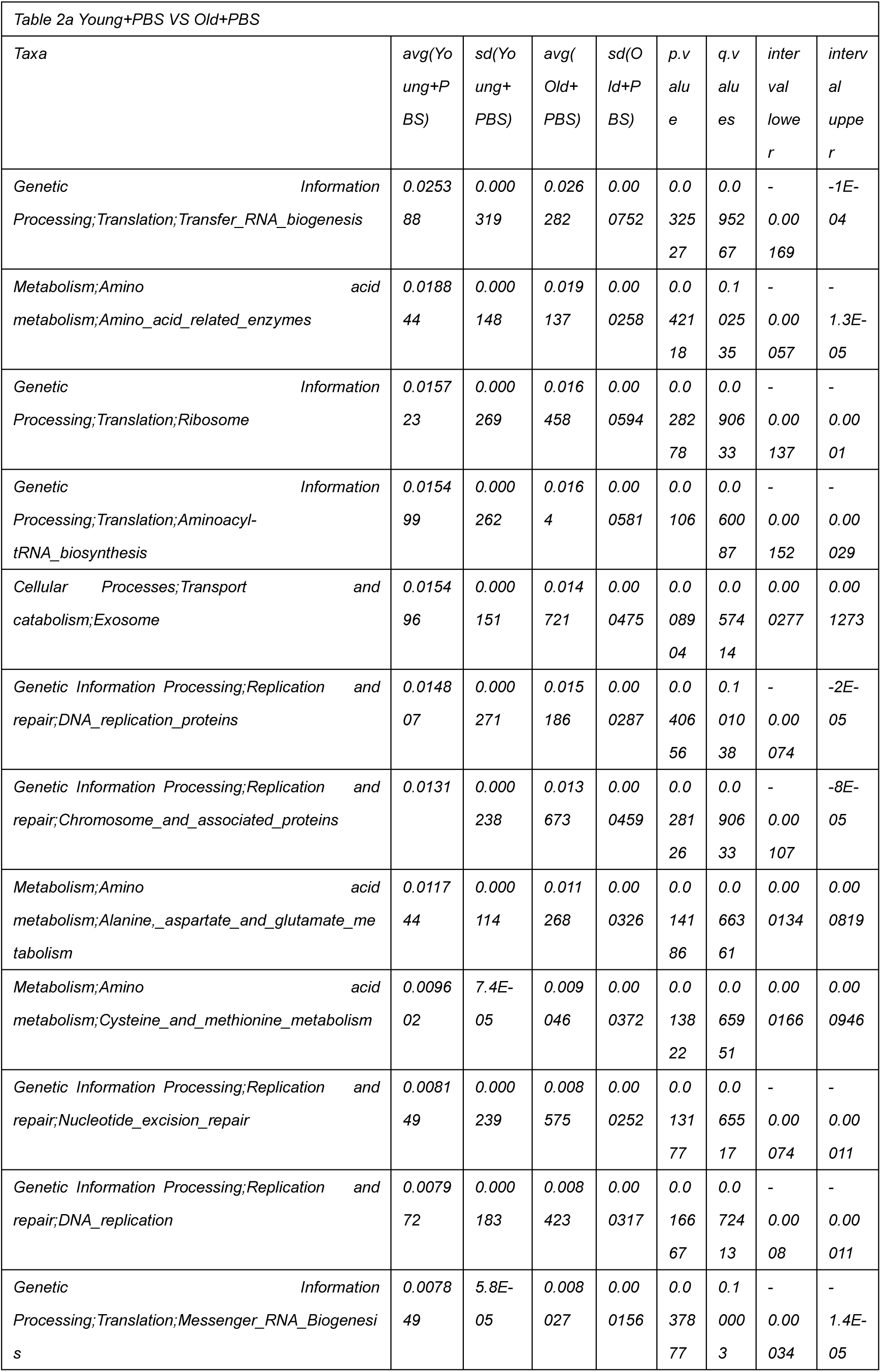
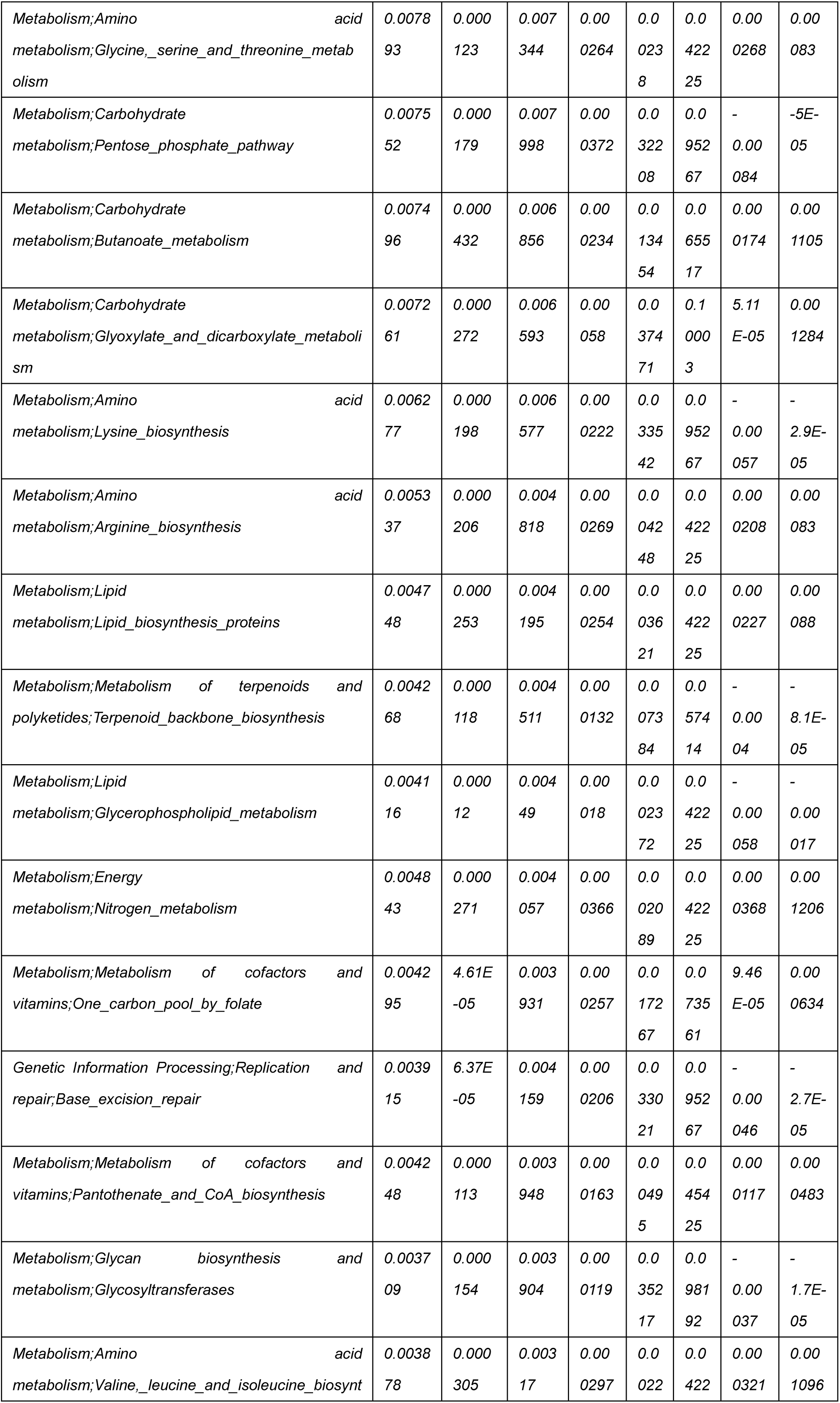
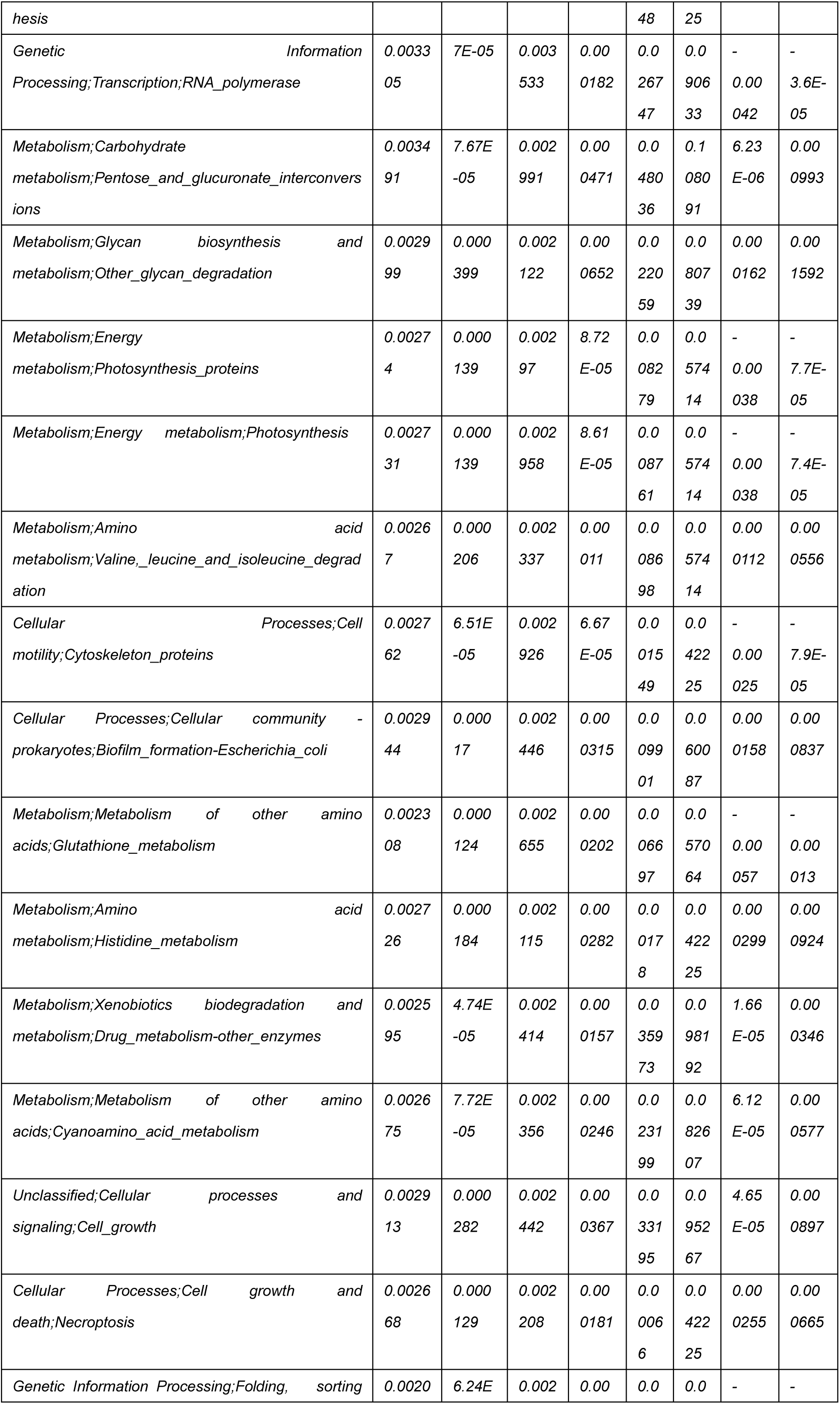
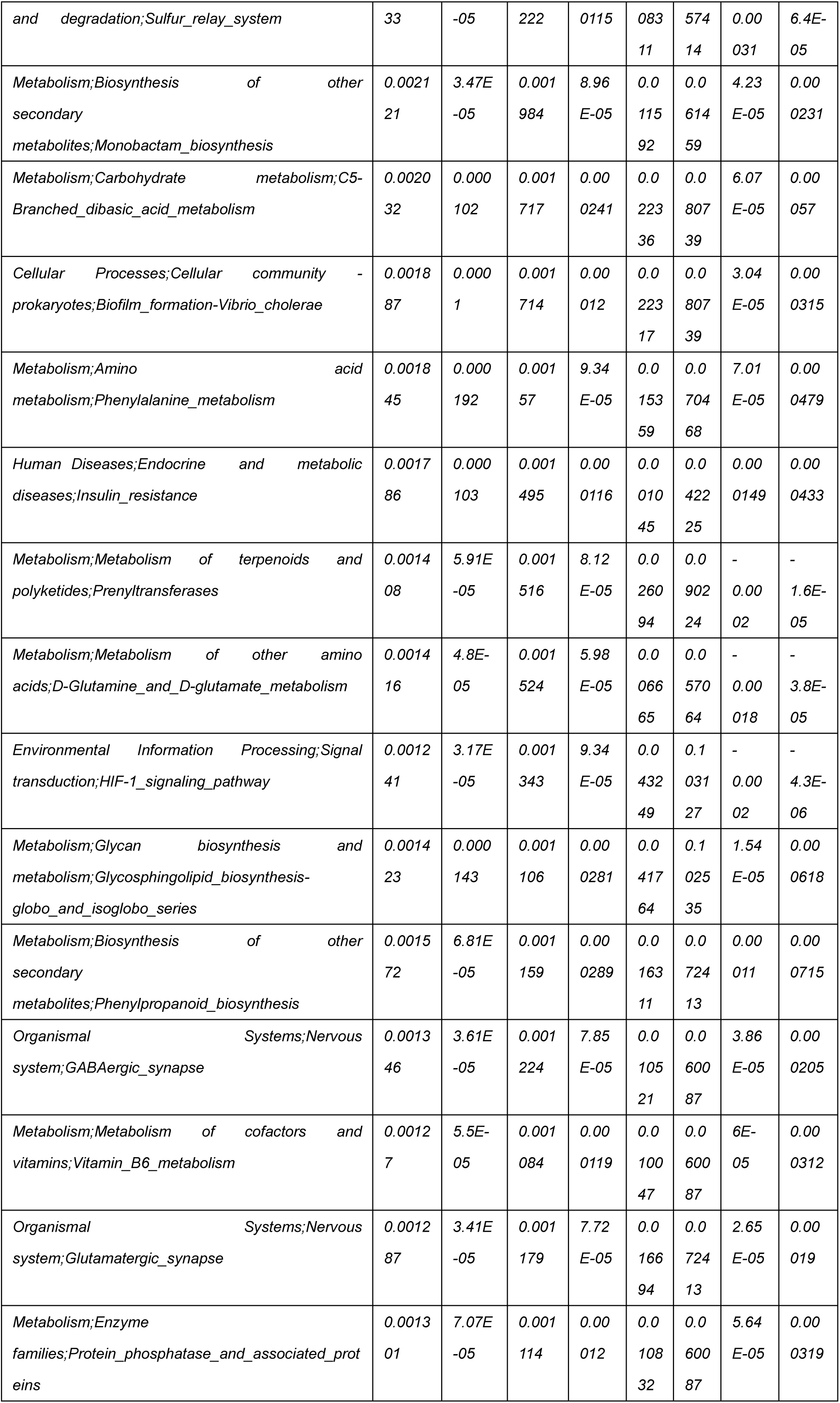
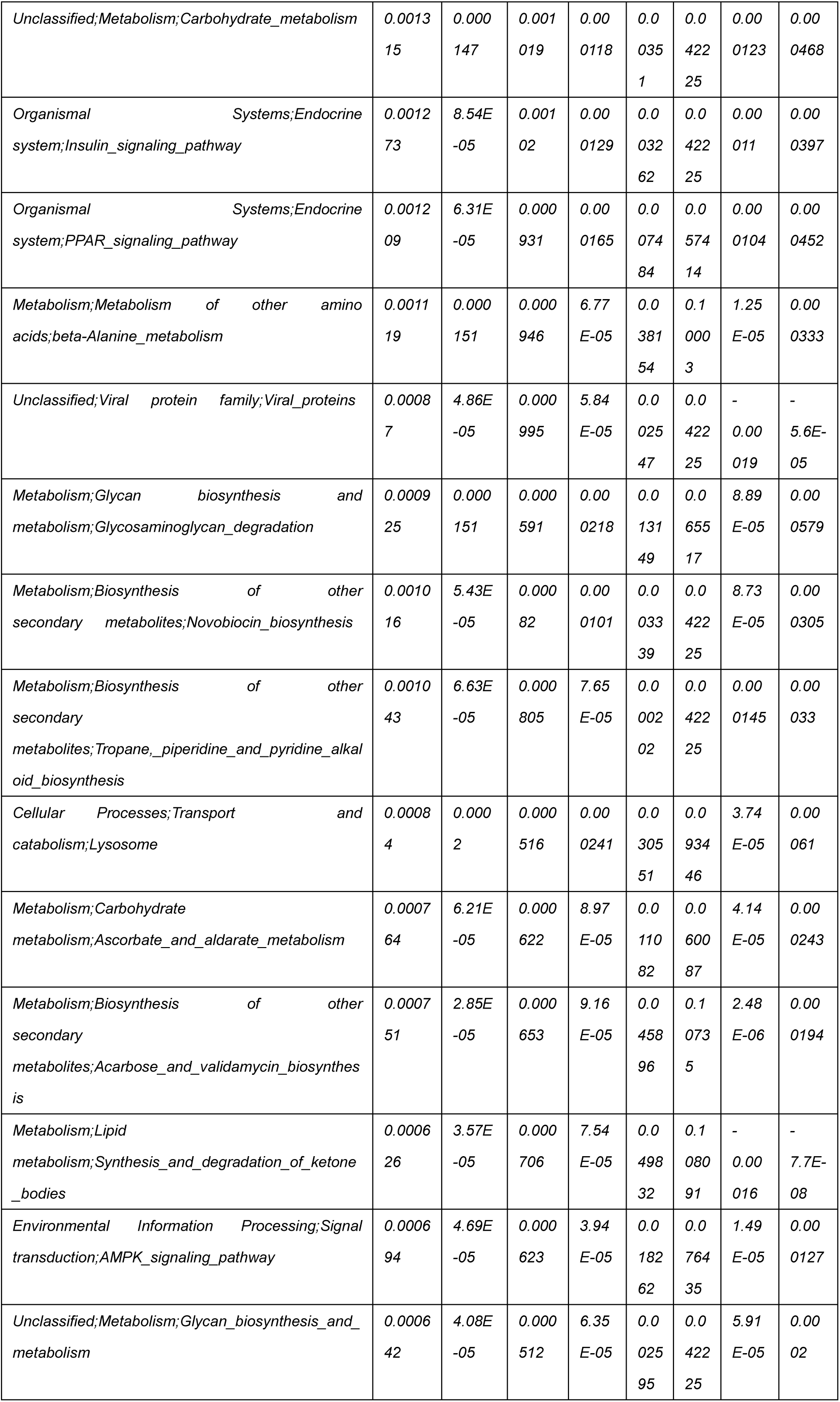
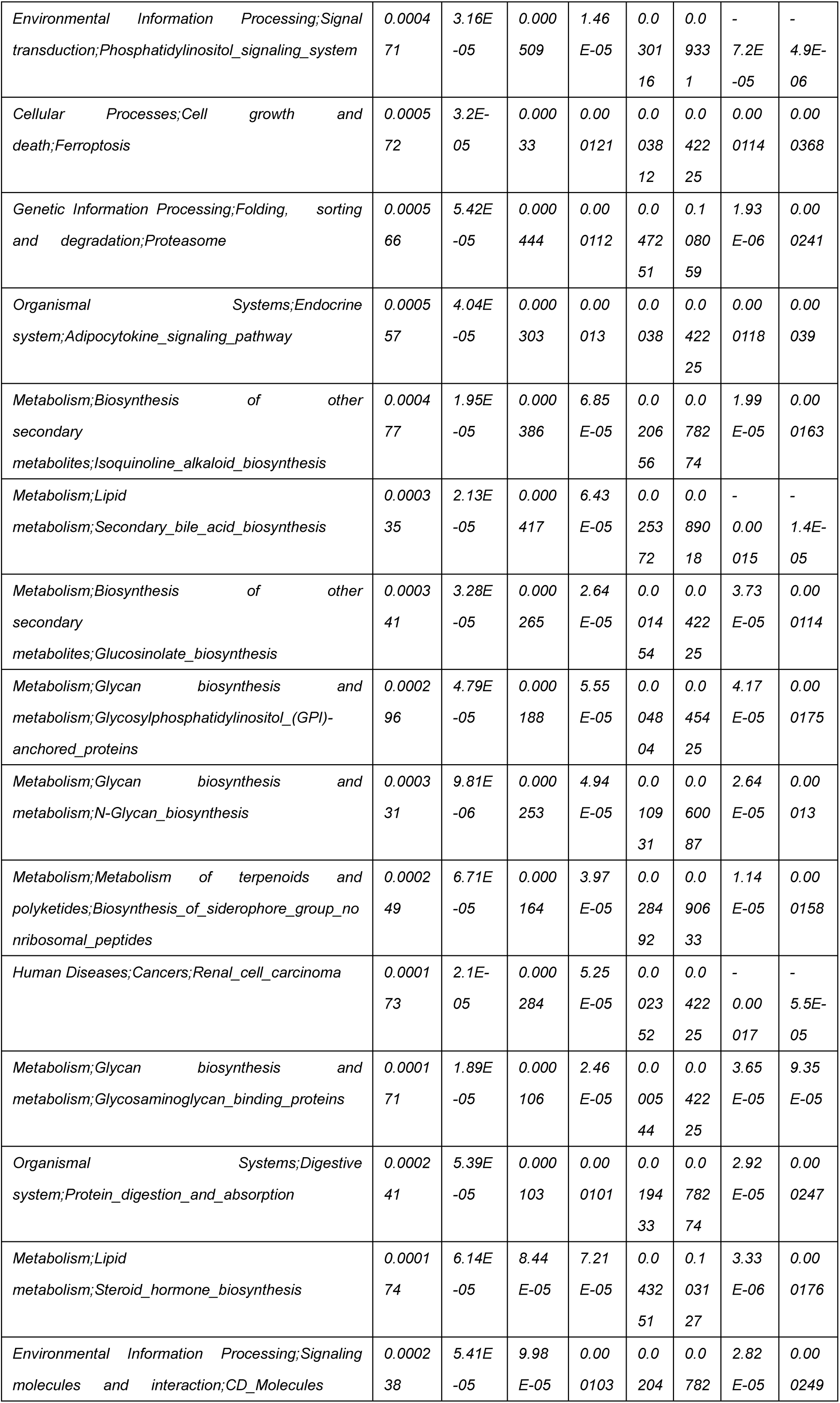
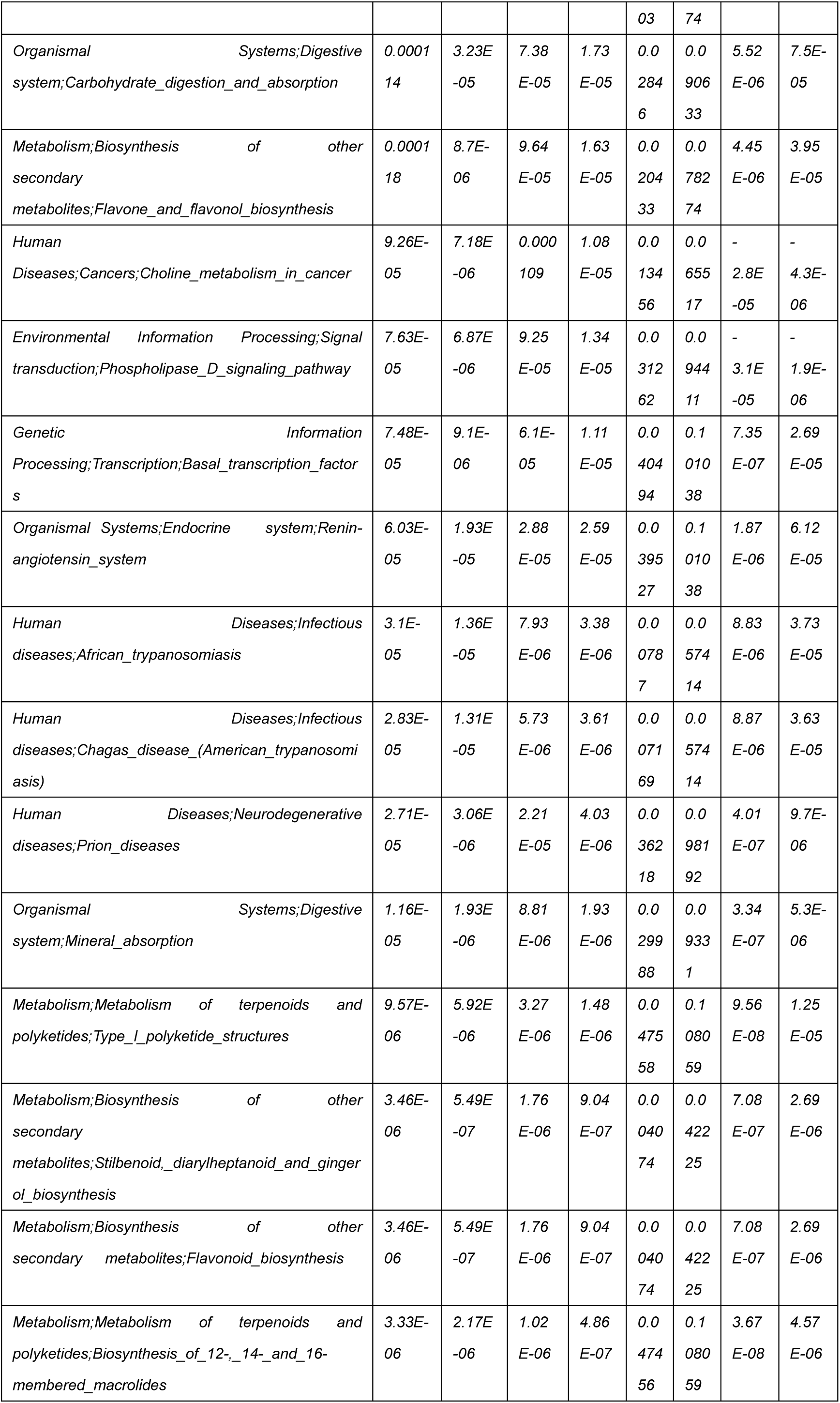
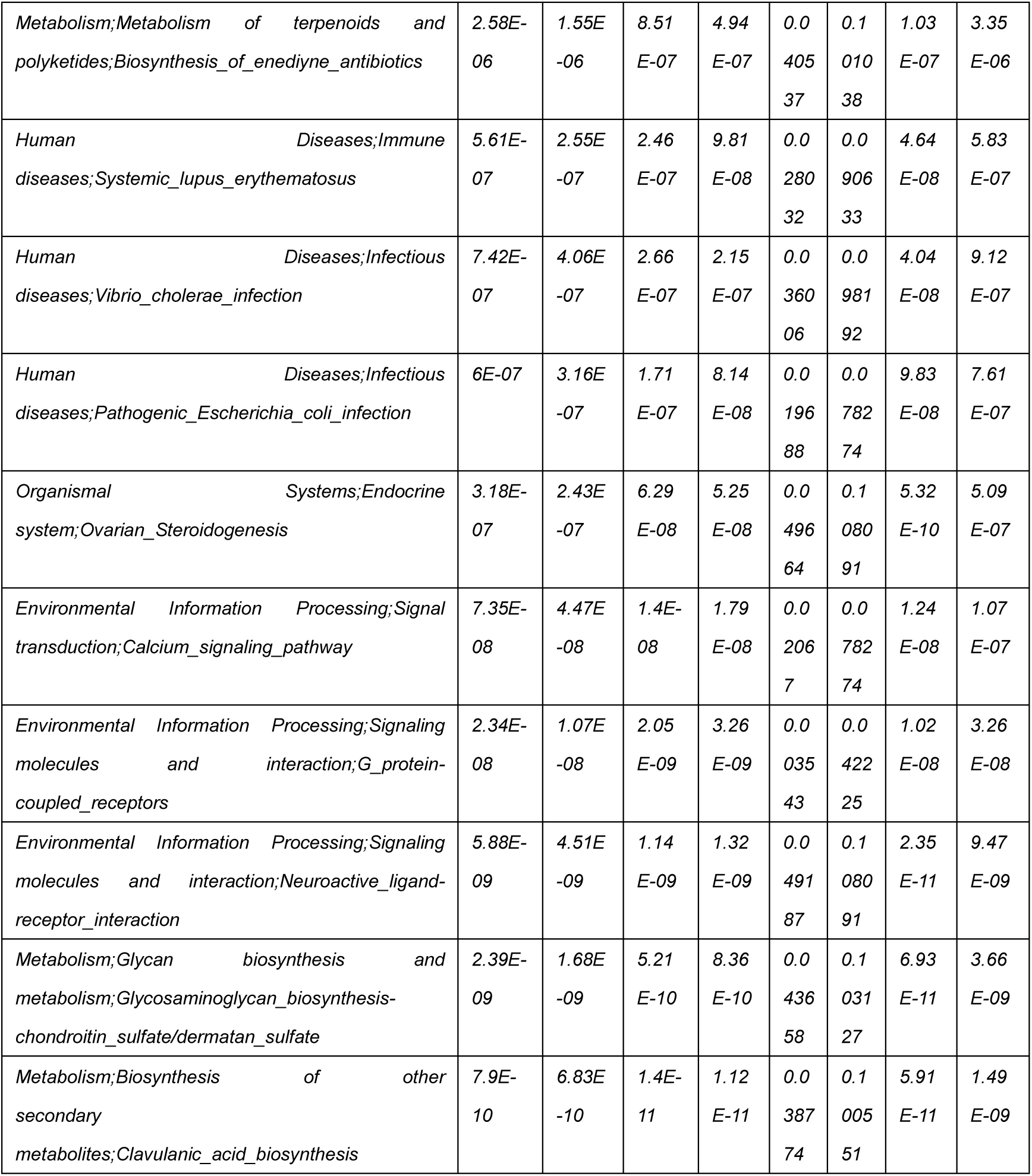
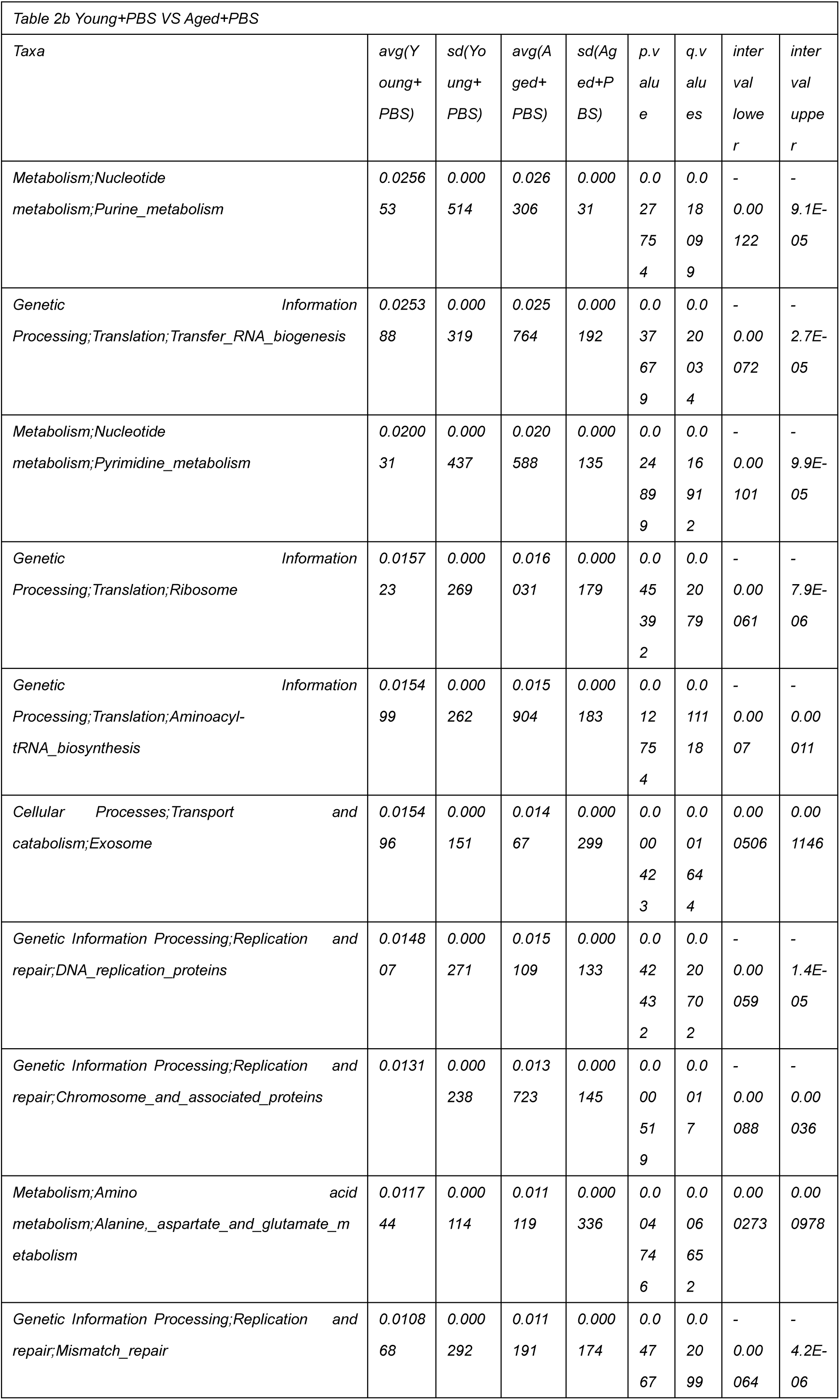
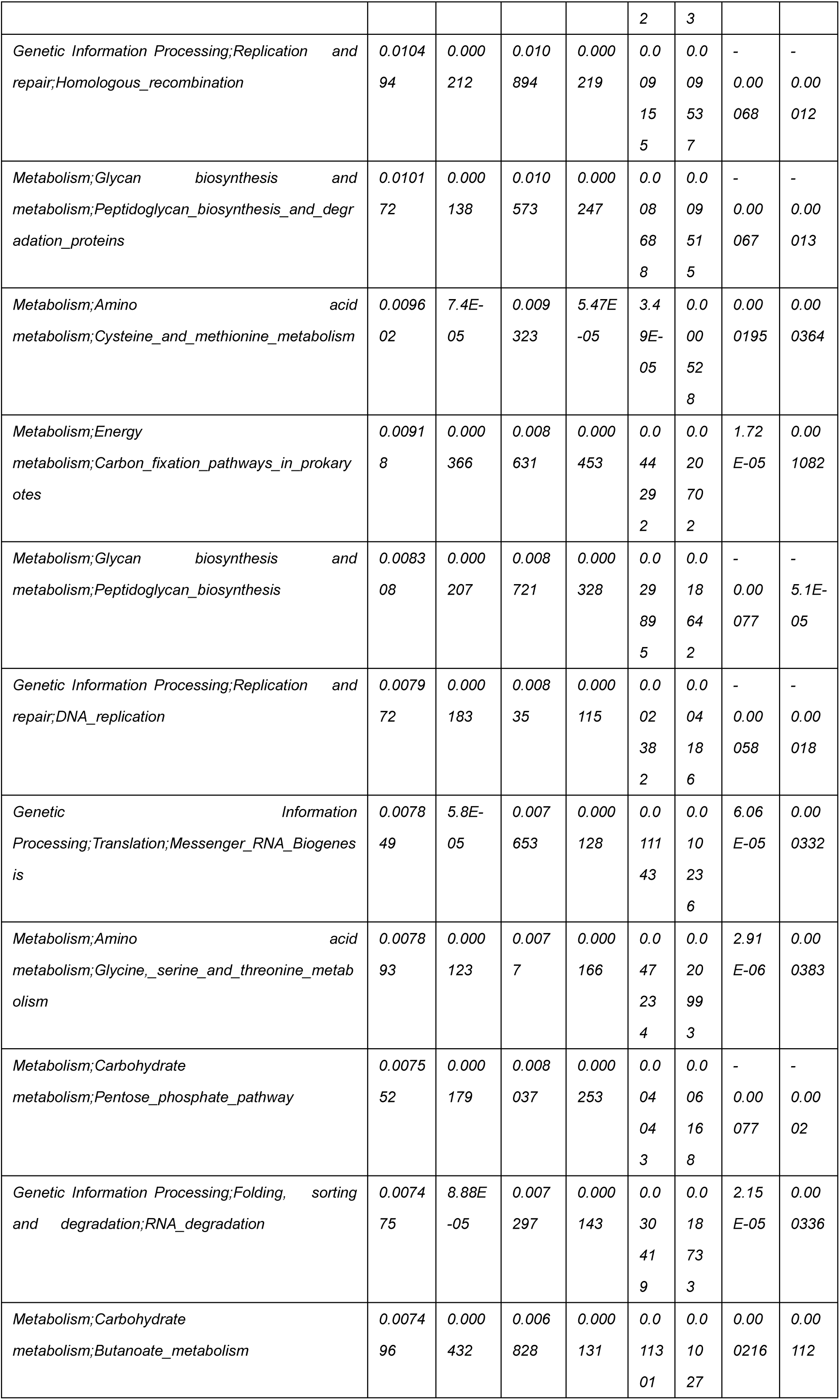
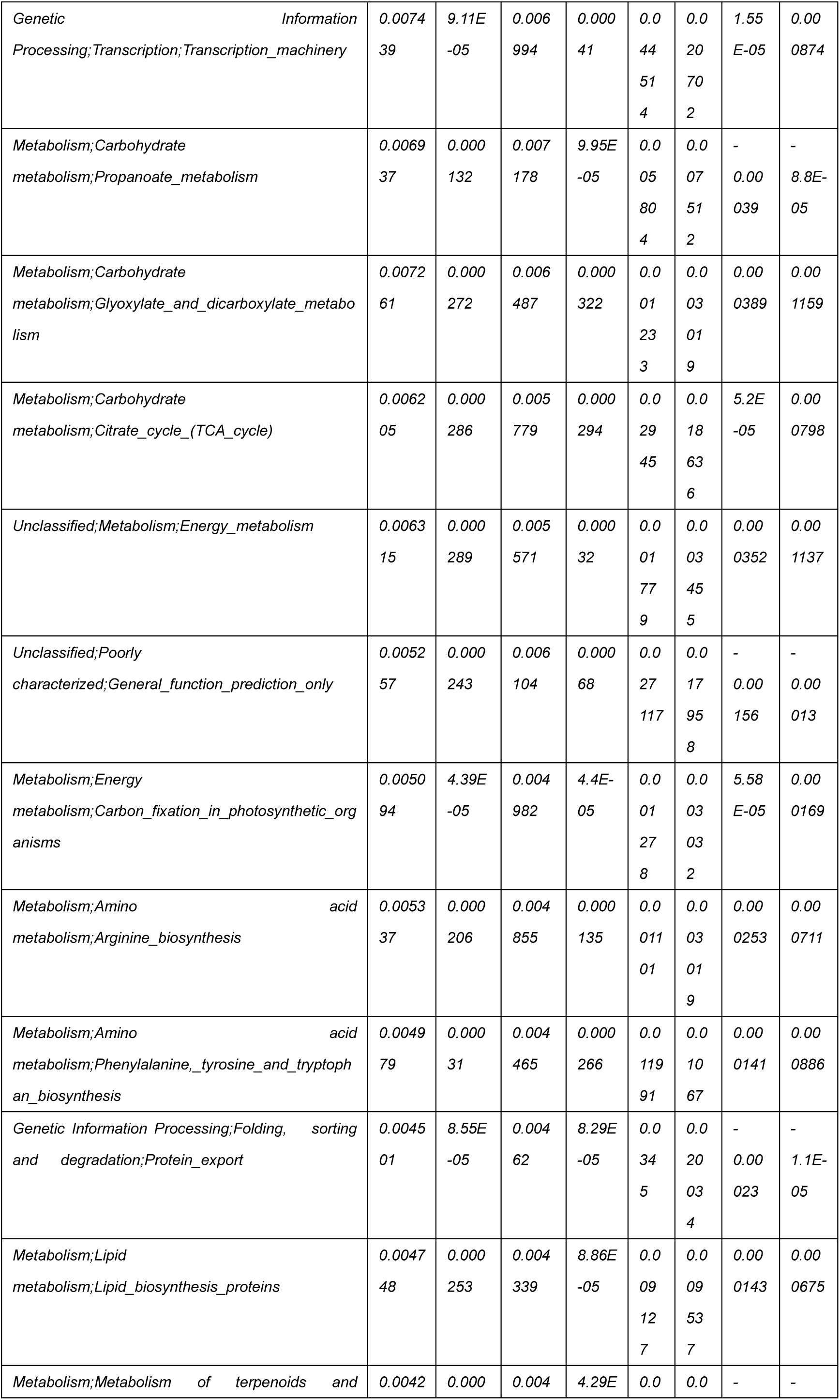
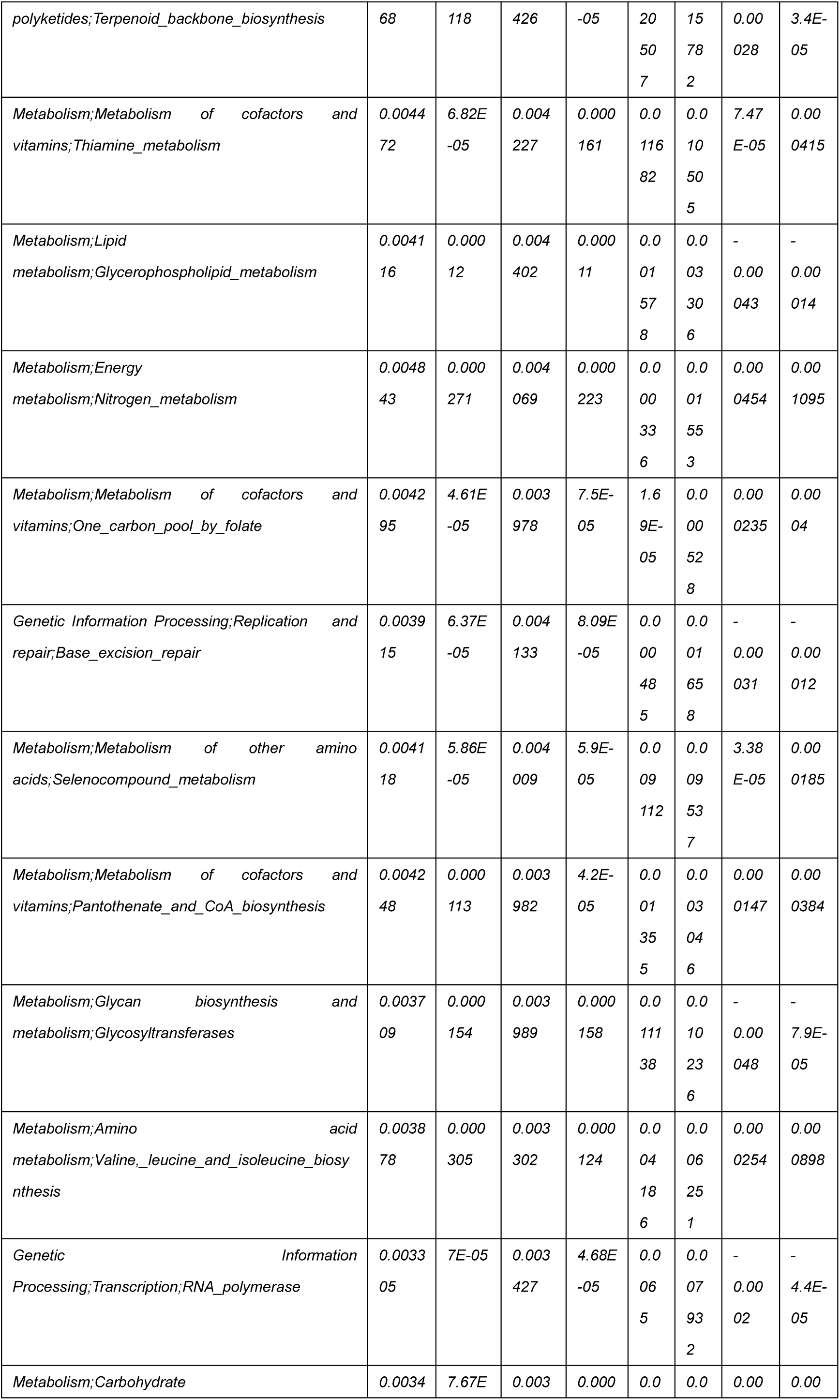
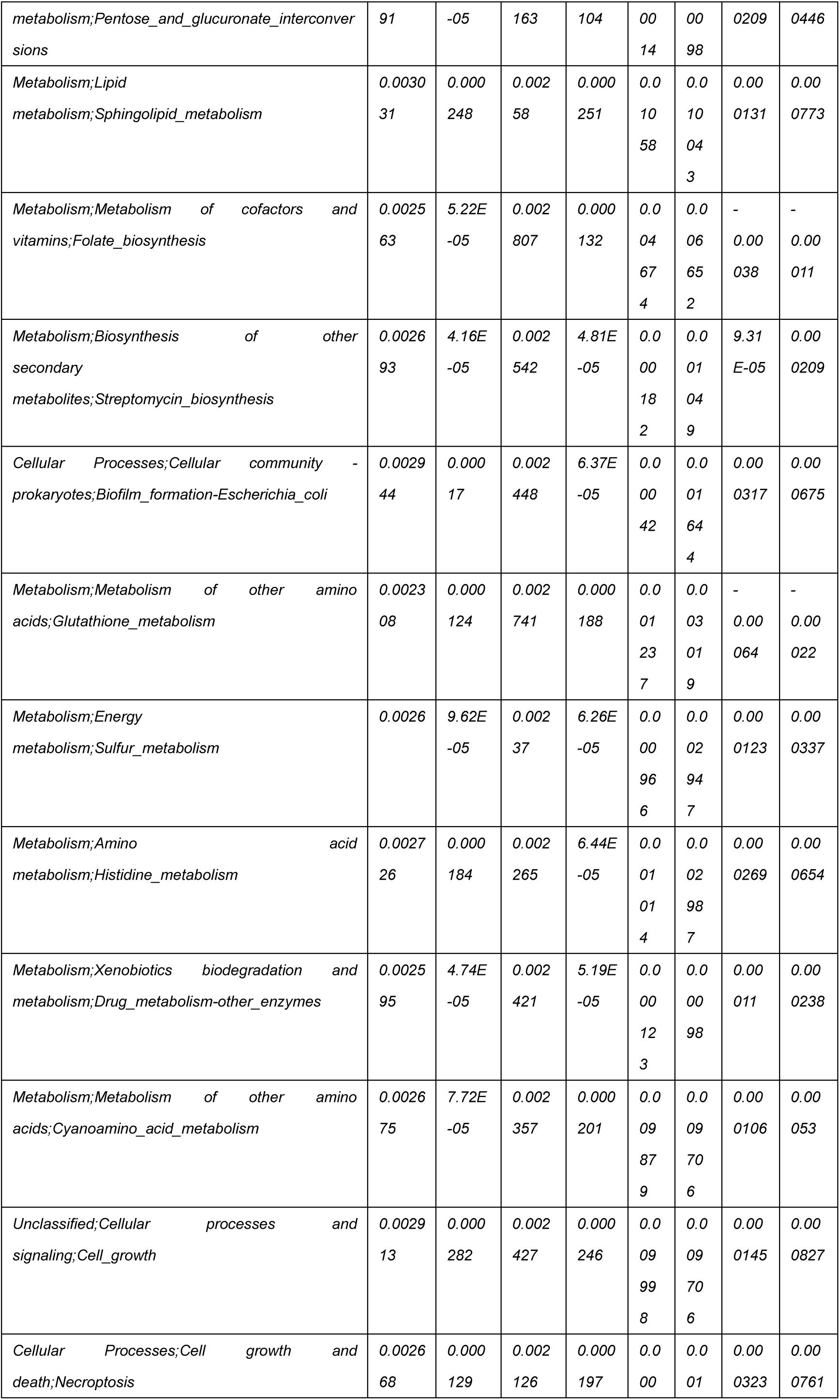
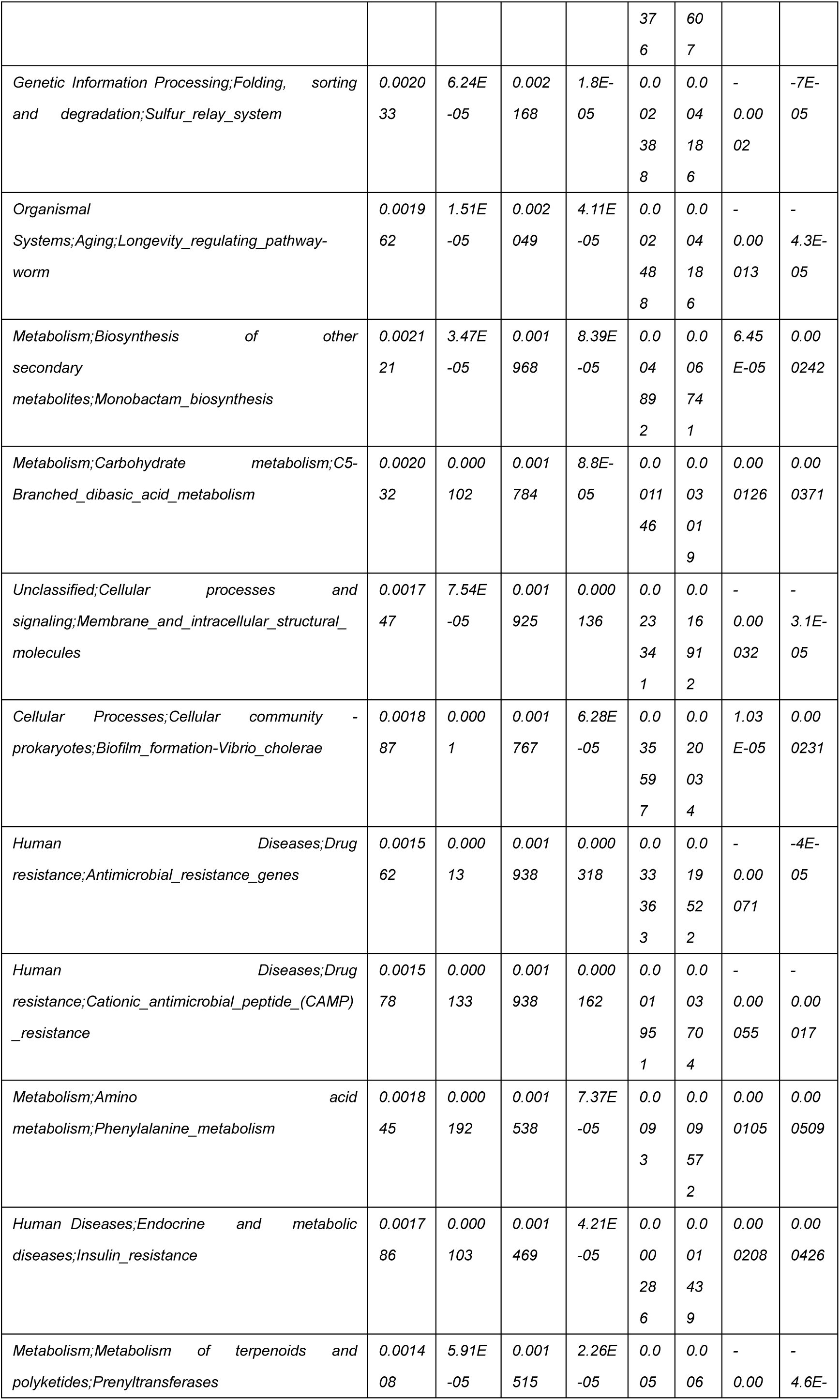
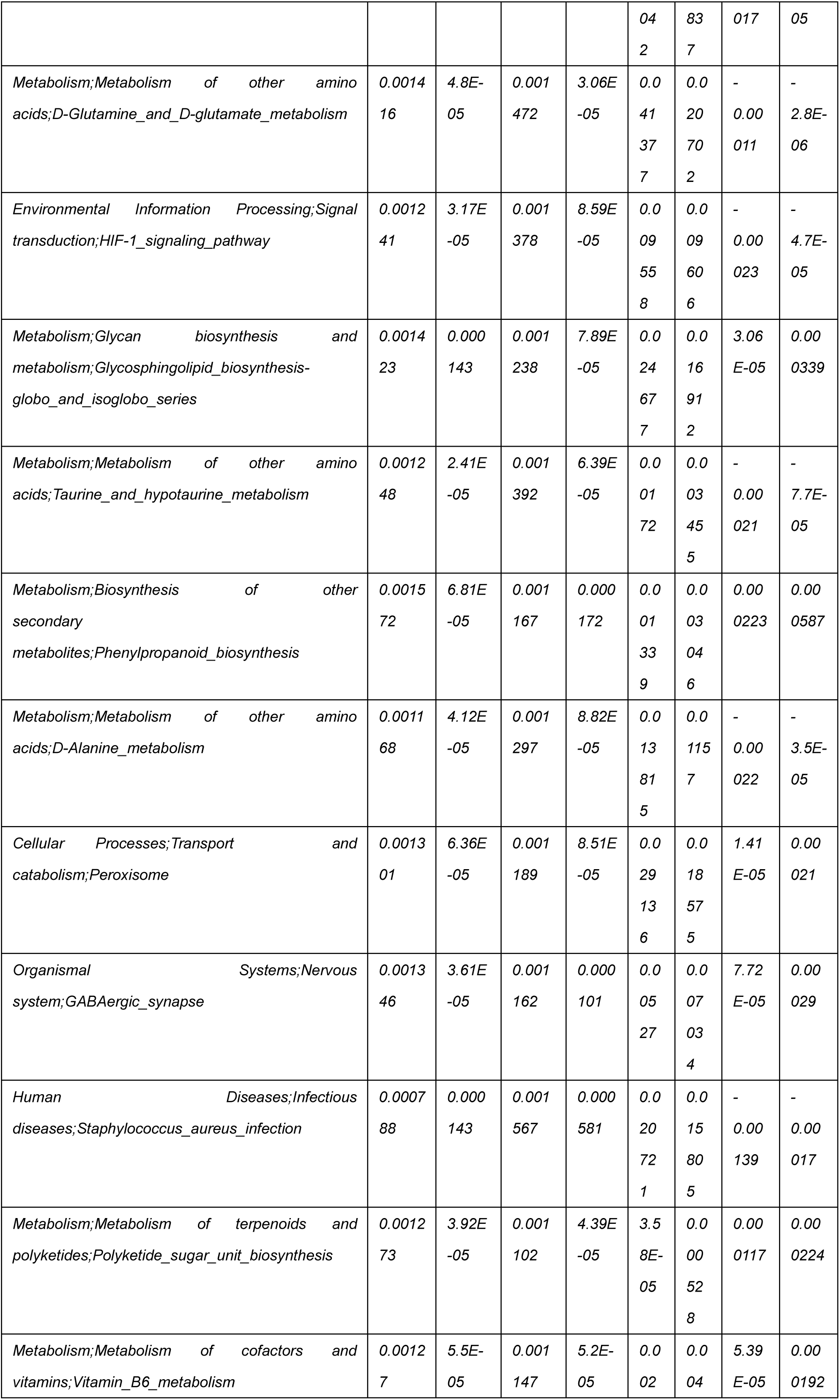
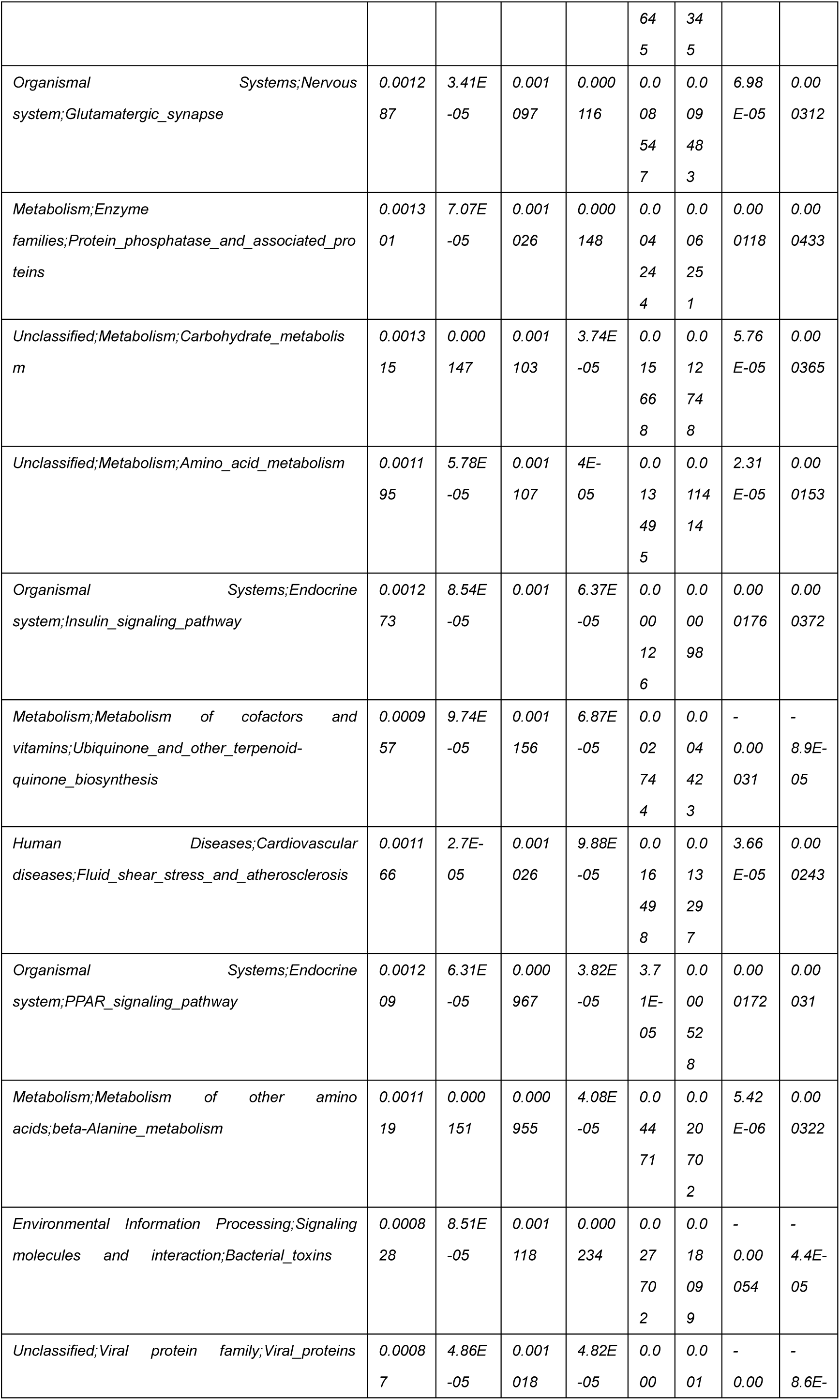
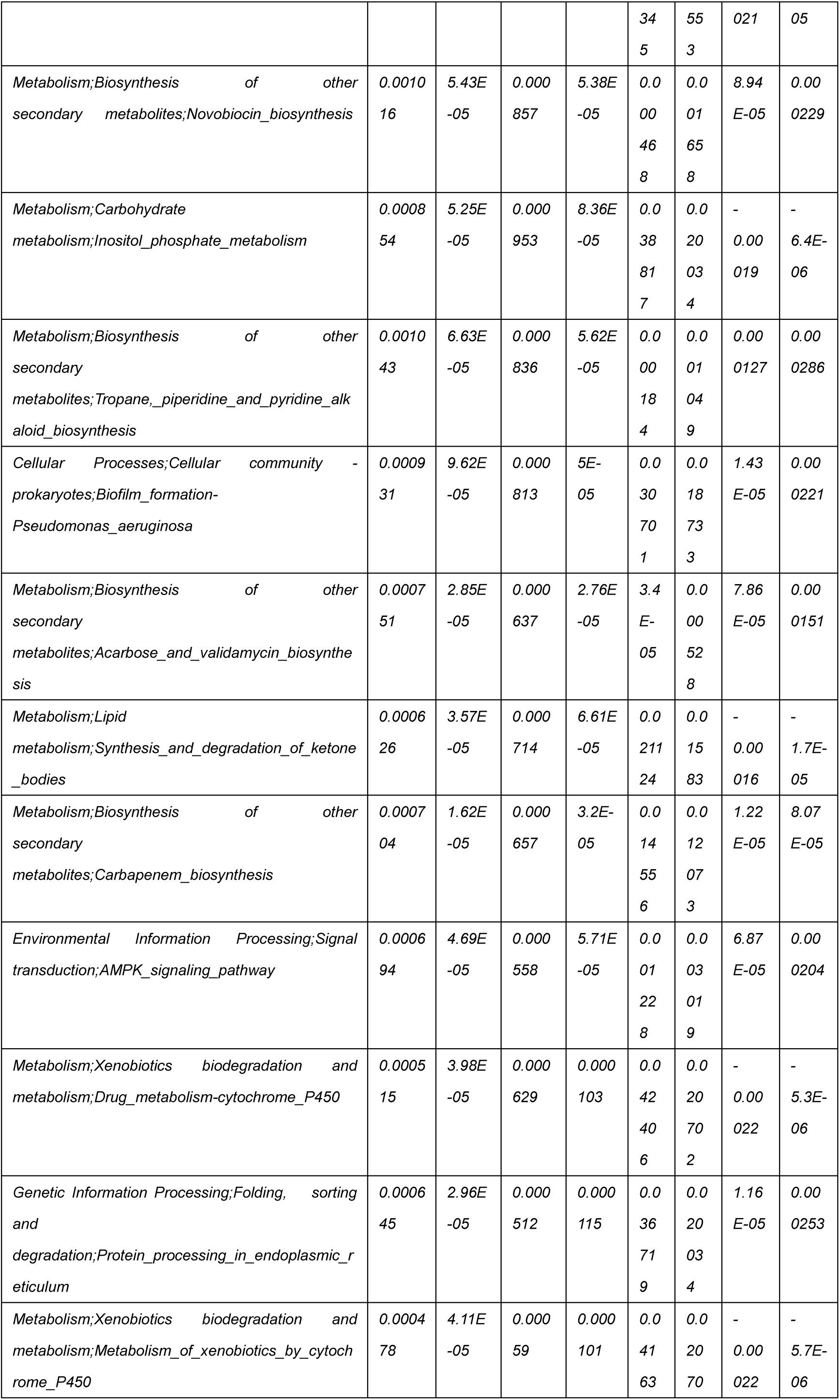
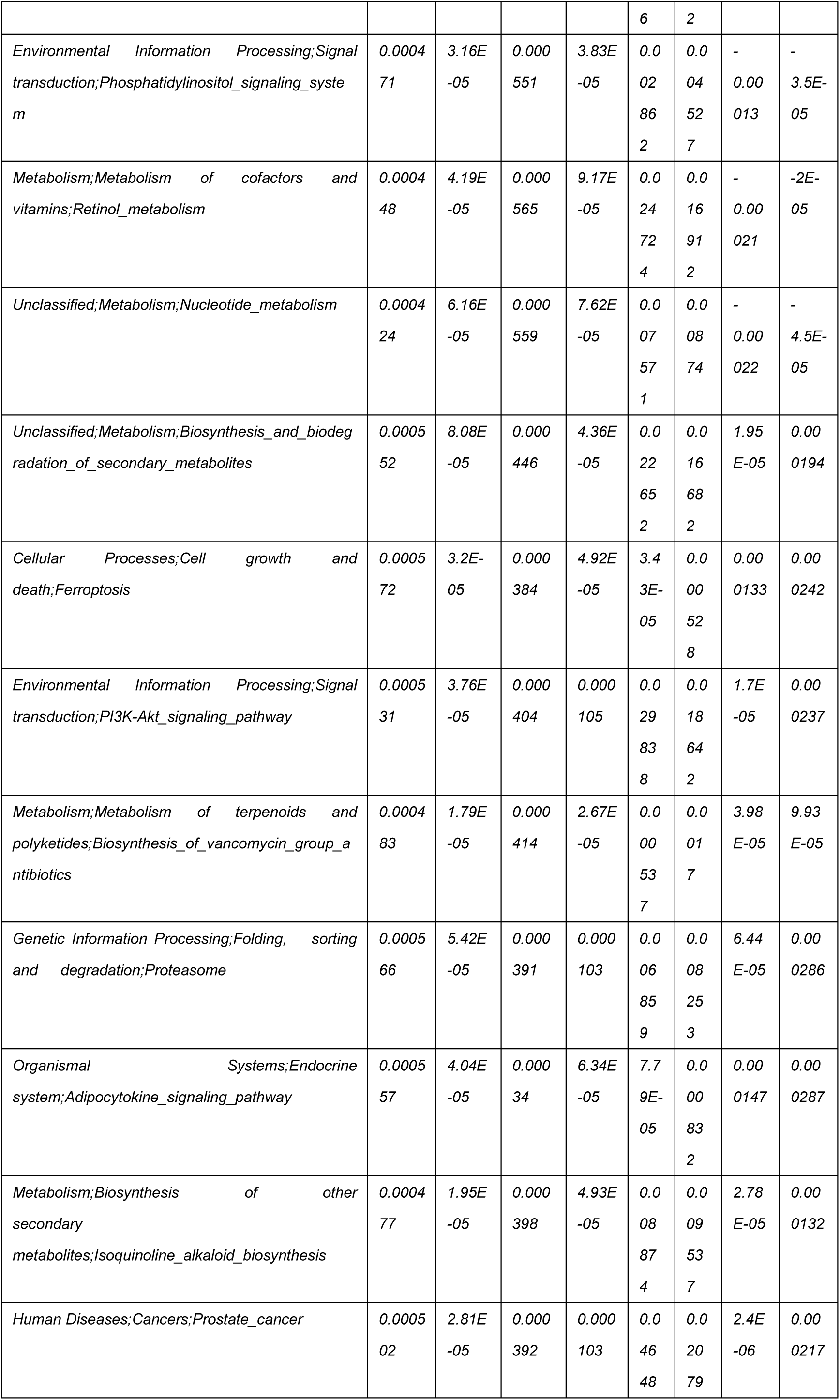
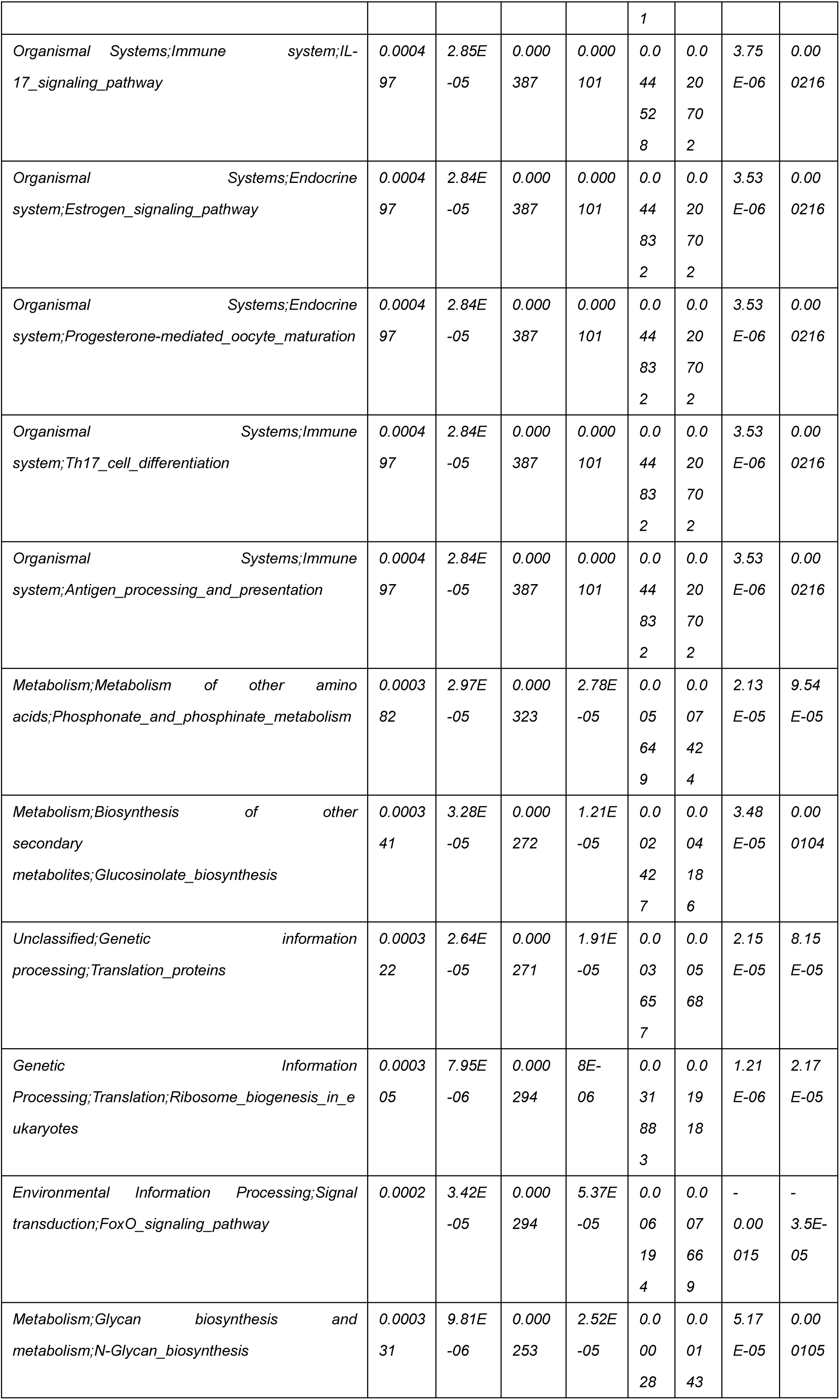
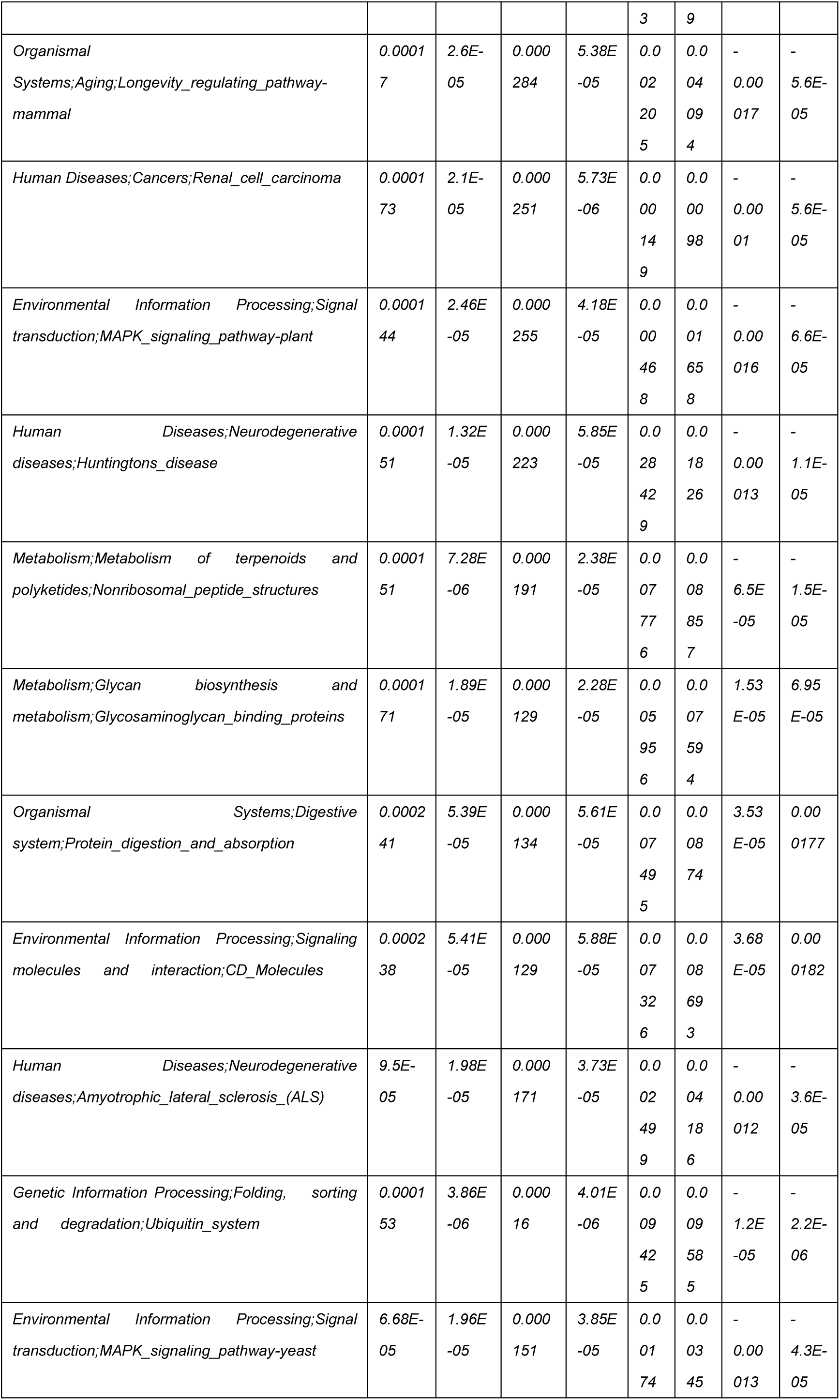
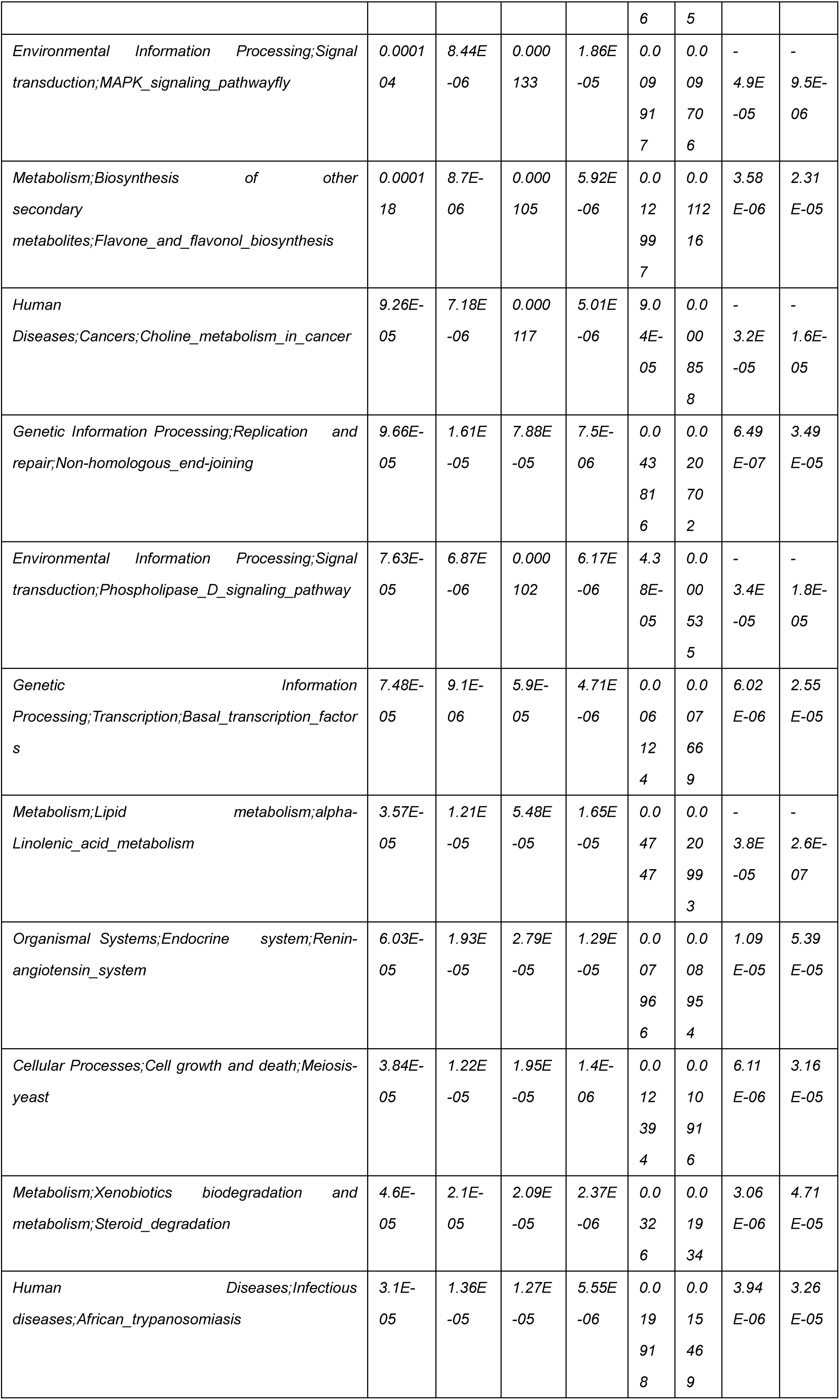
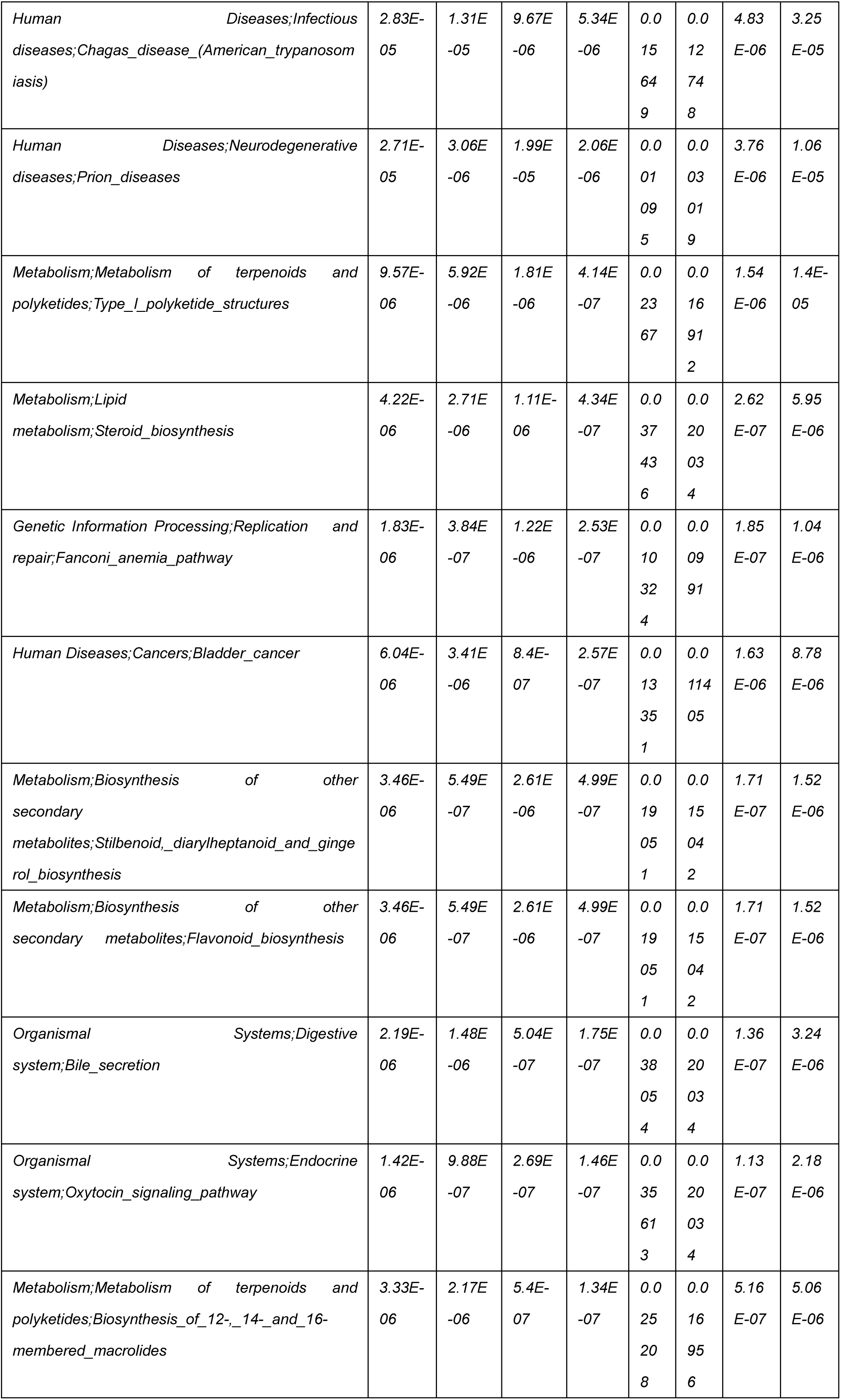
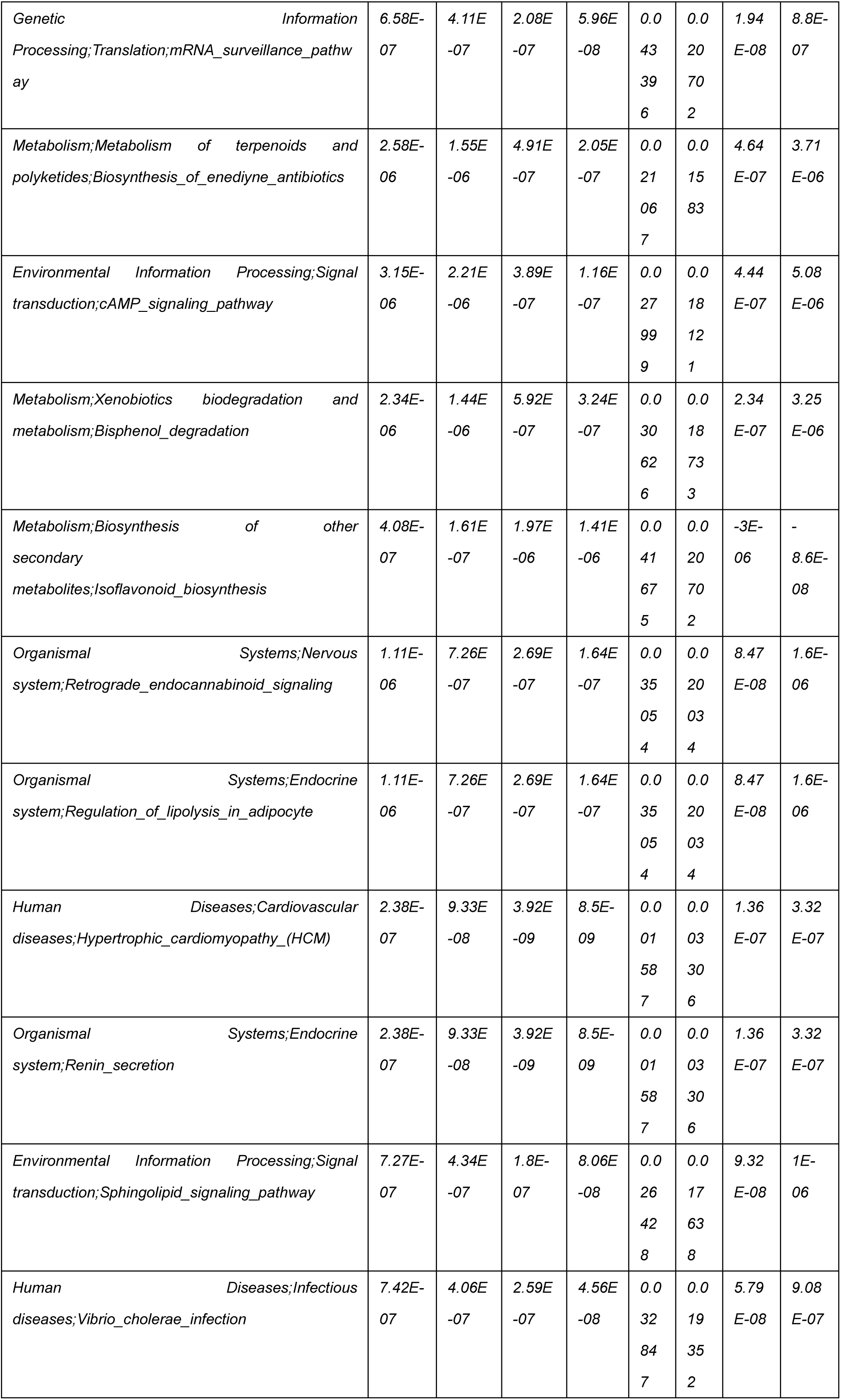
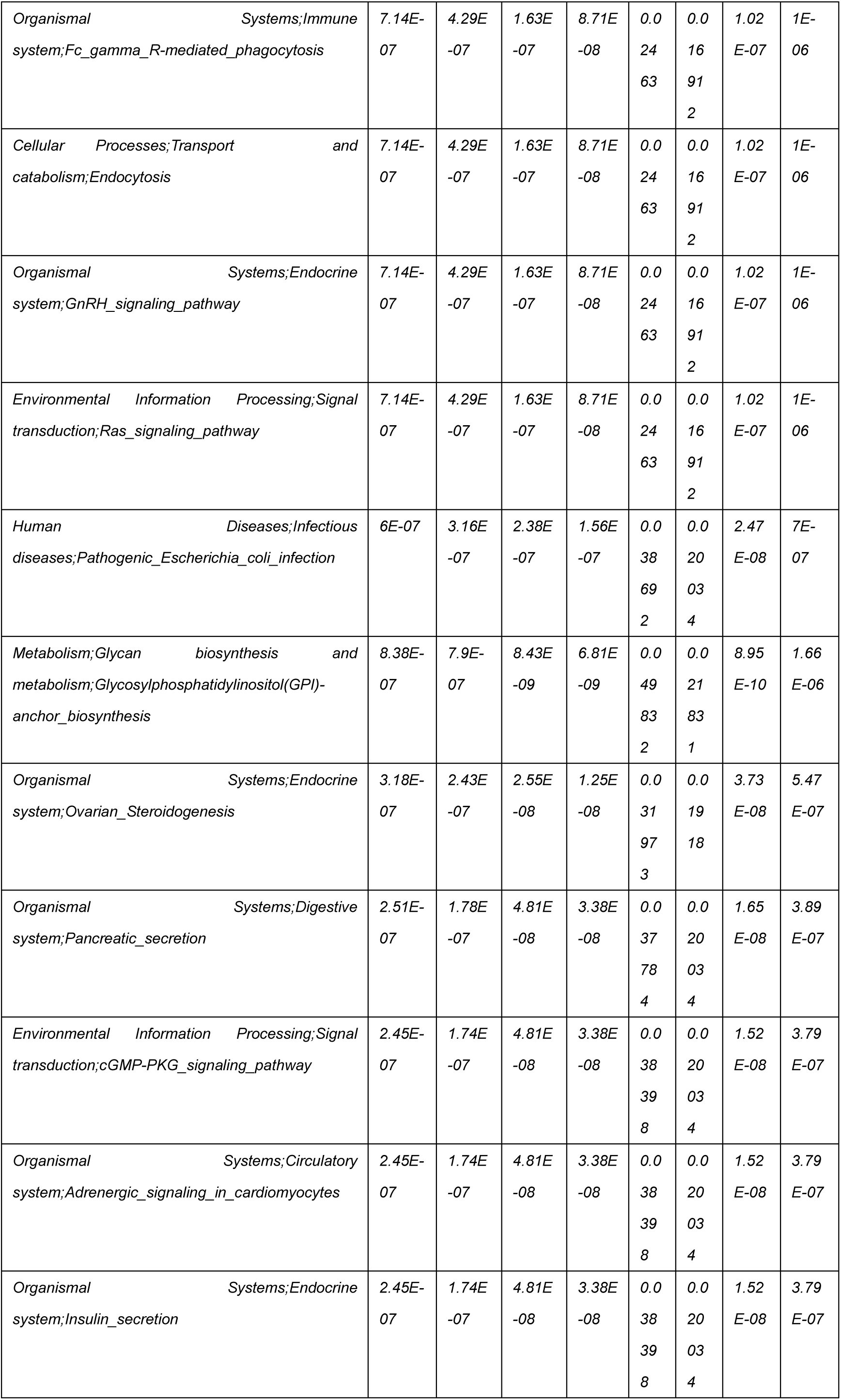
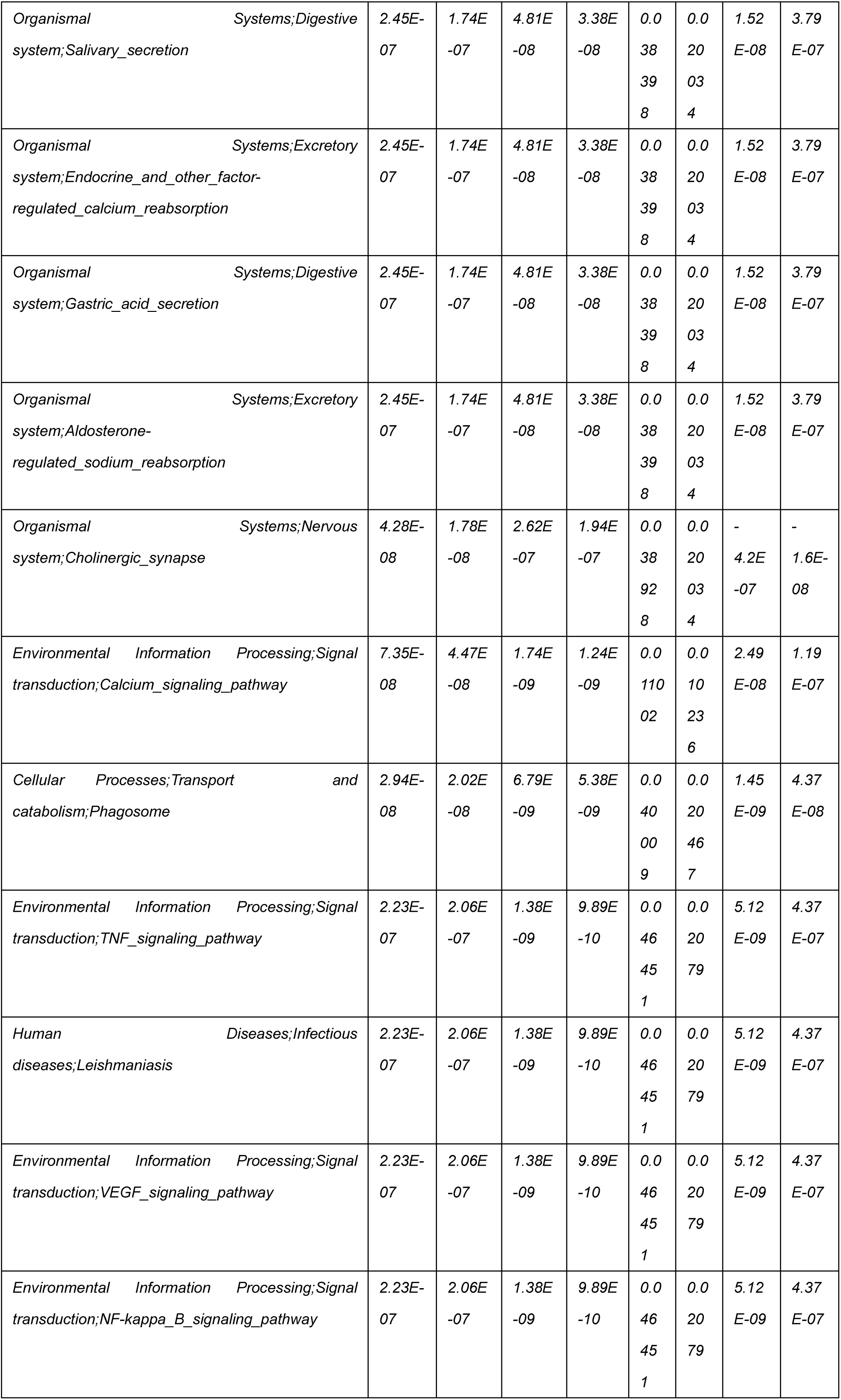
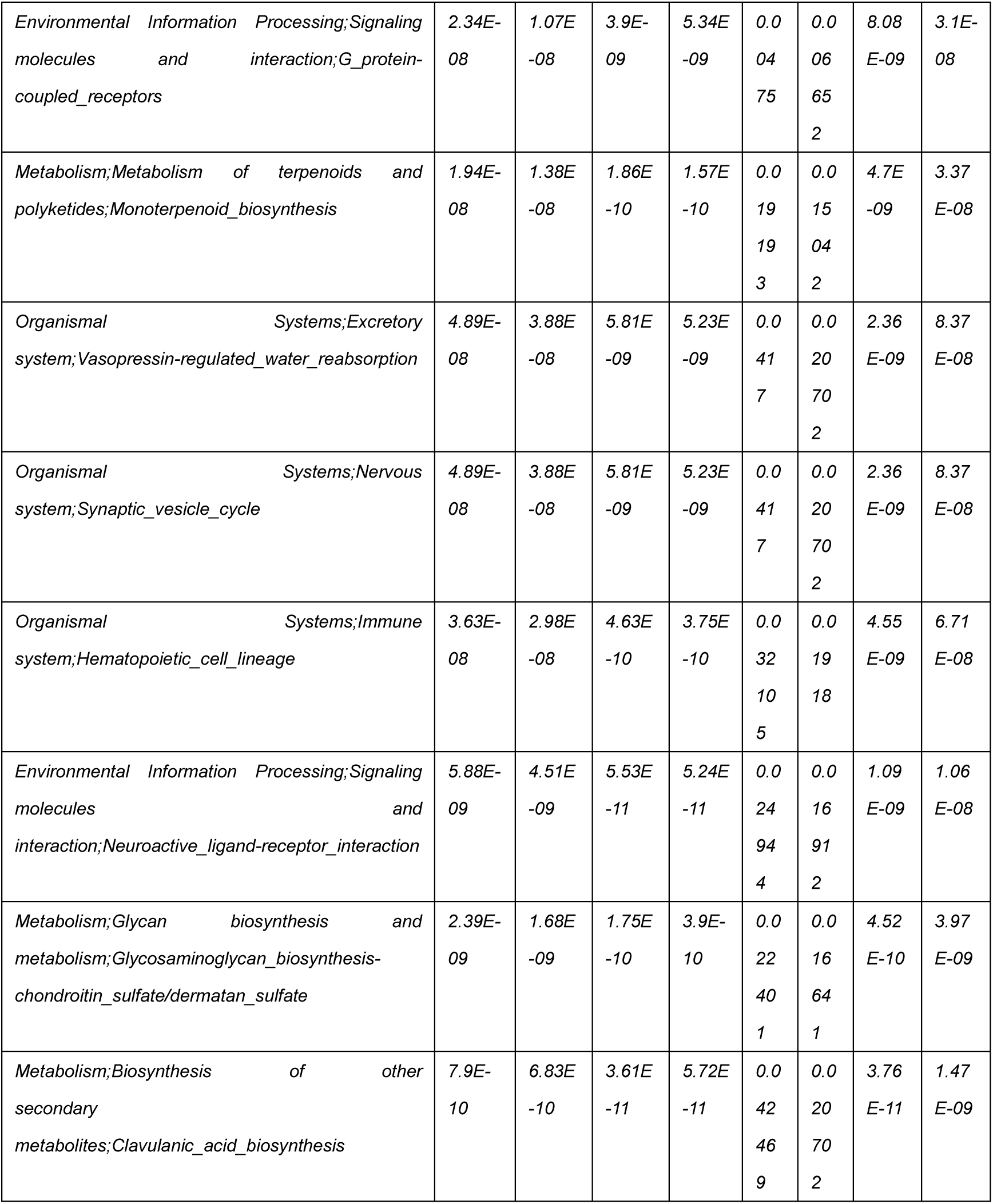
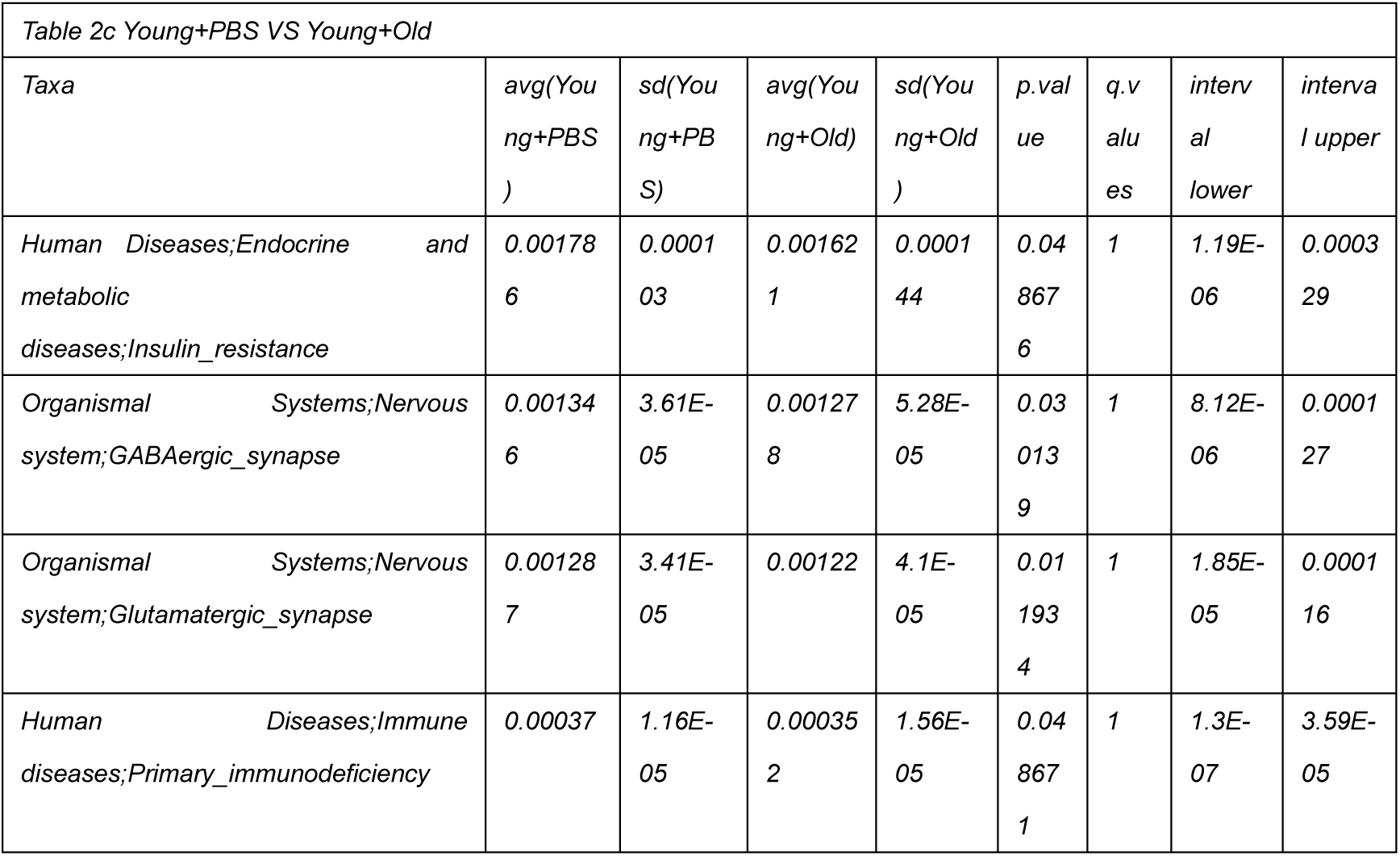
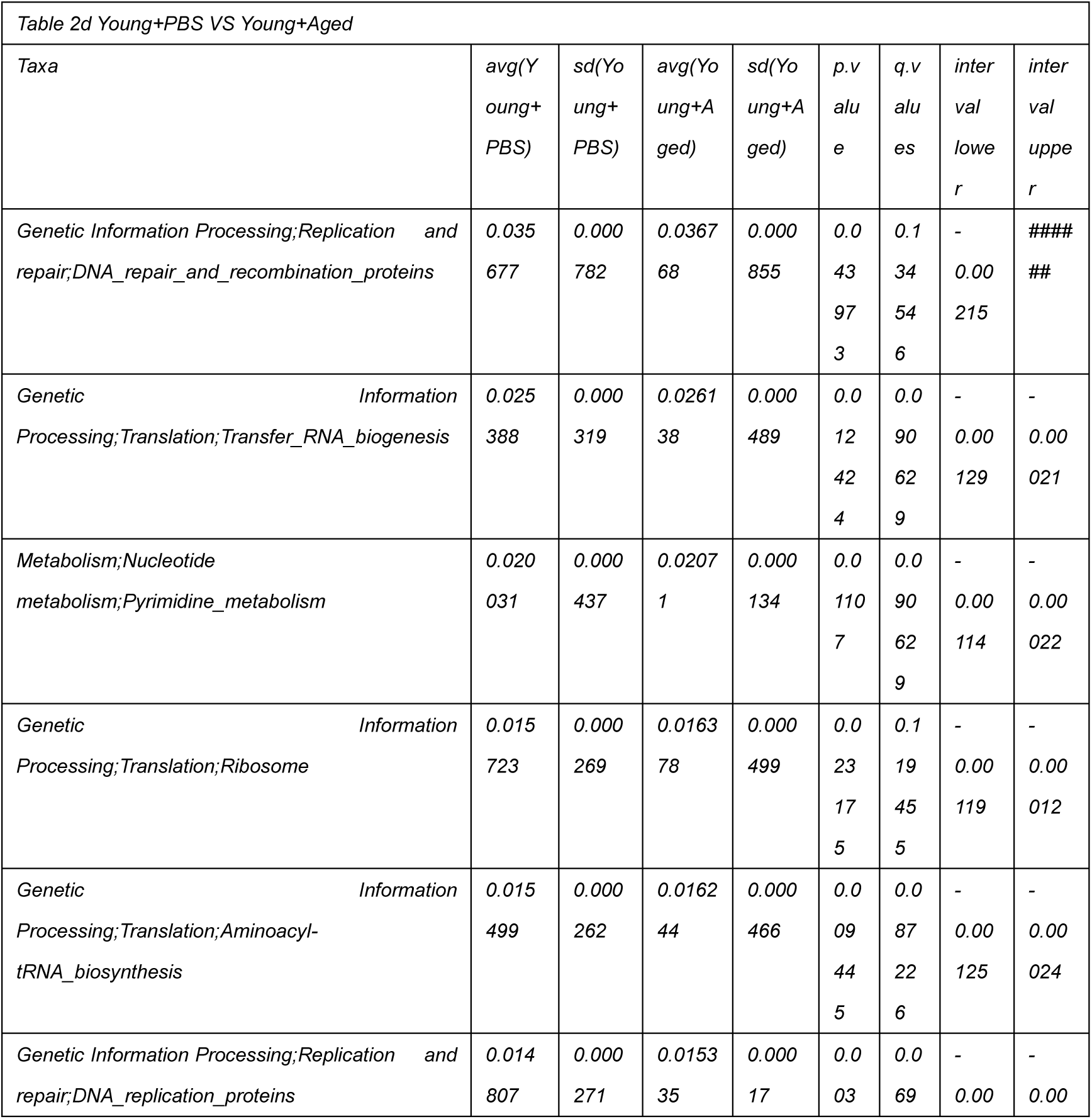
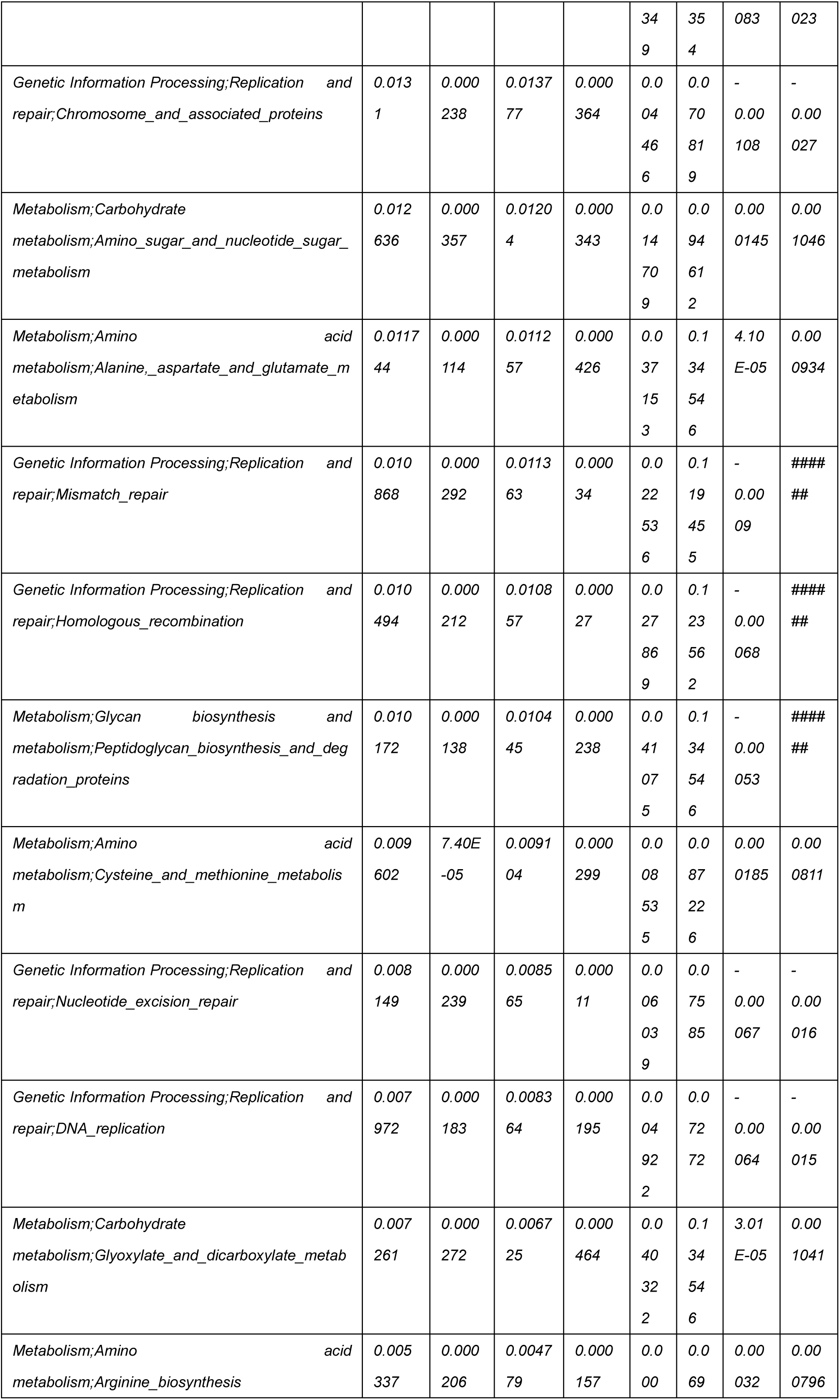
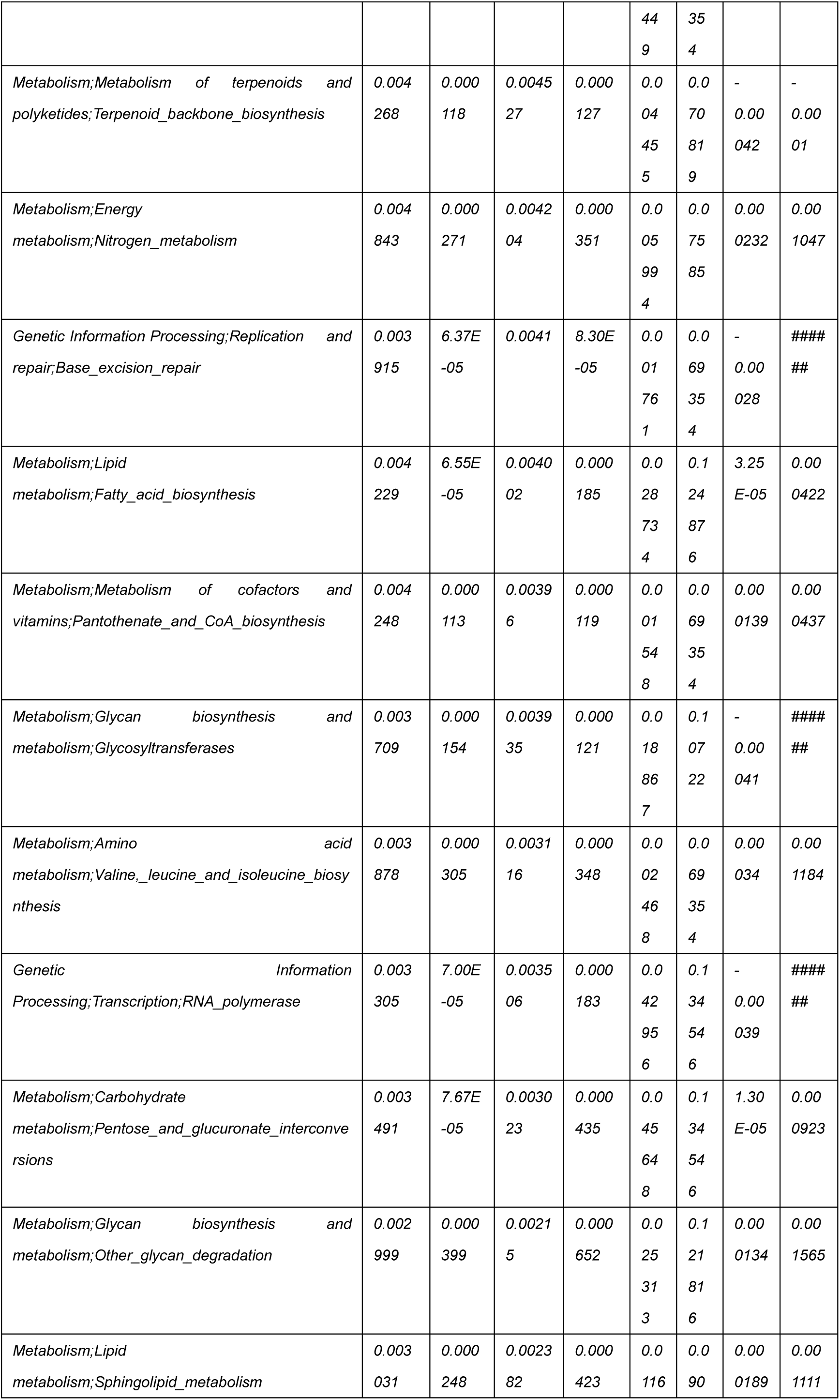
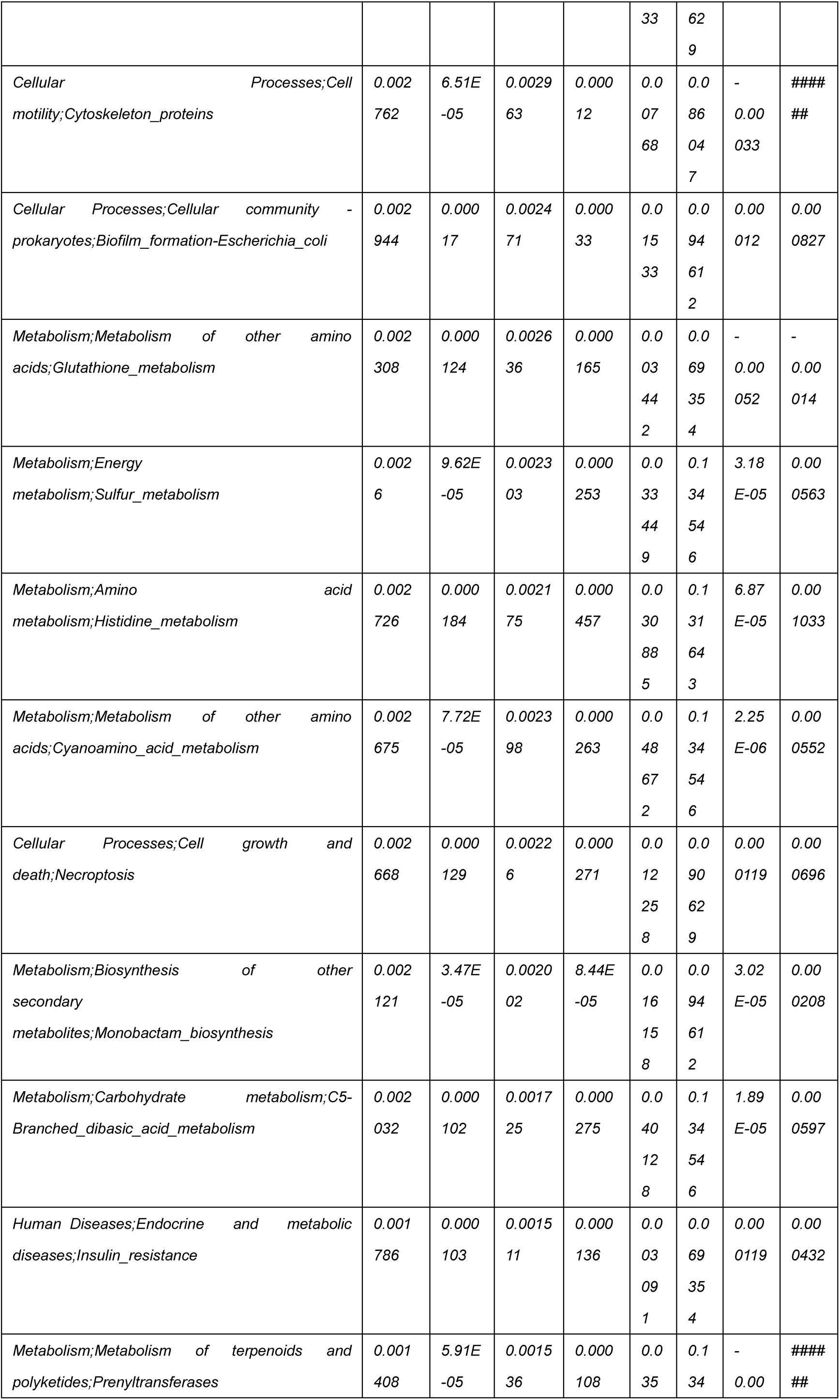
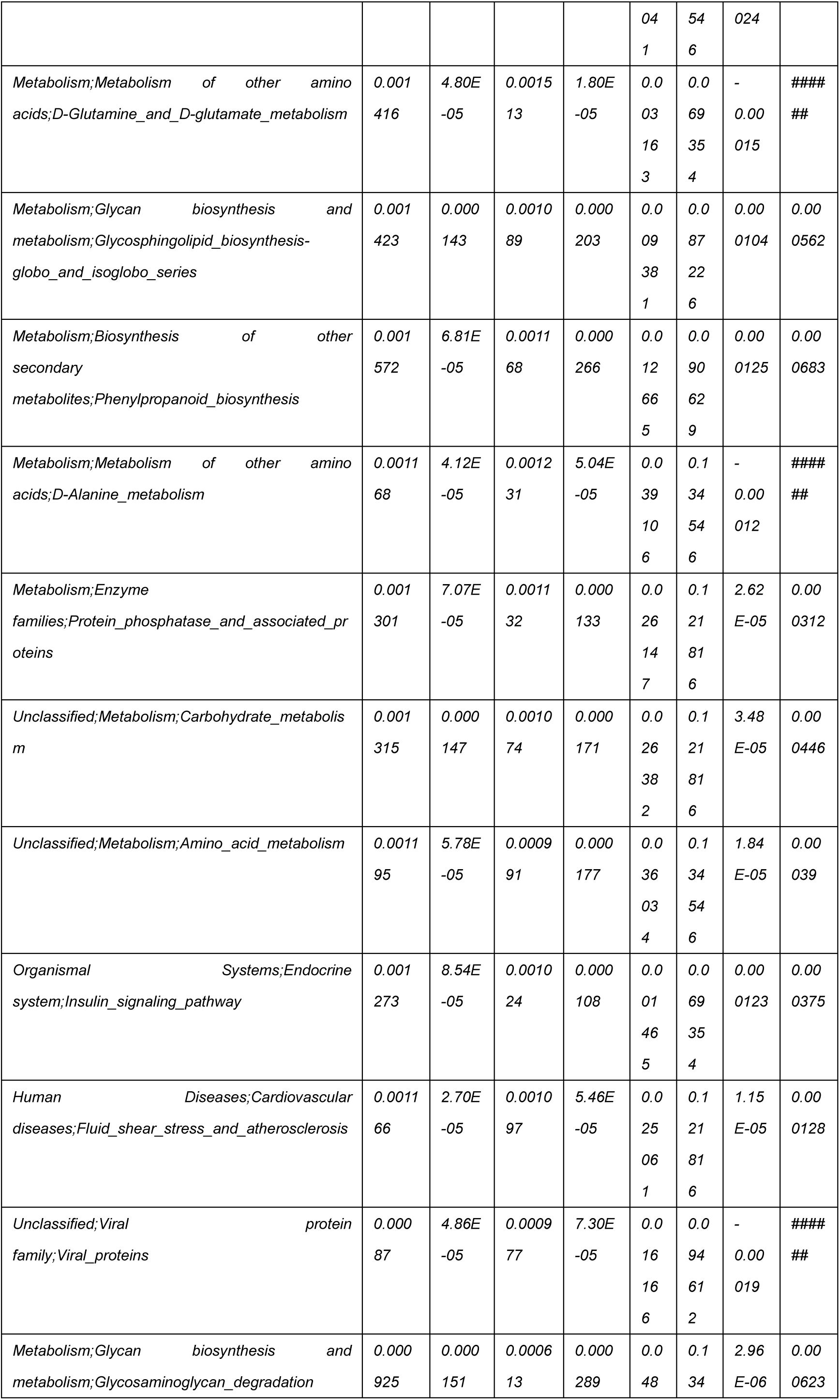
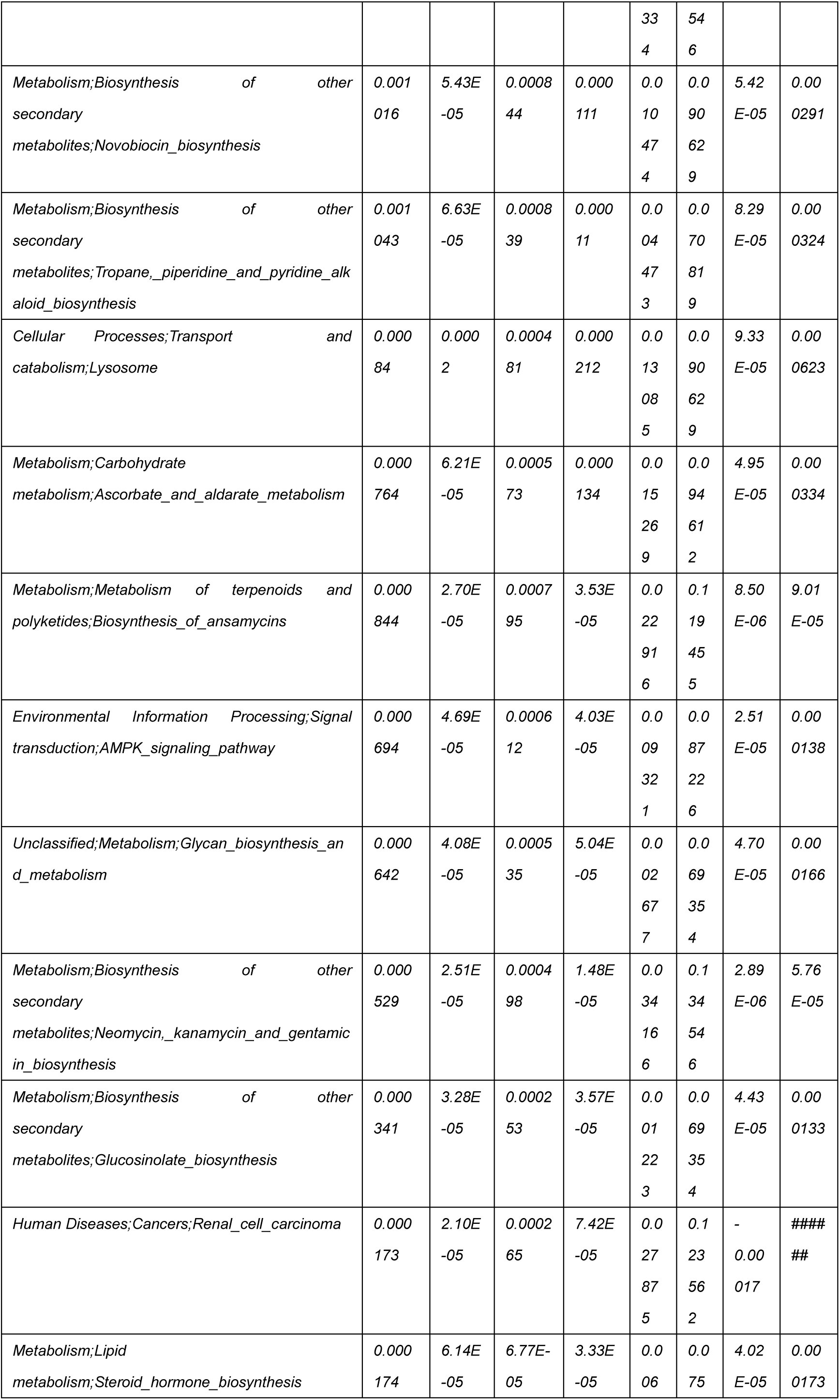
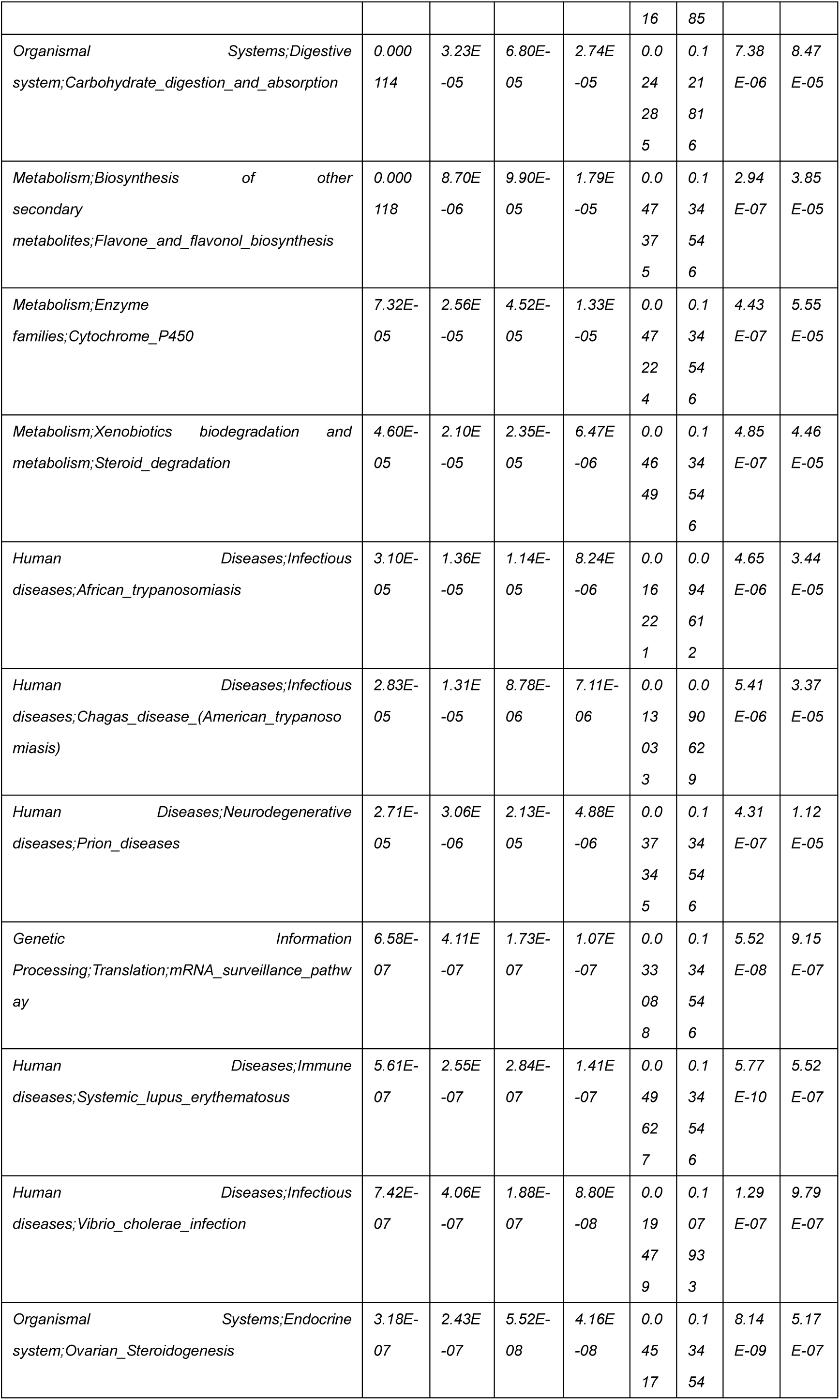
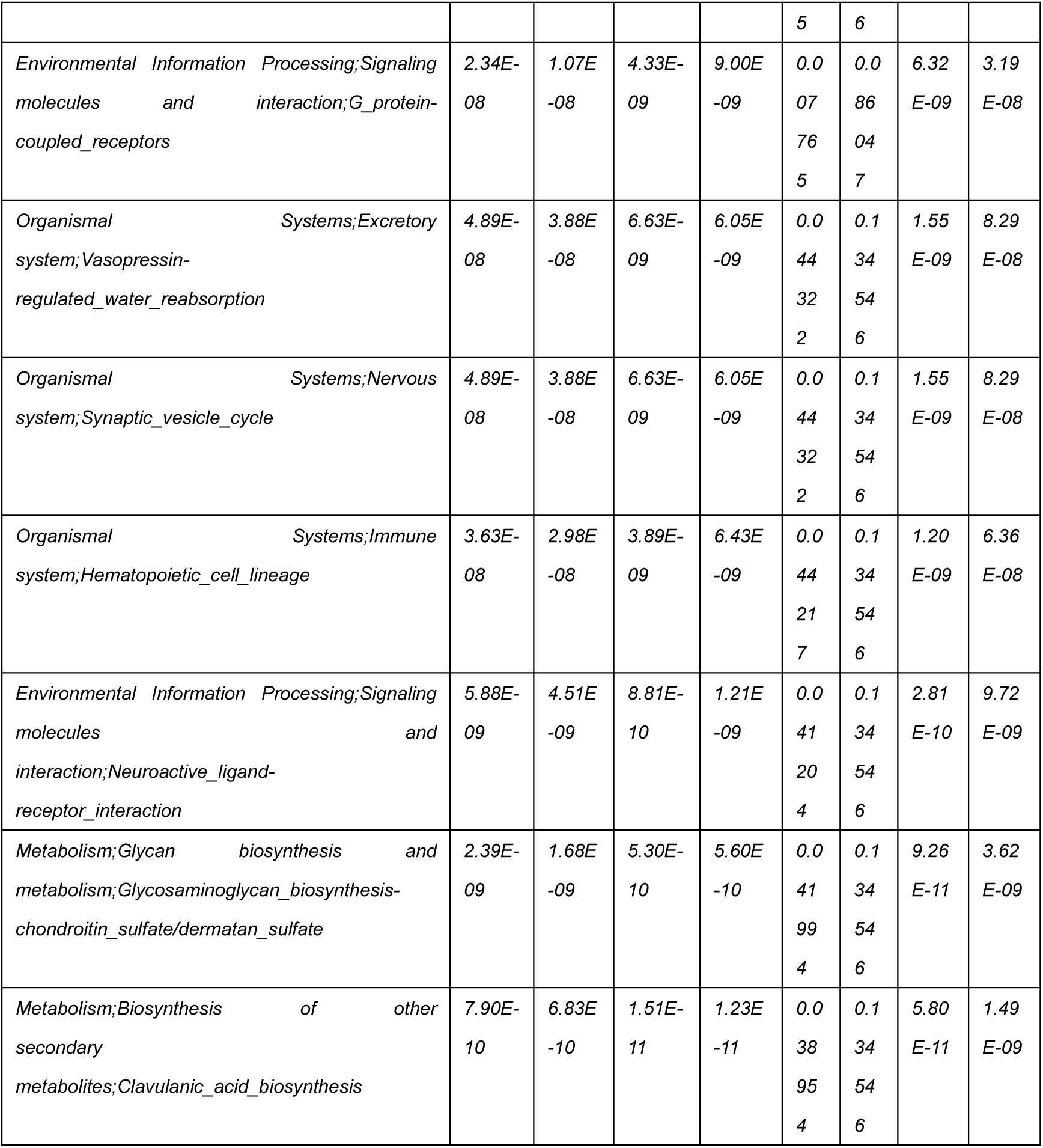
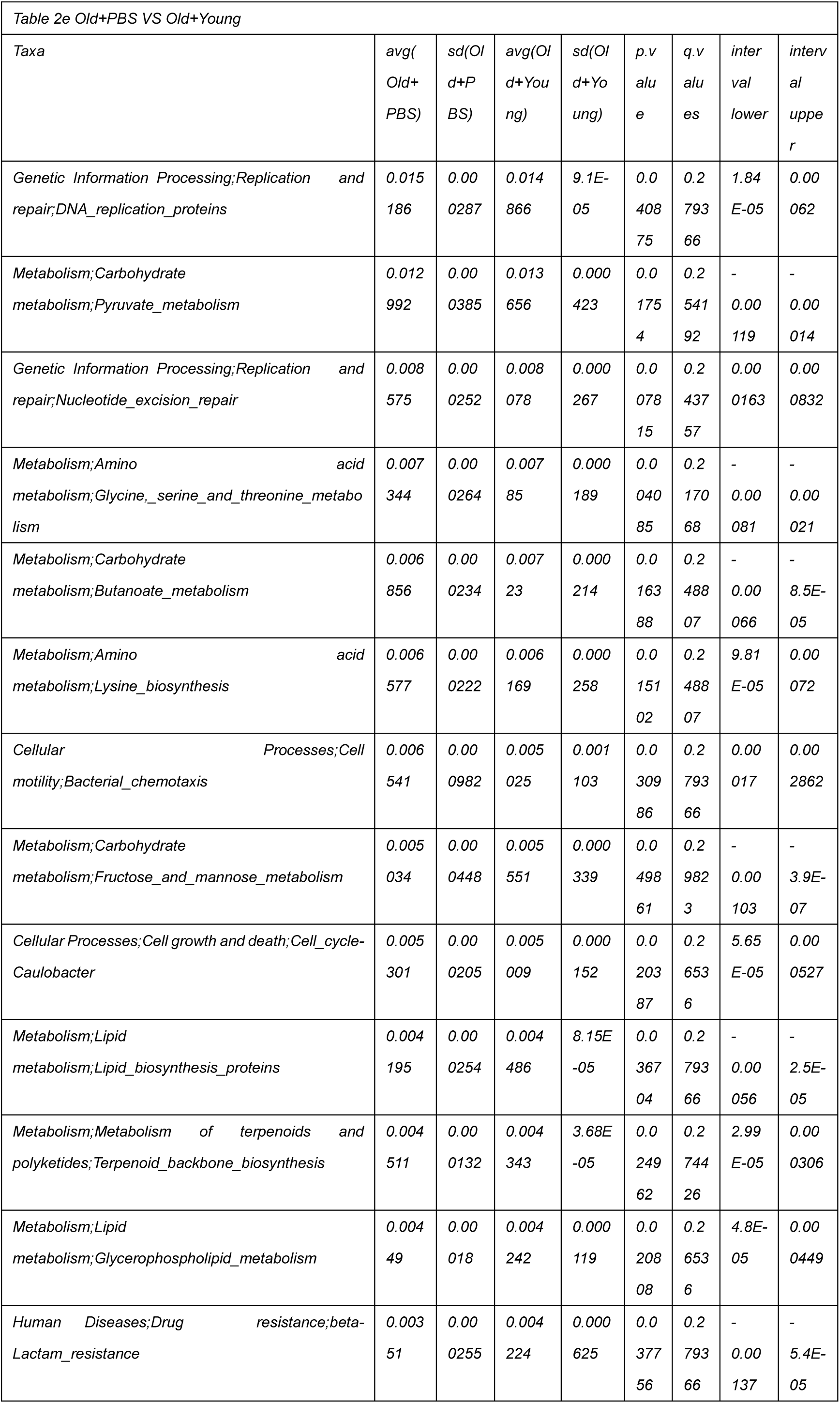
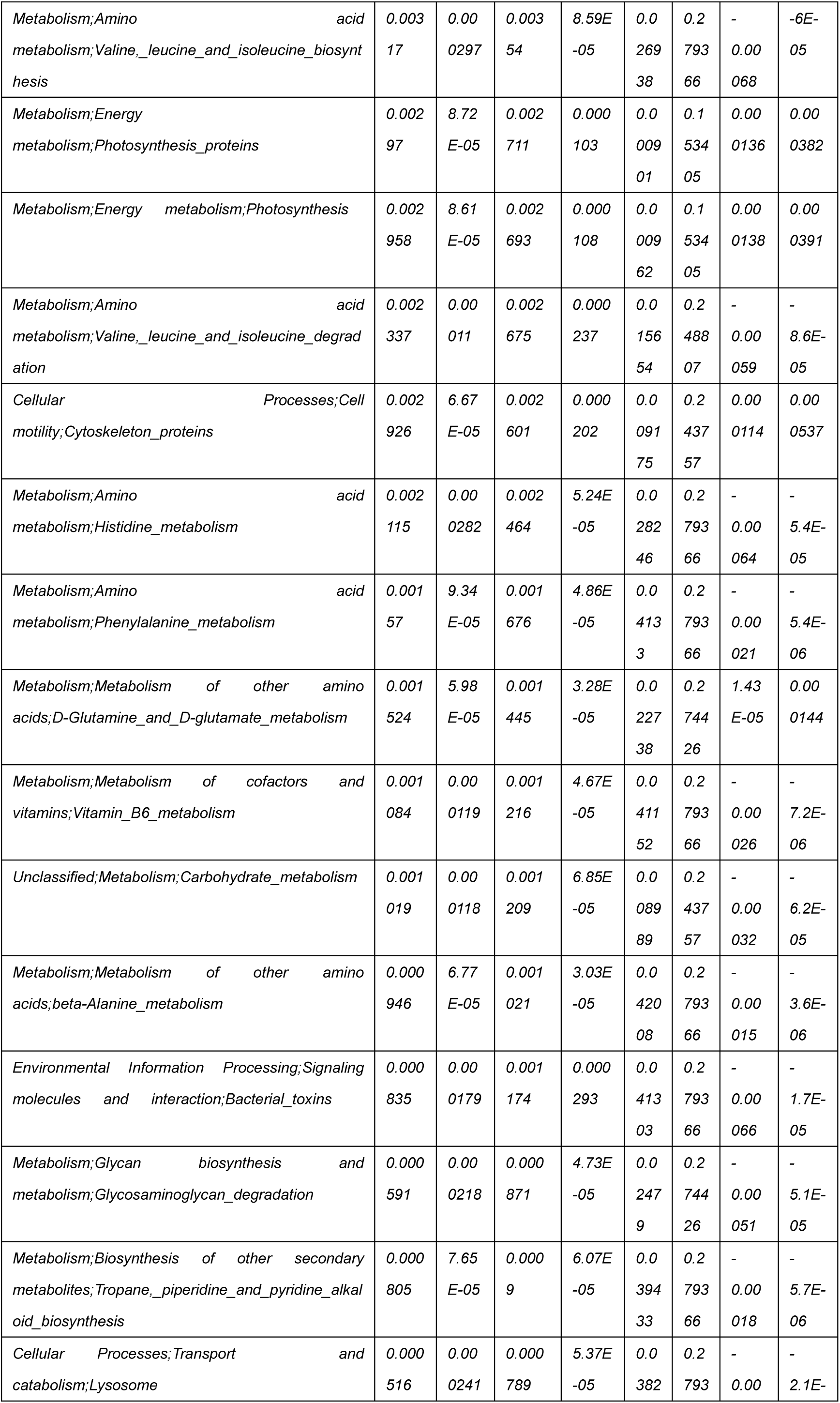
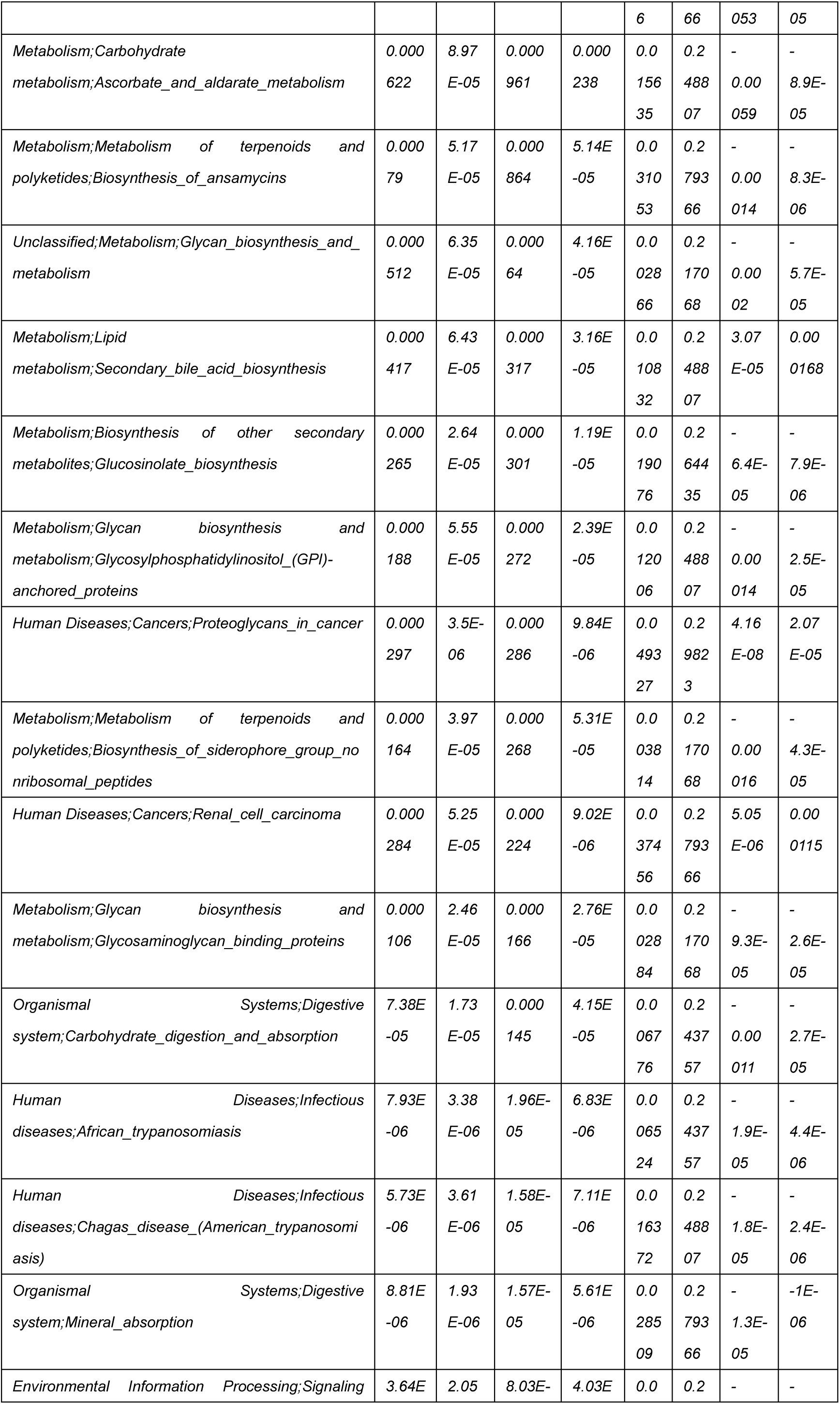
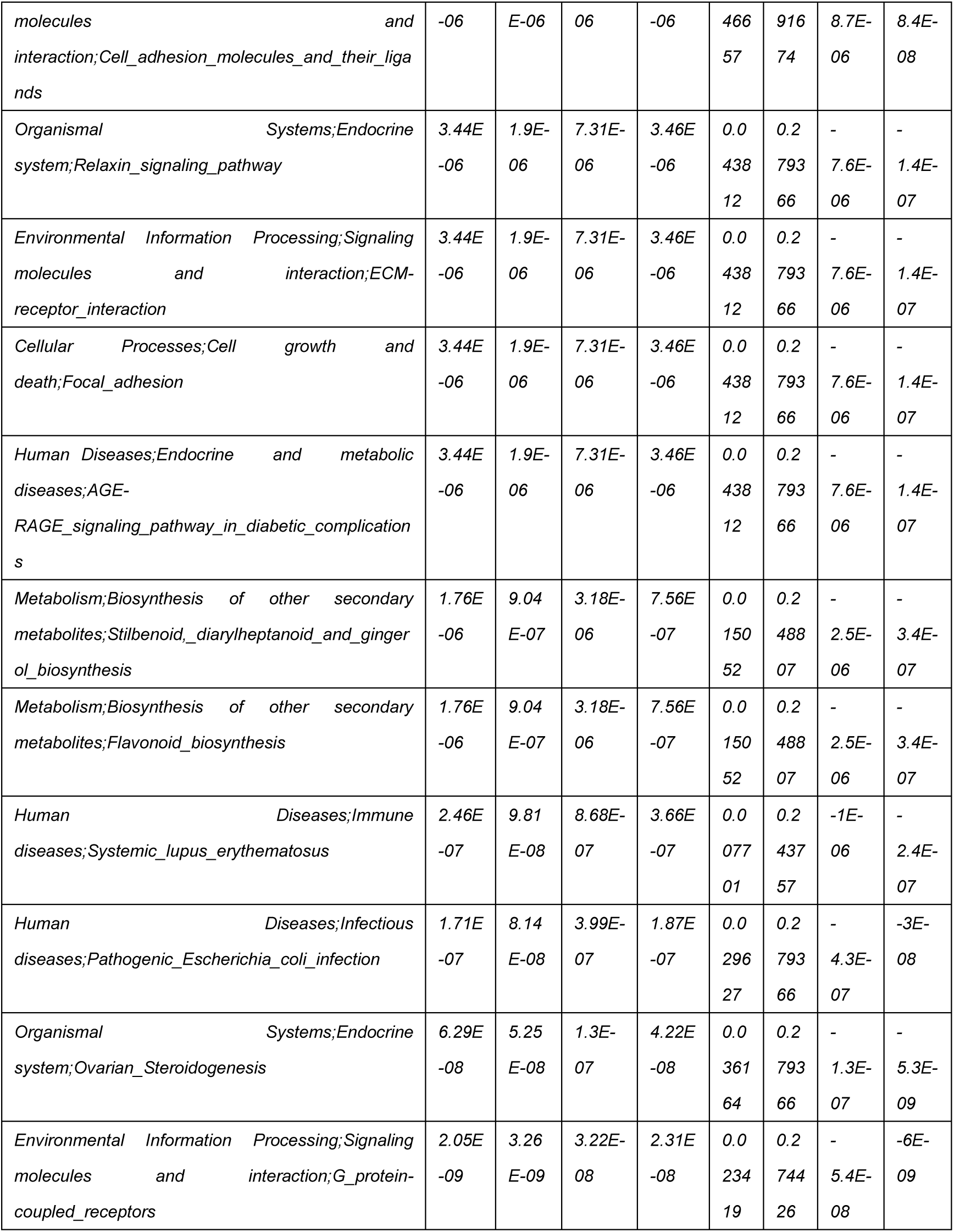
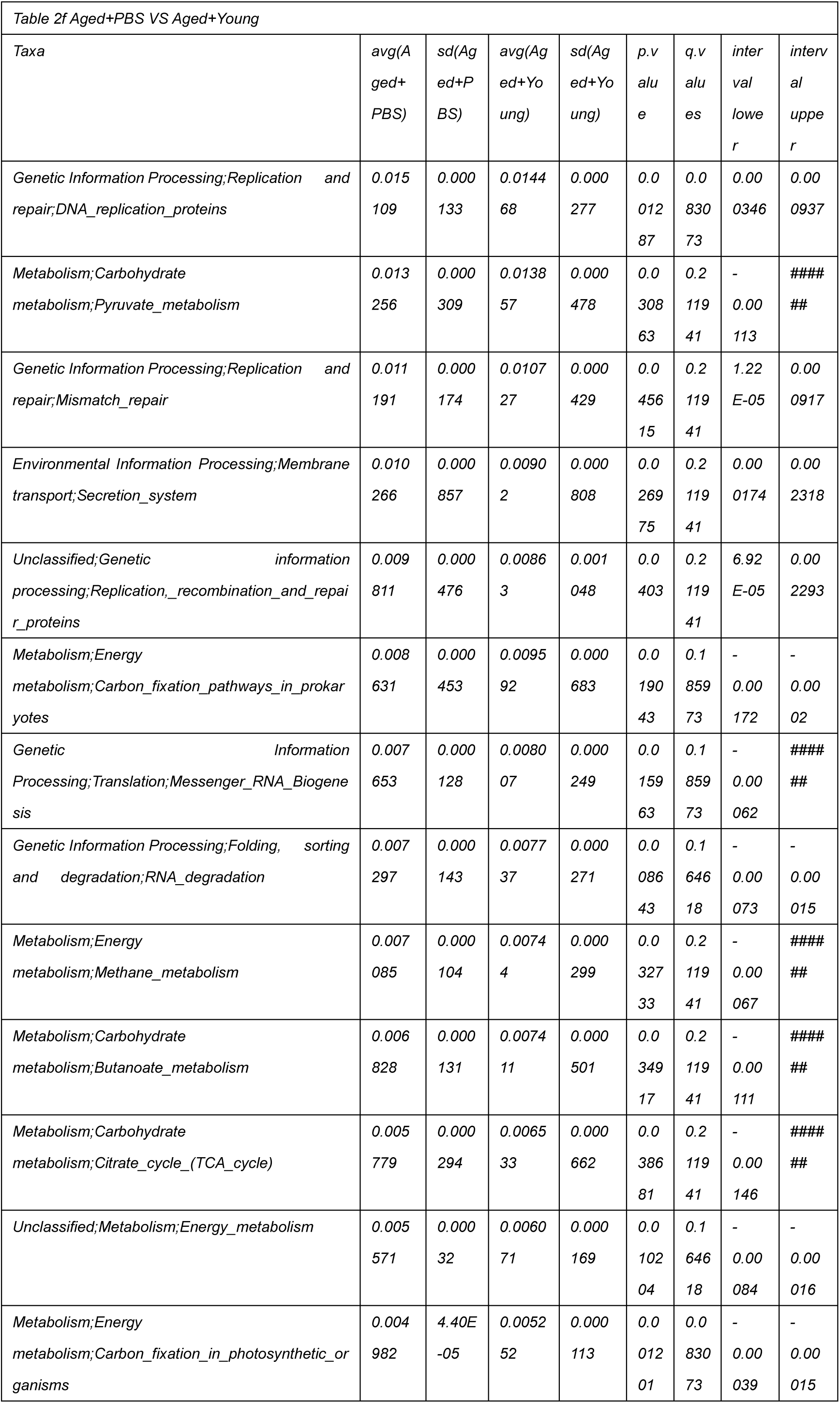
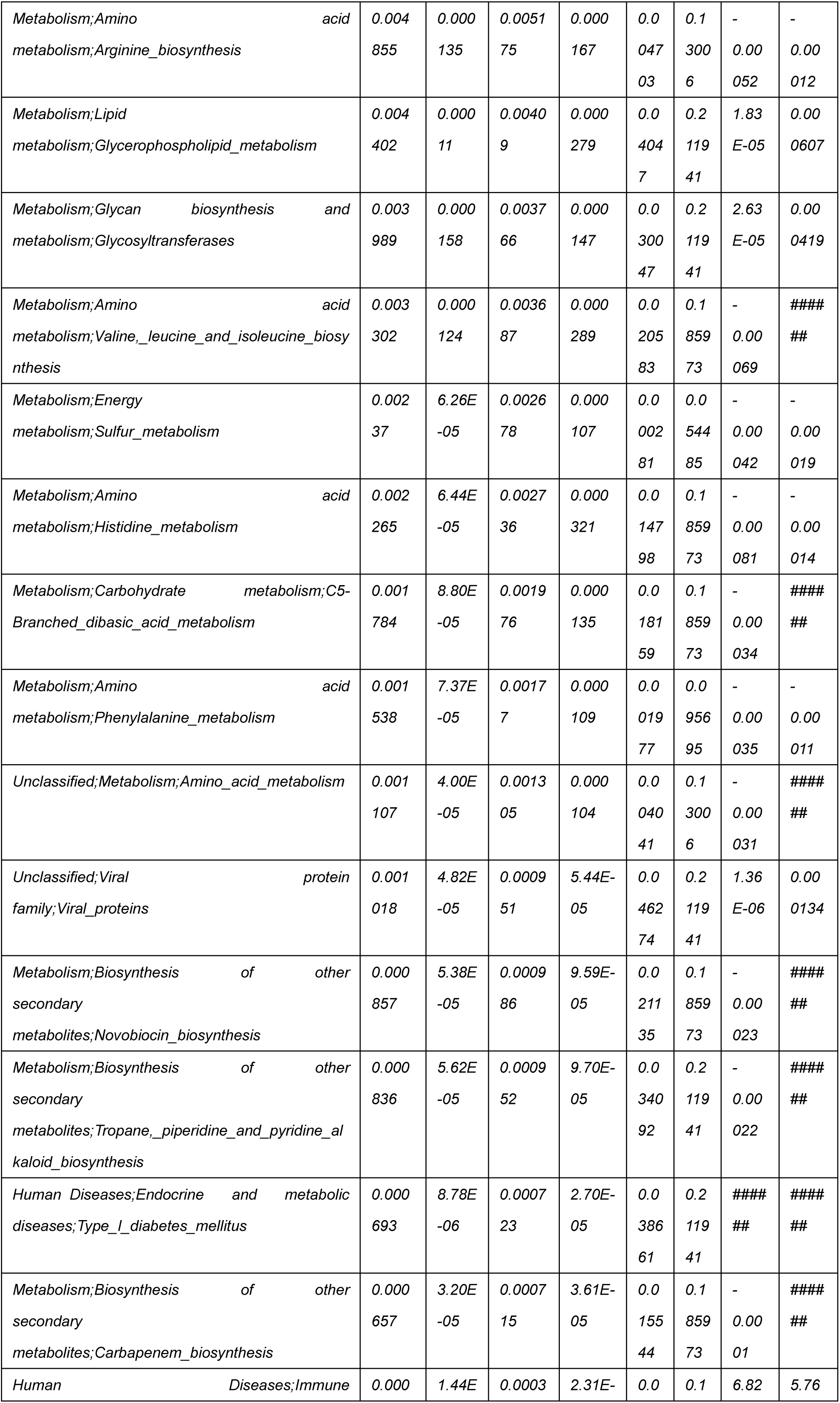
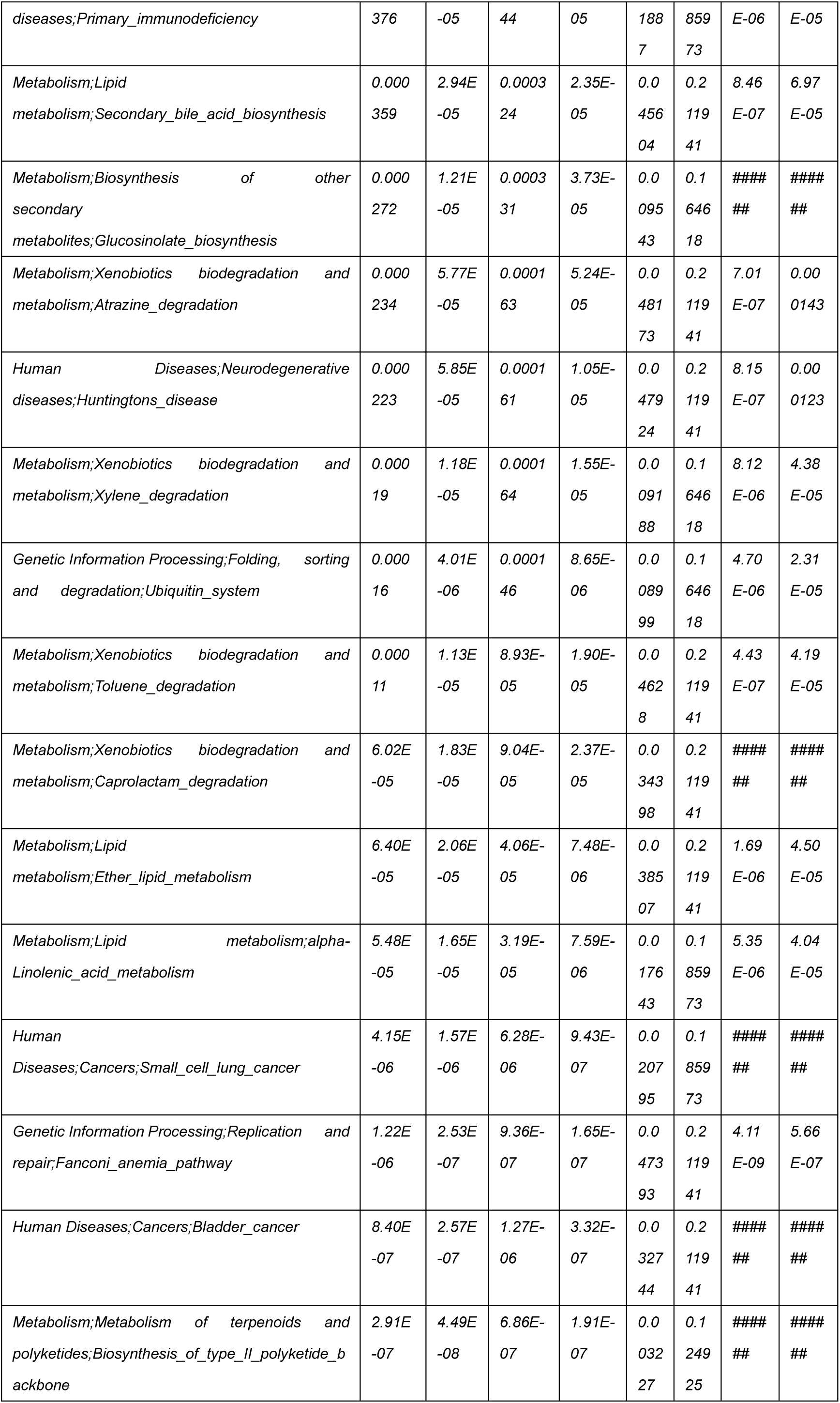
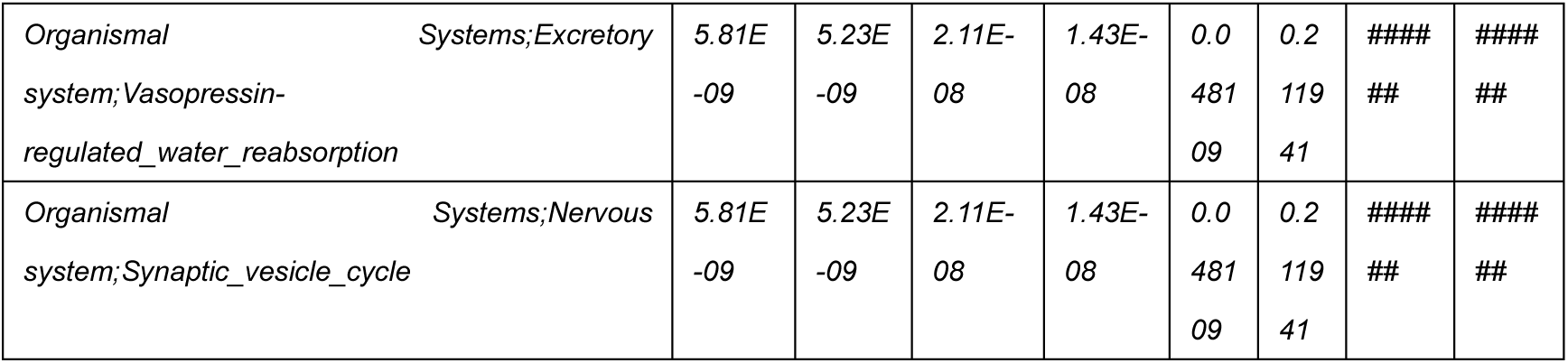
Changes in fecal microbiota metabolism and pathways after cross-age FMT.

### Cytokine determination

For detection of the IL-β, IL-6 and TNF-α concentrations in mouse foot tissue, foot tissue was harvested from different treatment groups, and 10% tissue homogenates were prepared using RIPA buffer. The prepared samples were then centrifuged at 3,000 × g and 4°C for 10 min, and supernatants were collected. All prepared foot tissue supernatants were determined using IL-β, IL- 6 and TNF-α ELISA kits at proper dilutions according to the manufacturer’s instructions (BioLegend, CA, USA). For the detection of cytokines in serum and peritoneal fluid, samples were separated by centrifugation (500 × g, 4°C, 10 min) and detected using IL-β, IL-6 and TNF-α ELISA kits according to the manufacturer’s instructions (BioLegend, CA, USA).

### Serum biochemistry analysis

The levels of serum uric acid (SUA), serum alanine aminotransferase (ALT), serum aspartate aminotransferase (AST), serum creatinine (Cr) and blood urea nitrogen (BUN) were measured using enzymatic kits from Nanjing Jiancheng Bioengineering Institute, Nanjing, China.

### Assay of uric acid synthesis enzymes

To detect the activity of Adenosine deaminase (ADA) and Guanine Deaminase (GDA) in the mouse liver, the liver was collected from mice in the various treatment groups and homogenized in sterile physiological saline solution (10-fold weight ratio of tissue). Following preparation, the samples were subjected to centrifugation at 3,000 × g and 4°C for 10 minutes, and the supernatants were collected. The activity of ADA and GDA in the liver supernatants was determined using ELISA kits (FEIYA BIOTECHNOLOGY, China) according to the manufacturer’s instructions with appropriate dilutions. To measure the activity of xanthine oxidase (XOD) in the liver and kidney of mice, tissues were collected from mice in the different treatment groups and homogenized in the buffer provided in the kit (10-fold weight ratio of tissue). The samples were then centrifuged at 8,000 × g and 4°C for 10 minutes, and the supernatants were collected. The activity of XOD in the liver and kidney supernatants was assessed using an ELISA kit (Beijing Solarbio Science & Technology Co., Ltd, China) according to the manufacturer’s instructions.

### Fecal microbiota composition analysis

For specific details, please refer to Supplementary material 1.

### Untargeted metabolomic analysis

For specific details, please refer to Supplementary Material 2.

### Analysis of short-chain fatty acids in faeces

For specific details, please refer to Supplementary Material 3.

### Statistical analysis

All values were expressed as the means ± SEM. Raw data were subjected to one-way ANOVA to evaluate statistical significance between at least three groups, and pairwise comparison was conducted using Student’s t-test. The results were considered statistically significant at p < 0.05.

## Data and code availability

The 16S rDNA sequencing data and untargeted metabolomic data presented in this study can be found in online repositories. The findings of this study have been deposited into the CNGB Sequence Archive (CNSA) of China National GenBank Database (CNGBdb) under the accession number CNP0004751. The names of the repository/repositories and accession number(s) can be found below: https://db.cngb.org/cnsa/project/CNP0004751_46c178da/reviewlink/.

The lead contact will provide original western blot images and microscopy data from this paper upon request.

No unique code was generated in this study.

For any further information needed to reanalyze the data presented in this paper, it can be obtained from the lead contact upon request.

## Declaration of interests

The authors declare no competing interests.

## Acknowledgments

This work was supported by the National Key R&D Program of China (No. 2023YFD1801100), the National Natural Science Foundation of China (Nos. 32202889 and 31972749) and the Natural Science Foundation of Jilin Province (No. 20220101304JC).

## Author contributions

The study was designed by Naisheng Zhang, Wenlong Zhang, and Hang Gao. Ning Song conducted all mouse experiments and performed the statistical analysis. Mice sample collection was carried out by Jianhao Li, Mingze Wang and Yi Liu. Ning Song was responsible for software applications, investigation, and writing the original draft. Jianhao Li and Mingze Wang helped with validation and analysis. Ning Song and Yi Liu were responsible for resources and data curation. Wenlong Zhang and Zhiming Ma wrote and revised the final draft of the manuscript, and all authors participated in its revision and gave their approval.

---

**Supplementary Fig 1.**
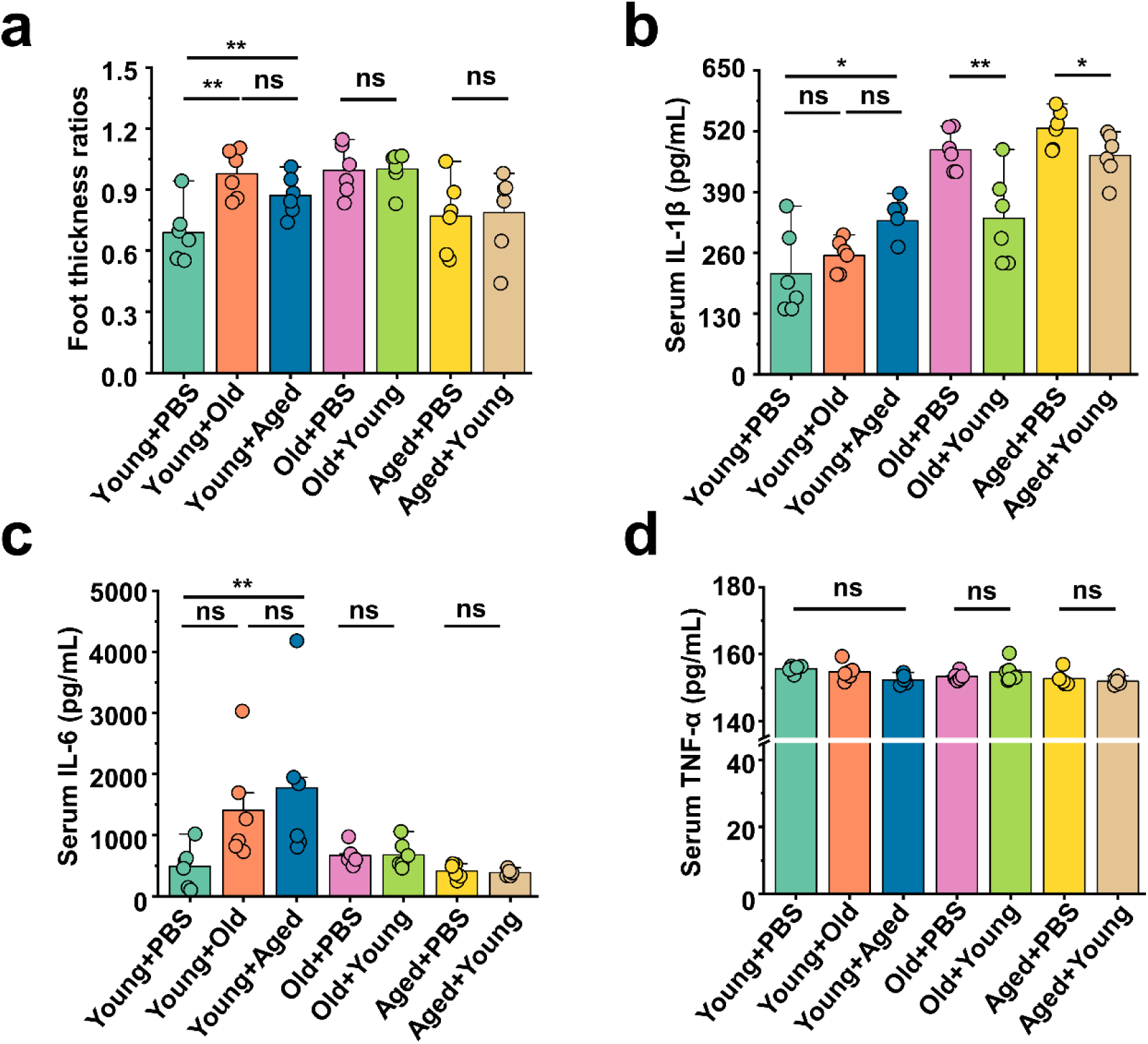
(a) All groups’ foot thickness ratios were tested after MSU administration (n=6). (b-d) The serum concentrations of IL-1β (b), IL-6 (c) and TNF-α (d) inflammatory parameters were measured in the indicated mice(n=6). Values are presented as the mean ± SEM. Differences were assessed by t-test or One-Way ANOVA and denoted as follows: *p < 0.05, **p < 0.01, and ***p < 0.001, “ns” indicates no significant difference between groups.

**Supplementary Fig 2.**
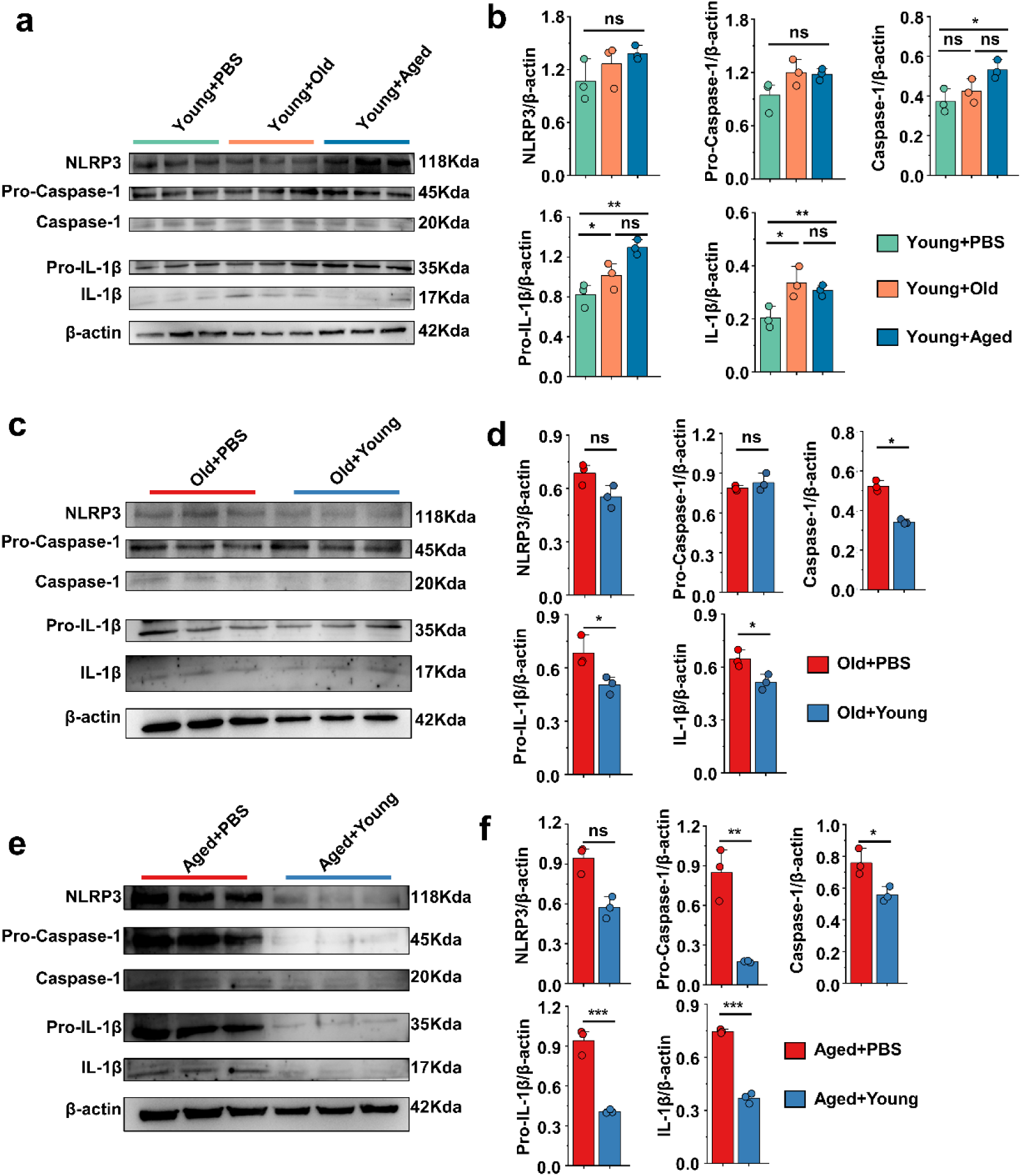
(a-b) Representative western blot images and band density (Young+PBS, Young+Old and Young+Aged) of peritoneal cells NLRP3 pathways proteins (n=3). (c-d) Representative western blot images and band density (Old+PBS and Old+Young) of peritoneal cells NLRP3 pathways proteins (n=3). (e-f) Representative western blot images and band density (Aged+PBS and Aged+Young) of peritoneal cells NLRP3 pathways proteins (n=3). Values are presented as the mean ± SEM. Differences were assessed by t-test or One-Way ANOVA and denoted as follows: *p < 0.05, **p < 0.01, and ***p < 0.001, “ns” indicates no significant difference between groups.

**Supplementary Fig 3.**
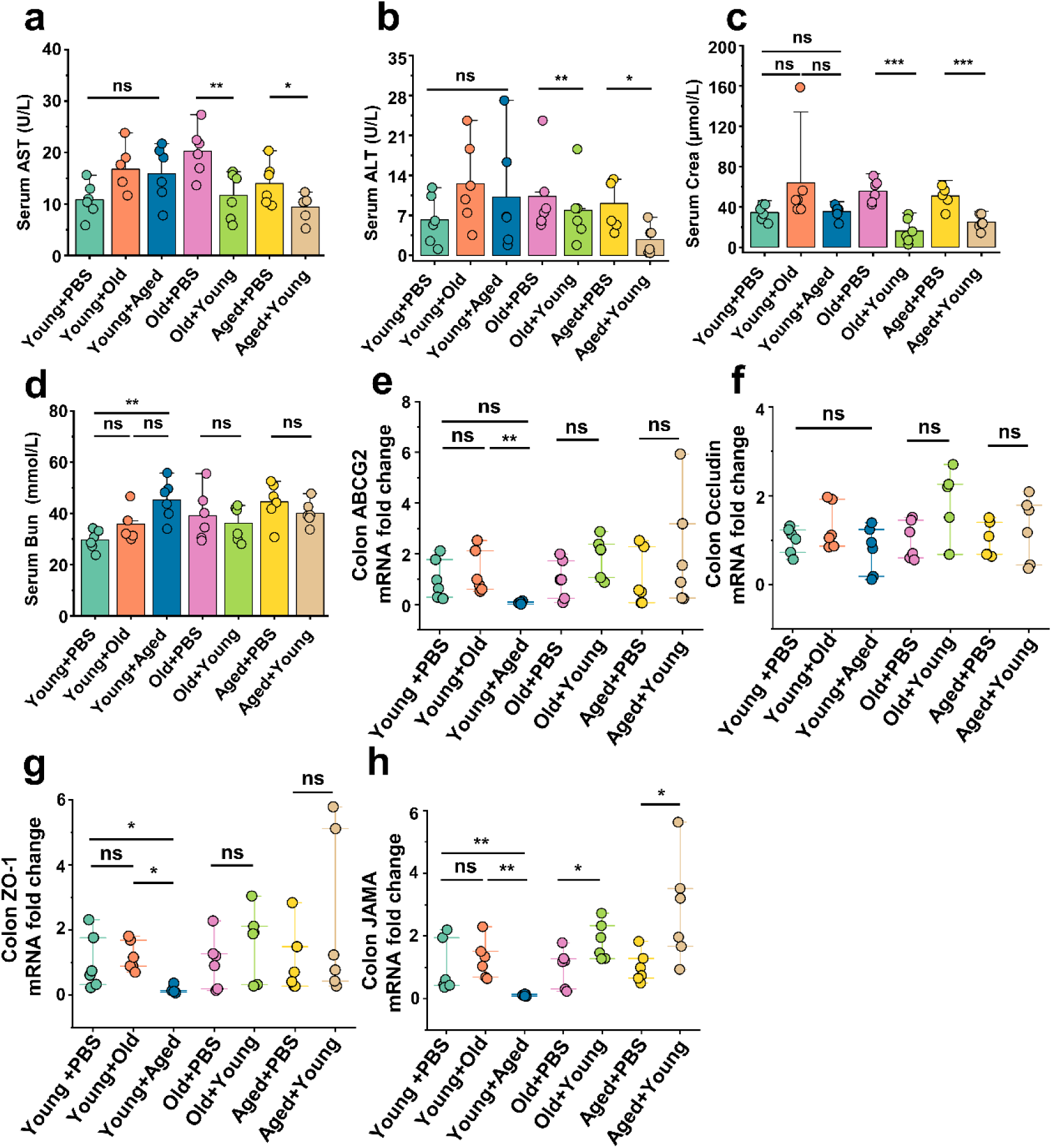
(a and b) Serum AST (a) and ALT (b) concentrations in the cross-age fecal microbiota transplantation group and its control group (n=6). (c and d) Serum Crea (c) and BUN (d) concentrations in the cross-age fecal microbiota transplantation group and its control group (n=6). (e-h) Relative colonic gene expression in the indicated groups by qPCR (n = 6), including ABCG2 (e), Occludin (f), ZO-1 (g) and JAMA (h). Values are presented as the mean ± SEM. Differences were assessed by t-test or One-Way ANOVA and denoted as follows: *p < 0.05, **p < 0.01, and ***p < 0.001, “ns” indicates no significant difference between groups.

**Supplementary Fig 4.**
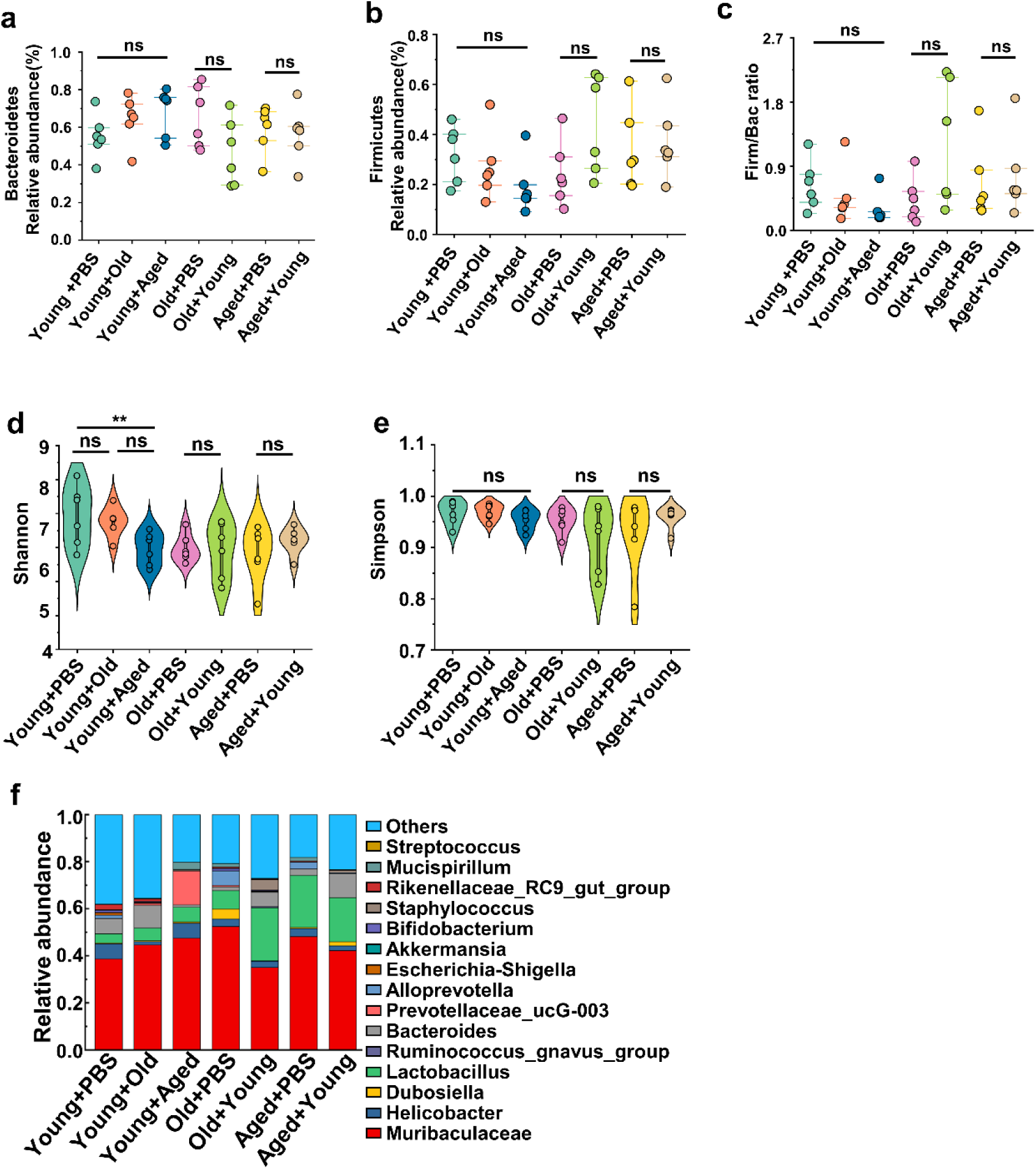
(a) Bacteroidetes relative abundance of the indicated groups (n=6). (b) Firmicutes relative abundance of the indicated groups (n=6). (c) The Firm/Bac ratio in the different groups (n=6). (d-e) Alpha diversity indices including Shannon (d) and Simpson (e) index in the indicated groups (n=6). (f) Bacterial composition at the genus levels (top 15) of the indicated groups(n=6). Values are presented as the mean ± SEM. Differences were assessed by t-test or One-Way ANOVA and denoted as follows: *p < 0.05, **p < 0.01, and ***p < 0.001, “ns” indicates no significant difference between groups.

**Supplementary Fig 5.**
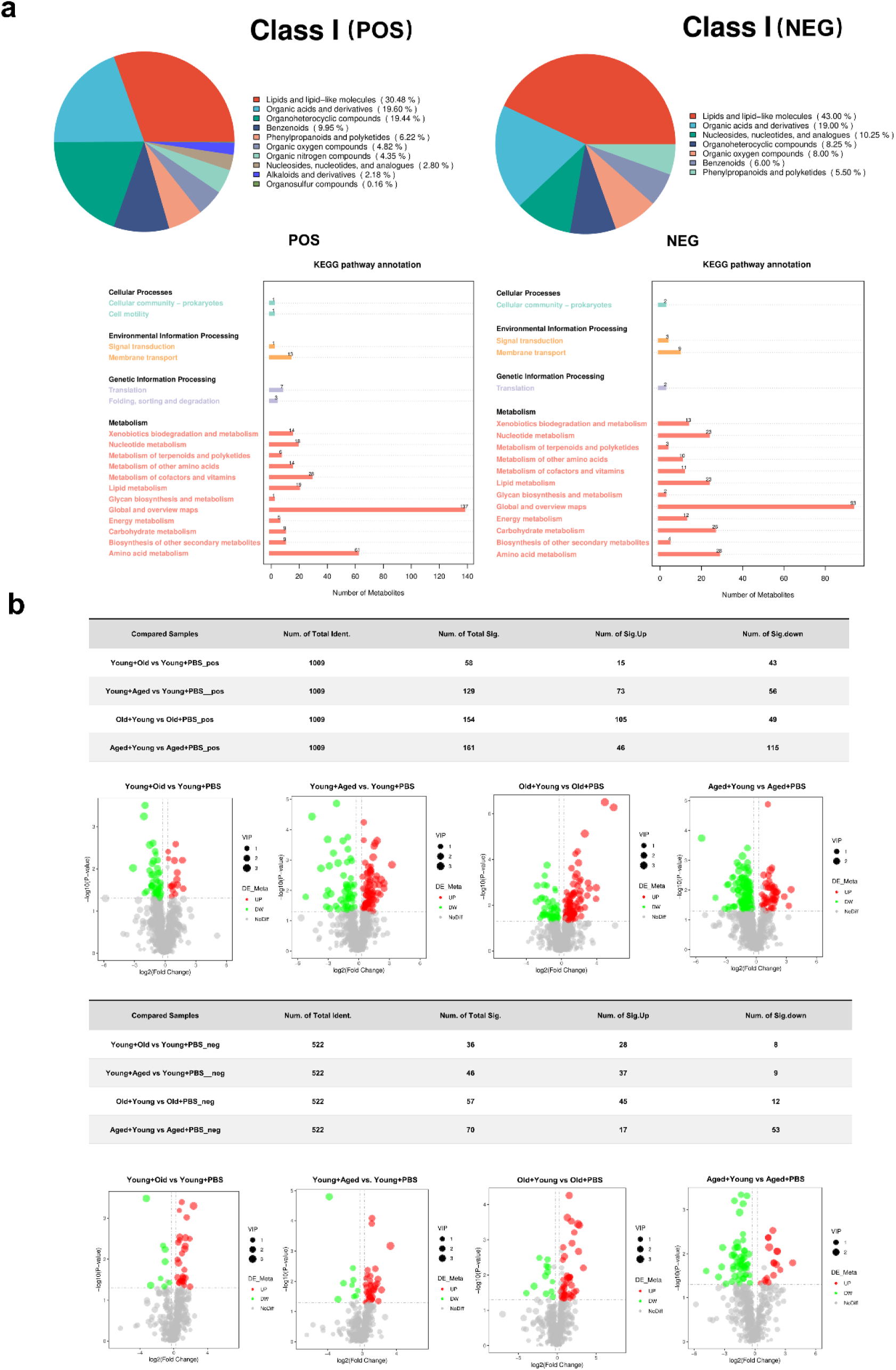
(a) Metabolite classification of various groups. (b) Volcano plots of differential metabolites in each group. The horizontal axis of volcano plot represents the log_2_ (Fold Change) in metabolite abundance between different groups, while the vertical axis of volcano plot represents the significance level (-log_10_ (P-value)) of the differences. Each point on the volcano plot represents a metabolite, with significantly upregulated metabolites depicted as red points and significantly downregulated metabolites depicted as green points. The size of the circles corresponds to the VIP (Variable Importance in Projection) value.

**Supplementary Fig 6.**
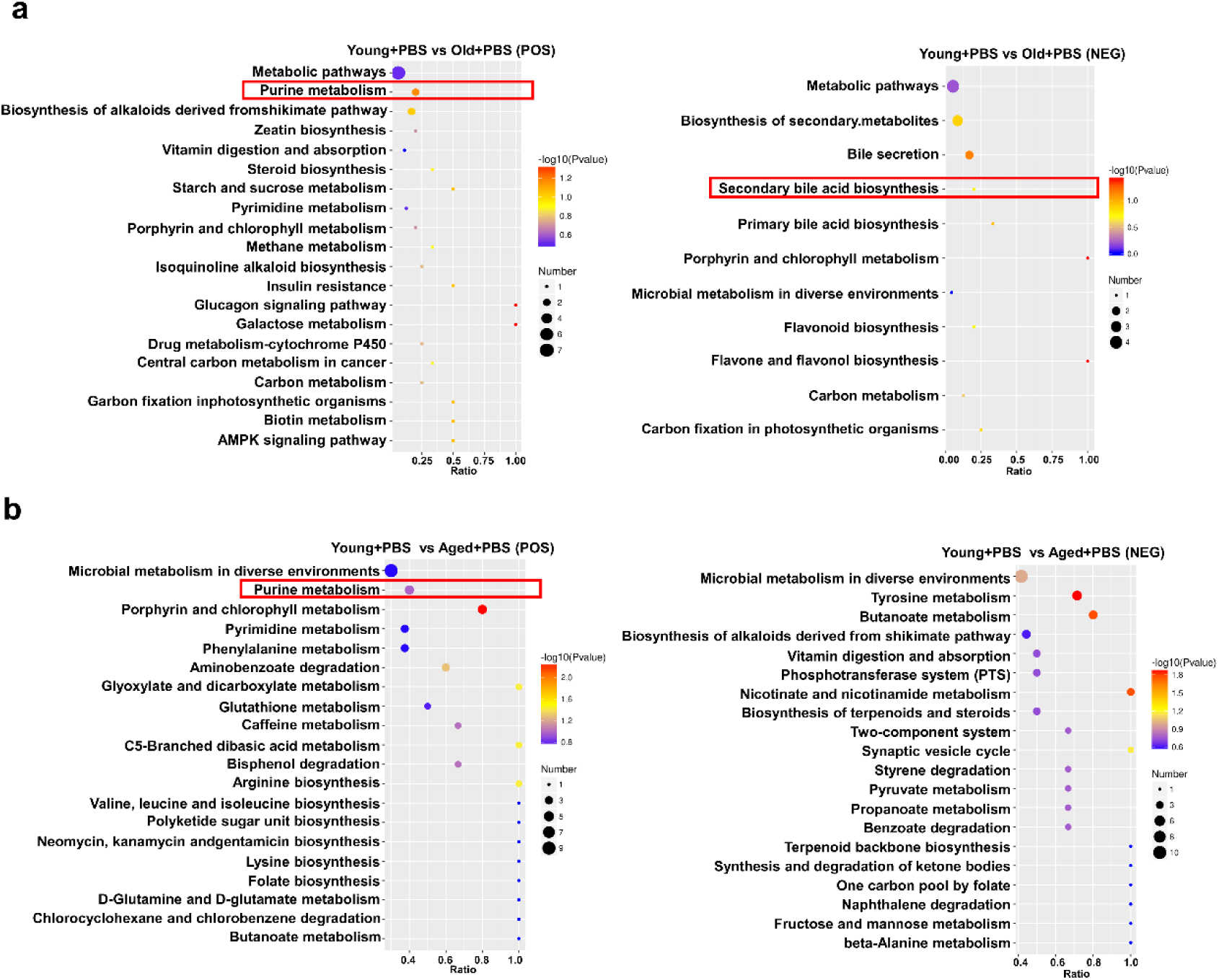
(a) Comparison of differential metabolite volcano plots and KEGG enrichment bubble plots between Young+PBS and Old+PBS. (b) Comparison of differential metabolite volcano plots and KEGG enrichment bubble plots between Young+PBS and Aged+PBS. To visualize the differential metabolites resulting from pairwise comparisons, we generated volcano plots that provide a clear representation of the upregulation and downregulation of metabolites, especially those with significant fold change differences. The enrichment analysis was performed at the KEGG Pathway level using a hypergeometric test, as shown in the figure below. The pathways that were significantly enriched in the differential metabolites compared to the background of all identified metabolites. Pathway enrichment analysis enables us to determine the major biochemical metabolic pathways and signaling transduction pathways that are implicated by the differential metabolites.

**Supplementary Fig 7.**
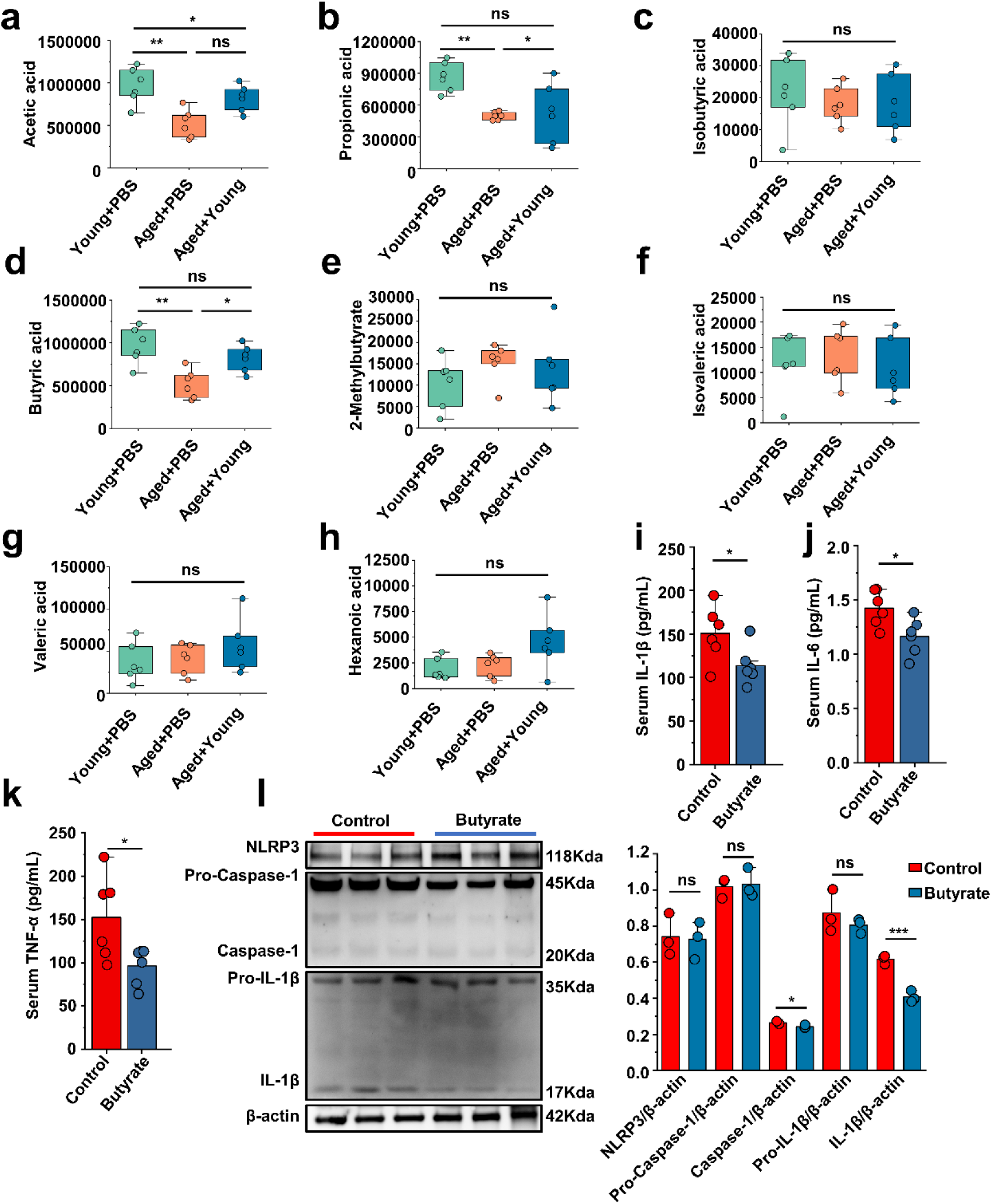
(a-h) The difference analysis of the detected short chain fatty acids, including acetic acid (a), propionic acid (b), isobutyric acid (c), butyric acid (d), 2-merhylbutyrate (e), isovaleric acid (f), valeric acid (g), hexanoic acid (h). (i-k) The serum concentrations of IL-1β (i), IL-6 (j) and TNF-α (k) inflammatory parameters were measured in the indicated mice (n=6). (l) Representative western blot images and band density of peritoneal cells NLRP3 pathways proteins (n=3) Values are presented as the mean ± SEM. Differences were assessed by t-test or One-Way ANOVA and denoted as follows: *p < 0.05, **p < 0.01, and ***p < 0.001, “ns” indicates no significant difference between groups.

**Supplementary Fig 8.**
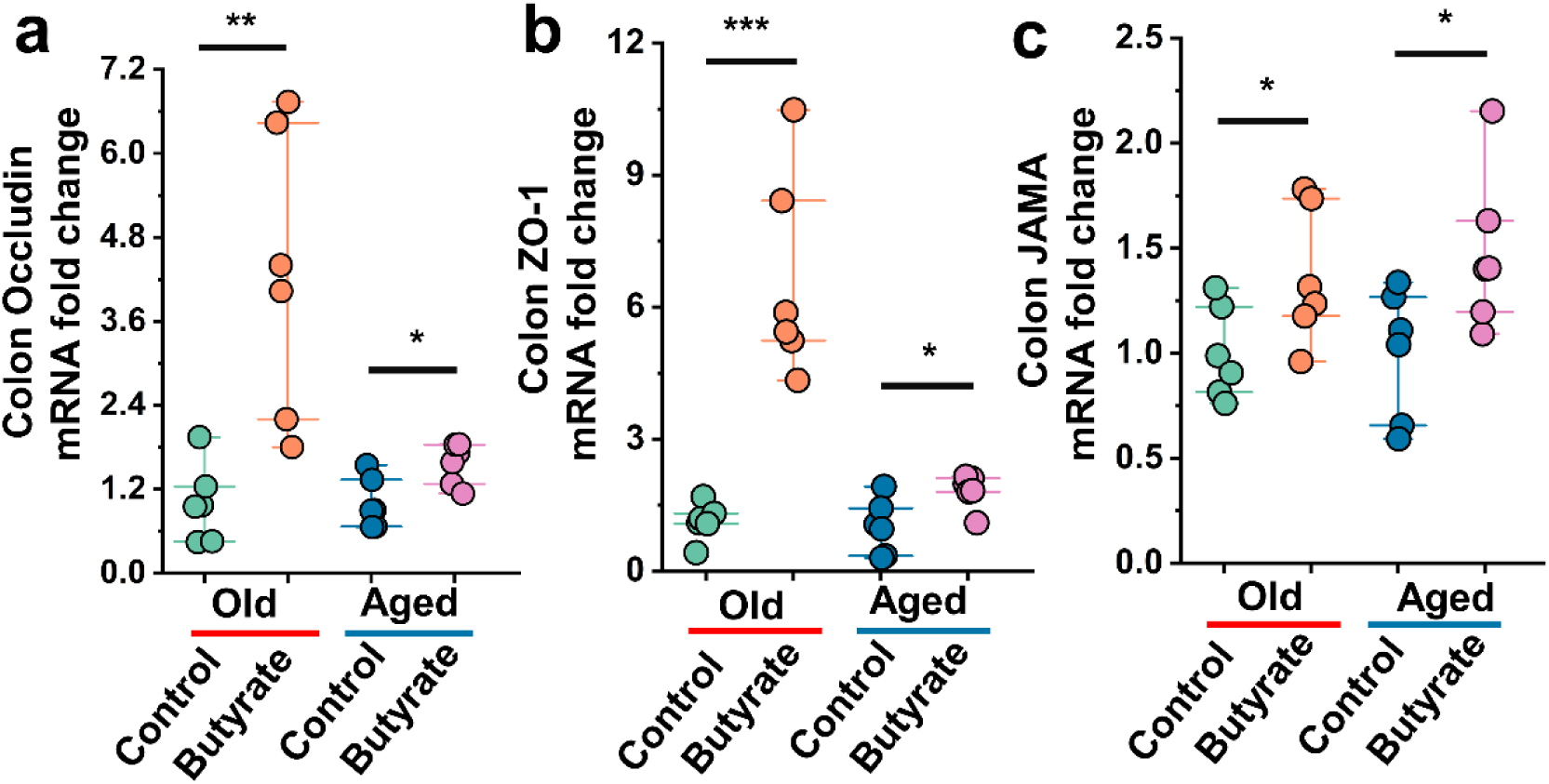
(a-c) Relative colonic gene expression in the indicated groups by qPCR (n = 6), including Occludin (a), ZO-1 (b) and JAMA (c). Values are presented as the mean ± SEM. Differences were assessed by t-test or One-Way ANOVA and denoted as follows: *p < 0.05, **p < 0.01, and ***p < 0.001, “ns” indicates no significant difference between groups.

## Notes

https://db.cngb.org/cnsa/project/CNP0004751_46c178da/reviewlink/

### Competing Interest Statement

The authors have declared no competing interest.

### Summary of Updates

The content in Supplement 4 has been adjusted, and modifications have been made to the discussion section.

