## Supplementary material 1 for "Microbiota from young mice counteracts susceptibility to age-related gout through modulating butyric acid levels in aged mice"

**Supplemental Information 1**

**16S rDNA amplicon sequencing**

1. **Extraction of genome DNA**

The CTAB/SDS method was used to extract the total genome DNA in samples. DNA concentration and purity were monitored on 1% agarose gels. According to the concentration, DNA was diluted to 1ng/µL with sterile water.

2. **Amplicon Generation**

16S rRNA/18SrRNA/ITS genes in distinct regions (16S V4/16S V3/16S V3-V4/16S V4-V5, 18S V4/18S V9, ITS1/ITS2, Arc V4) were amplified with specific primer (e.g. 16S V4: 515F- 806R, 18S V4: 528F-706R, 18S V9: 1380F-1510R, et. al) and barcodes. All PCR mixtures contained 15 µL of Phusion® High-Fidelity PCR Master Mix (New England Biolabs), 0.2 µM of each primer and 10ng target DNA, and cycling conditions consisted of a first denaturation step at 98℃ for 1 min, followed by 30 cycles at 98℃ (10s), 50℃(30s) and 72℃(30s) and a final 5min extension at 72℃.

3. **PCR Products quantification and qualification**

Mix an equal volume of 1X loading buffer (contained SYB green) with PCR products and perform electrophoresis on 2% agarose gel for DNA detection. The PCR products were mixed inequal proportions, and then Qiagen Gel Extraction Kit (Qiagen, Germany) was used to purify the mixed PCR products.

4. **Library preparation and sequencing**

Following manufacturer’s recommendations, sequencing libraries were generated with NEBNext® Ultra™ IIDNA Library Prep Kit (Cat No. E7645). The library quality was evaluated on the Qubit@ 2.0 Fluorometer (Thermo Scientific) and Agilent Bioanalyzer 2100 system. Finally, the library was sequenced on an Illumina NovaSeq platform and 250 bp paired-end reads were generated.

**Data analysis**

**1. Paired-end reads merged and quality control**

1.1 Data Split Paired-end reads were assigned to samples based on their unique barcodes and were truncated by cutting off the barcodes and primer sequences.

1.2 Paired-end reads merged Paired-end reads were merged using FLASH (Version 1.2.11, http://ccb.jhu.edu/software/FLASH/)[1], a very fast and accurate analysis tool designed to merge paired-end reads when at least some of the reads overlap with the reads generated from the opposite end of the same DNA fragment, and the splicing sequences were called Raw Tags.

1.3 Data Filtration Quality filtering on the raw tags were performed using the fastp (Version 0.20.0) software to obtain high-quality Clean Tags.

1.4 Chimera Removal The Clean Tags were compared with the reference database (Silva database https://www.arb- silva.de/ for 16S/18S, Unite database https://unite.ut.ee/ for ITS) using Vsearch (Version 2.15.0) to detect the chimera sequences, and then the chimera sequences were removed to obtain the Effective Tags[2].

**2. ASVs Denoise and Species annotation**

2.1 ASVs Denoise For the Effective Tags obtained previously, denoise was performed with DADA2 or deblur module in the QIIME2 software (Version QIIME2-202006) to obtain initial ASVs (Amplicon Sequence Variants) (default: DADA2), and then ASVs with abundance less than 5 were filtered out[3].

2.2 Species Annotation Species annotation was performed using QIIME2 software. For 16S/18S, the annotation database is Silva Database, while for ITS, it is Unite Database.

2.3 Phylogenetic Relationship Construction In order to study phylogenetic relationship of each ASV and the differences of the dominant species among different samples(groups), multiple sequence alignment was performed using QIIME2 software.

2.4 Data Normalization The absolute abundance of ASVs was normalized using a standard of sequence number corresponding to the sample with the least sequences. Subsequent analysis of alpha diversity and beta diversity were all performed based on the output normalized data.

3. Alpha Diversity In order to analyze the diversity, richness and uniformity of the communities in the sample, alpha diversity was calculated from 7 indices in QIIME2, including Observed_otus, Chao1, Shannon, Simpson, Dominance, Good’s coverage and Pielou_e.

Three indices were selected to identify community richness:

Observed_otus – the number of observed species ([http://scikit-bio.org/docs/latest/generated/s kbio.diversity.alpha.observed_otus.html](http://scikit-bio.org/docs/latest/generated/s%20kbio.diversity.alpha.observed_otus.html));

Chao – the Chao1 estimator (<http://scikit-bio.org/docs/latest/generated/skbio.diversity.alpha>. chao1.html#skbio. diversity. alpha. chao1);

Dominance – the Dominance index (<http://scikit-bio.org/docs/latest/generated/skbio.diversity>. alpha.dominance.html#skbio. diversity. alpha. dominance);

Two indices were used to identify community diversity:

Shannon – the Shannon index (<http://scikit-bio.org/docs/latest/generated/skbio.diversity.alpha.shannon.html#skbio.diversity.alpha.shannon>);

Simpson–the Simpson index (<http://scikit-bio.org/docs/latest/generated/skbio.diversity.alpha.simpson.html#skbio.diversity.alpha.simpson>);

One indice was used to calculate sequencing depth:

Coverage – the Good’s coverage (<http://scikit-bio.org/docs/latest/generated/skbio.diversity.al> pha.goods_coverage.html#skbio.diversity.alpha.goods_coverage);

One indice was used to calculate species evenness:

Pielou_e – Pielou’s evenness index (<http://scikit-bio.org/docs/latest/generated/skbio.diversity>. alpha.pielou_e.html#skbio. diversity. alpha. pielou_e).

**3. Beta Diversity**

In order to evaluate the complexity of the community composition and compare the differences between samples(groups), beta diversity was calculated based on weighted and unweighted unifrac distances in QIIME2.

Cluster analysis was performed with principal component analysis (PCA), which was applied to reduce the dimension of the original variables using the ade4 package and ggplot2 package in Rsoftware (Version 3.5.3).

Principal Coordinate Analysis (PCoA) was performed to obtain principal coordinates and visualize differences of samples in complex multi-dimensional data. A matrix of weighted or unweighted unifrac distances among samples obtained previously was transformed into a newset of orthogonal axes, where the maximum variation factor was demonstrated by the first principal coordinate, and the second maximum variation factor was demonstrated by the second principal coordinate, and so on. The three-dimensional PCoA results were displayed using QIIME2package, while the two-dimensional PCoA results were displayed using ade4 package and ggplot2package in R software (Version 2.15.3).

To study the significance of the differences in community structure between groups, the adonis and anosim functions in the QIIME2 software were used to do analysis. To find out the significantly different species at each taxonomic level (Phylum, Class, Order, Family, Genus, Species), the R software (Version 3.5.3) was used to do MetaStat and T-test analysis. The LEfSe software (Version 1.0) was used to do LEfSe analysis (LDA score threshold: 4) to find out the biomarkers. Further, to study the functions of the communities in the samples and find out the different functions of the communities in the different groups, the PICRUSt2 software (Version 2.1.2-b) was used for function annotation analysis.
