## Supplementary material 2 for "Microbiota from young mice counteracts susceptibility to age-related gout through modulating butyric acid levels in aged mice"

**Supplemental Information 2**

**Untargeted Metabolomics - Materials and Methods**

1. **Metabolites Extraction**

Fecal samples (100 mg) were individually grounded with liquid nitrogen and the homogenate was resuspended with prechilled 80% methanol by well vortex. The samples were incubated on ice for 5 min and then were centrifuged at 15,000 g, 4°Cfor 20 min. Some of supernatant was diluted to final concentration containing 53%methanol by LC-MS grade water. The samples were subsequently transferred to afresh Eppendorf tube and then were centrifuged at 15000 g, 4°C for 20 min. Finally, the supernatant was injected into the LC-MS/MS system analysis[1].

2. **UHPLC-MS/MS Analysis**

UHPLC-MS/MS analyses were performed using a Vanquish UHPLC system (ThermoFisher, Germany) coupled with an Orbitrap Q Exactive TMHF-Xmass spectrometer (Thermo Fisher, Germany) in Novogene Co., Ltd. (Beijing, China). Samples were injected onto a Hypesil Gold column (100×2.1 mm, 1.9μm) using a 17-min linear gradient at a flow rate of 0.2mL/min. The eluents for the positive polarity mode were eluent A (0.1% FA in Water) and eluent B (Methanol). The eluents for the negative polarity mode were eluent A (5mMammoniumacetate, pH9.0) and eluent B (Methanol). The solvent gradient was set as follows: 2%B, 1.5 min; 2-85% B, 3 min; 85-100% B, 10 min；100-2% B, 10.1 min；2%B, 12 min. QExactive ^TM^ HF-X mass spectrometer was operated in positive/negative polarity mode with spray voltage of 3.5 kV, capillary temperature of 320°C, sheath gas flowrate of 35 psi and aux gas flow rate of 10 L/min, S-lens RF level of 60, Aux gas heater temperature of 350°C.

3. **Data processing and metabolite identification**

The raw data files generated by UHPLC-MS/MS were processed using the Compound Discoverer 3.1 (CD3.1, ThermoFisher) to perform peak alignment, peak picking, and quantitation for each metabolite. The main parameters were set as follows: retention time tolerance, 0.2 minutes; actual mass tolerance, 5ppm; signal intensity tolerance, 30%; signal/noise ratio, 3; and minimum intensity, et al. After that, peak intensities were normalized to the total spectral intensity. The normalized data was used to predict the molecular formula based on additive ions, molecular ion peaks and fragment ions. And then peaks were matched with the mzCloud(https://www.mzcloud.org/)，mzVault and MassList database to obtain the accurate qualitative and relative quantitative results. Statistical analyses were performed using the statistical software R (R version R-3.4.3), Python (Python 2.7.6 version) and CentOS (CentOS release 6.6). When data were not normally distributed, standardize according to the formula: sample raw quantitation value / (The sum of sample metabolite quantitation value / The sum of QC1 sample metabolite quantitation value) to obtain relative peak areas; And compounds whose CVs of relative peak areas in QC samples were greater than 30% were removed, and finally the metabolites' identification and relative quantification results were obtained.

4. Data Analysis

These metabolites were annotated using the KEGG database (https://www.genome.jp/kegg/pathway.html)， HMDB database (https://hmdb.ca/ metabolites) and LIPID Maps database (<http://www.lipidmaps.org/>). Principal components analysis (PCA) and Partial least squares discriminant analysis (PLS-DA) were performed at metaX[2] (a flexible and comprehensive software for processing metabolomics data). We applied univariate analysis (t-test) to calculate the statistical significance (P-value). The metabolites with VIP > 1 and P-value< 0.05 and foldchange≥2 or FC≤0.5 were differential metabolites. Volcano plots were used to filter metabolites of interest which based on log_2_ (Foldchange) and-log_10_ (p-value) of metabolites by ggplot2 in R language.

For clustering heat maps, the data were normalized using z-scores of the intensity areas of differential metabolites and were plotted by Pheatmap package in R language. The correlation between differential metabolites were analyzed by cor () in R language (method=pearson). Statistically significant of correlation between differential metabolites were calculated by cor. mtest () in R language. P-value < 0.05 was considered as statistically significant and correlation plots were plotted by corrplot package in R language. The functions of these metabolites and metabolic pathways were studied using the KEGG database. The metabolic pathways enrichment of differential metabolites was performed, when ratio was satisfied by x/n >y/N, metabolic pathway was considered as enrichment, when P-value of metabolic pathway < 0.05, metabolic pathway was considered as statistically significant enrichment.
