## Supplementary material 3 for "Microbiota from young mice counteracts susceptibility to age-related gout through modulating butyric acid levels in aged mice"

**Supplemental Information 3**

**1.Reagents and instruments**

**Equipment**

An ultra-high-performance liquid chromatography coupled to tandem mass spectrometry (UHPLC-MS/MS) system (Vanquish™ Flex UHPLC-TSQ Altis™, Thermo Scientific Corp., Germany).

**Materials and reagents**

All the 11 SCFA standards were obtained from ZZ Standards Co., LTD. (Shanghai, China). Methanol (Optima LC-MS), acetonitrile (Optima LC-MS), ammonium acetate, isopropanol (Optima LC-MS) were purchased from Thermo-Fisher Scientific (FairLawn, NJ, USA). Ultrapure water was purchased from Millipore (MA, USA).

**2. Standard solution preparation**

The stock solution of individual SCFA was mixed and prepared in SCFA-free matrix to obtain a series of SCFA calibrators. Certain concentrations of Isotope standard were compounded and mixed as Internal Standard (IS). The stock solution of all of these and working solution were stored in refrigerator of -20°C.

| Number | Name |
| --- | --- |
| 1 | Acetic acid |
| 2 | Propionic acid |
| 3 | Isobutyric acid |
| 4 | Butyric acid |
| 5 | 2-Methylbutyrate |
| 6 | Isovaleric acid |
| 7 | Valeric acid |
| 8 | Hexanoic acid |
| 9 | 2-Methylvalerate |
| 10 | 3-Methylvalerate |
| 11 | 4-Methylvaleric acid |

**3.Metabolites extraction**

The samples (100 mg) were resuspended with liquid nitrogen and then diluted to 100 times samples. Then 100 μL of them were taken respectively and homogenized with 400 μL of methanol (80%) and centrifuged to remove the protein. The supernatant was added to derivatization reagent (150 μL) and derivatized at 40℃ for 40 min. Then supernatant (190 μL) was homogenized with 10 μL mixed internal standard solution. Finally, injected into the LC-MS/MS system for analysis.

4. **LC-MS method**

An ultra-high-performance liquid chromatography coupled to tandem mass spectrometry (UHPLC-MS/MS) system (Vanquish™ Flex UHPLC-TSQ Altis™, Thermo Scientific Corp., Germany) was used to quantitate SCFA in Novogene Co., Ltd. (Beijing, China). Separation was performed on a Waters ACQUITY UPLC BEH C18 column (2.1×100mm, 1.7μm) which was maintained at 40°C. The mobile phase, consisting of 10 mM ammonium acetate in water (solvent A) and acetonitrile: isopropanol (1:1) (solvent B), was delivered at a flow rate of 0.30 mL/min. The solvent gradient was set as follows: initial 25% B, 2.5min; 25-30% B, 3 min; 30-35% B, 3.5 min; 35-38% B, 4 min; 38-40% B, 4.5 min; 40-45% B, 5 min; 45-50% B, 5.5min; 50-55% B, 6.5 min; 55-58% B, 7 min; 58-70% B, 7.5min; 70-100% B, 7.8 min; 100-25% B, 10.1min; 25% B, 12 min. The mass spectrometer was operated in negative multiple reaction mode (MRM) mode. Parameters were as follows: IonSpray Voltage (-4500V), Sheath Gas (35psi), Ion Source Temp (550°C), Auxiliary Gas (50psi), Collision Gas (55psi).
