## Supplementary material 4 for "Microbiota from young mice counteracts susceptibility to age-related gout through modulating butyric acid levels in aged mice"

1. **The results of IL-6 and TNF-α in foot tissue of all age groups.**

The levels of IL-6 was significantly elevated in the Old and Aged groups, after MSU stimulated. While no significant difference in foot tissue TNF-α concentration was found between Aged and Young groups, an significant upward trend was observed in Old group.

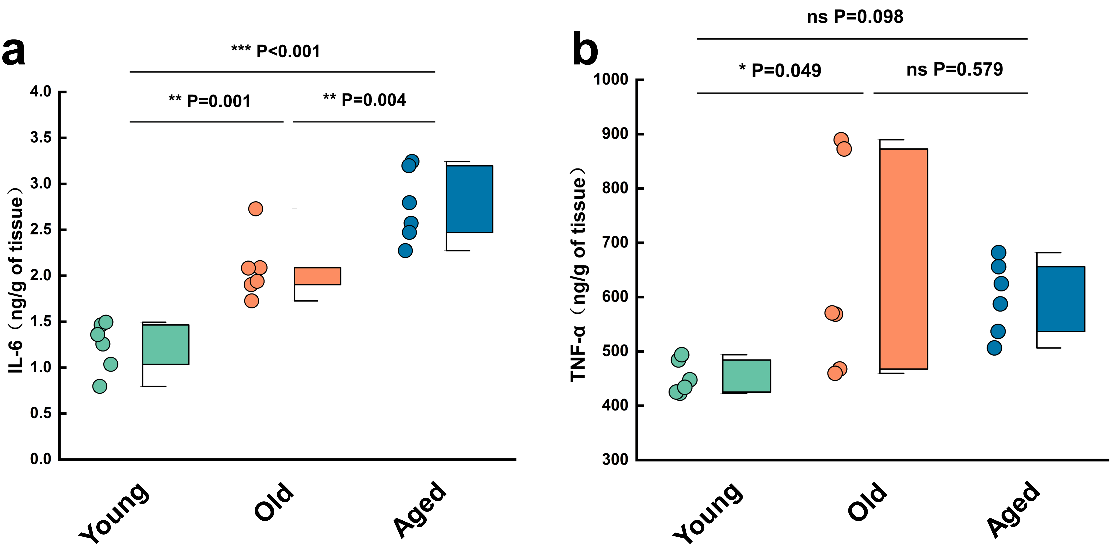

(a, b) The IL-6 (a) and TNF-αcontent (b) in foot tissue of all age groups (n=6).

Values are presented as the mean ± SEM. Differences were assessed by t-test or One-Way ANOVA and denoted as follows: *p < 0.05, **p < 0.01, and ***p < 0.001, “ns” indicates no significant difference between groups.

2. **The IL-1βcontent of Abx-untreated and untreated groups included at all ages**

There was no significant difference in the content of IL-1β in the foot tissues of mice of all ages without MSU stimulation, whether treated with antibiotics or untreated.

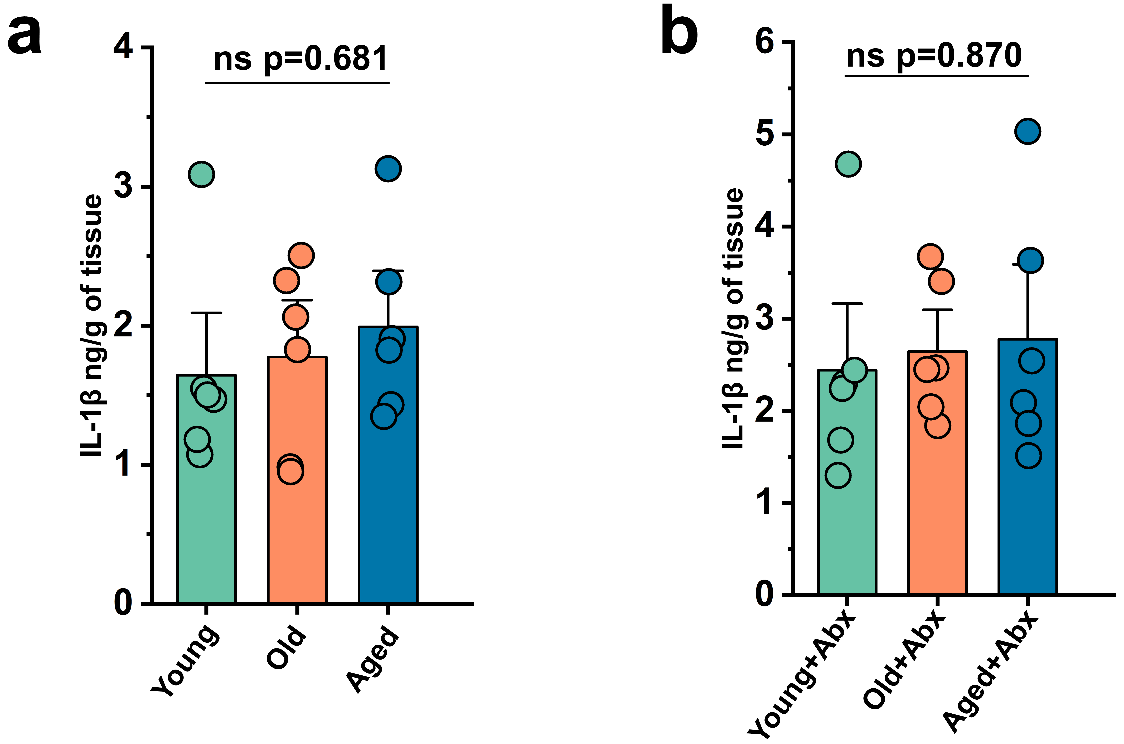

(a, b) The IL-1βcontent of untreated (a) and Abx-untreated (b) groups included at all ages (n=6).

Values are presented as the mean ± SEM. Differences were assessed by t-test or One-Way ANOVA and denoted as follows: *p < 0.05, **p < 0.01, and ***p < 0.001, “ns” indicates no significant difference between groups.

3. **Immunohistochemical slices of Zo-1 and Occludin in all groups**

To assess the expression of tight junction (TJ) proteins in the colon, immunohistochemical assays were conducted employing the SAP (Mouse/Rabbit) IHC Kit (MXB, China), as described by Zhao et al(1). Following dewaxing with xylene and hydration with alcohol, antigen retrieval was performed using sodium citrate. Subsequently, the sections were subjected to endogenous peroxidase inhibition for 40 minutes at room temperature, followed by three 5-minute washes with PBS. Nonimmune goat serum was applied for 40 minutes at RT to block non-specific binding sites. Overnight incubation at 4°C was then carried out with primary antibodies against ZO-1, and Occludin. The unbound primary antibodies were removed with PBS washes, followed by treatment with goat anti-rabbit IgG for 30 minutes at RT and subsequent incubation with horseradish peroxidase for 20 minutes. All sections were processed with a chromogenic developer for 5 minutes and nuclei were stained with hematoxylin for 2 minutes. After differentiation with 1% nitric acid alcohol, treatment with ammonium hydroxide, and dehydration, the sections were mounted with neutral resins and examined using an optical microscope (Olympus, Tokyo, Japan) and analyzed with Caseviewer2.0 software. Hematoxylin staining of the cell nuclei results in a blue color, while DAB staining yields a positive expression that appears as a brownish-yellow.

Our study, showed that compared with those of the same-age control group, the colon levels of ZO-1 and Occludin protein were significantly decreases in the Young+Old and Young+Aged groups, whereas the Old+Young and Aged+Young groups exhibited modest elevated in these levels.

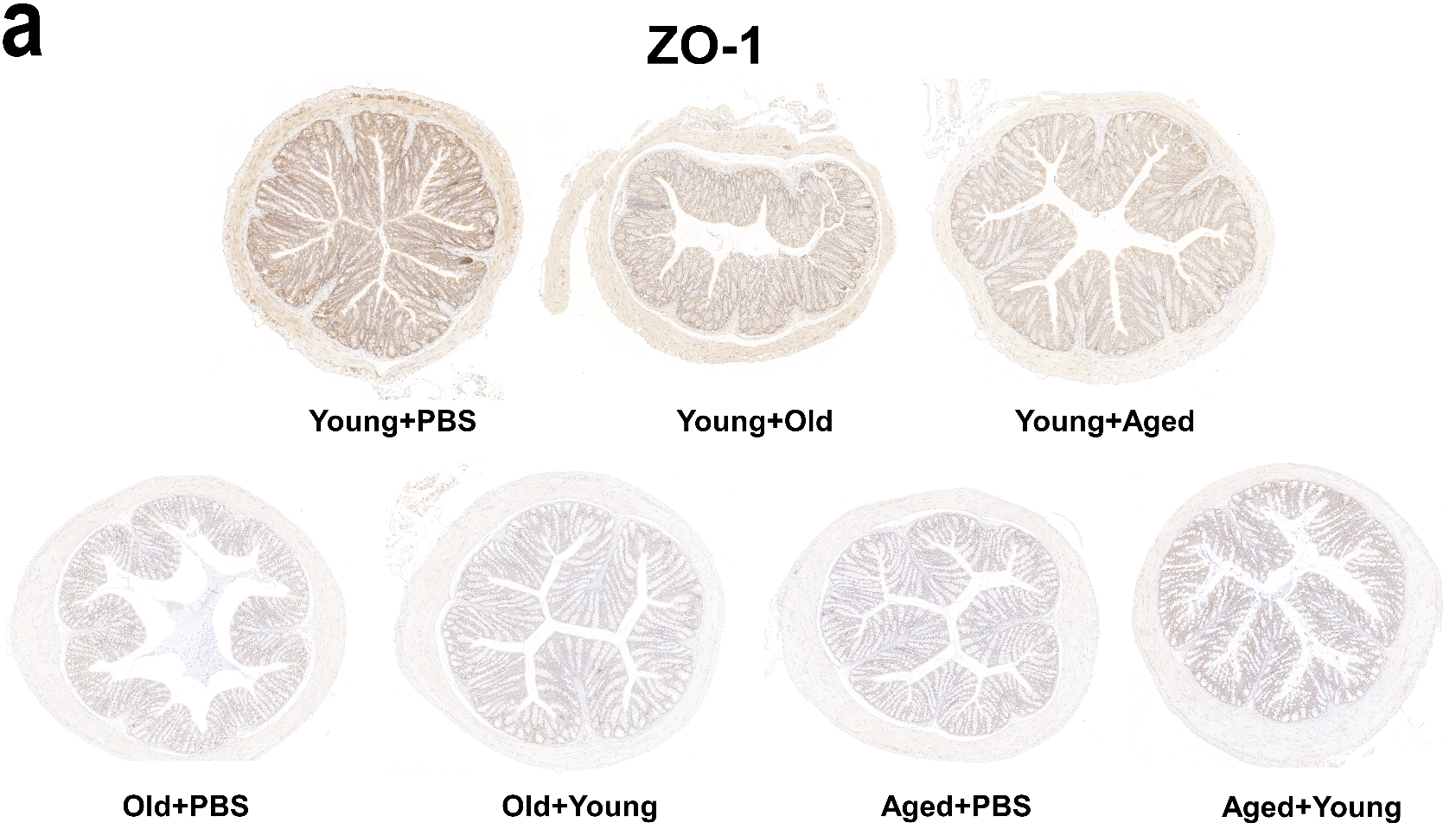

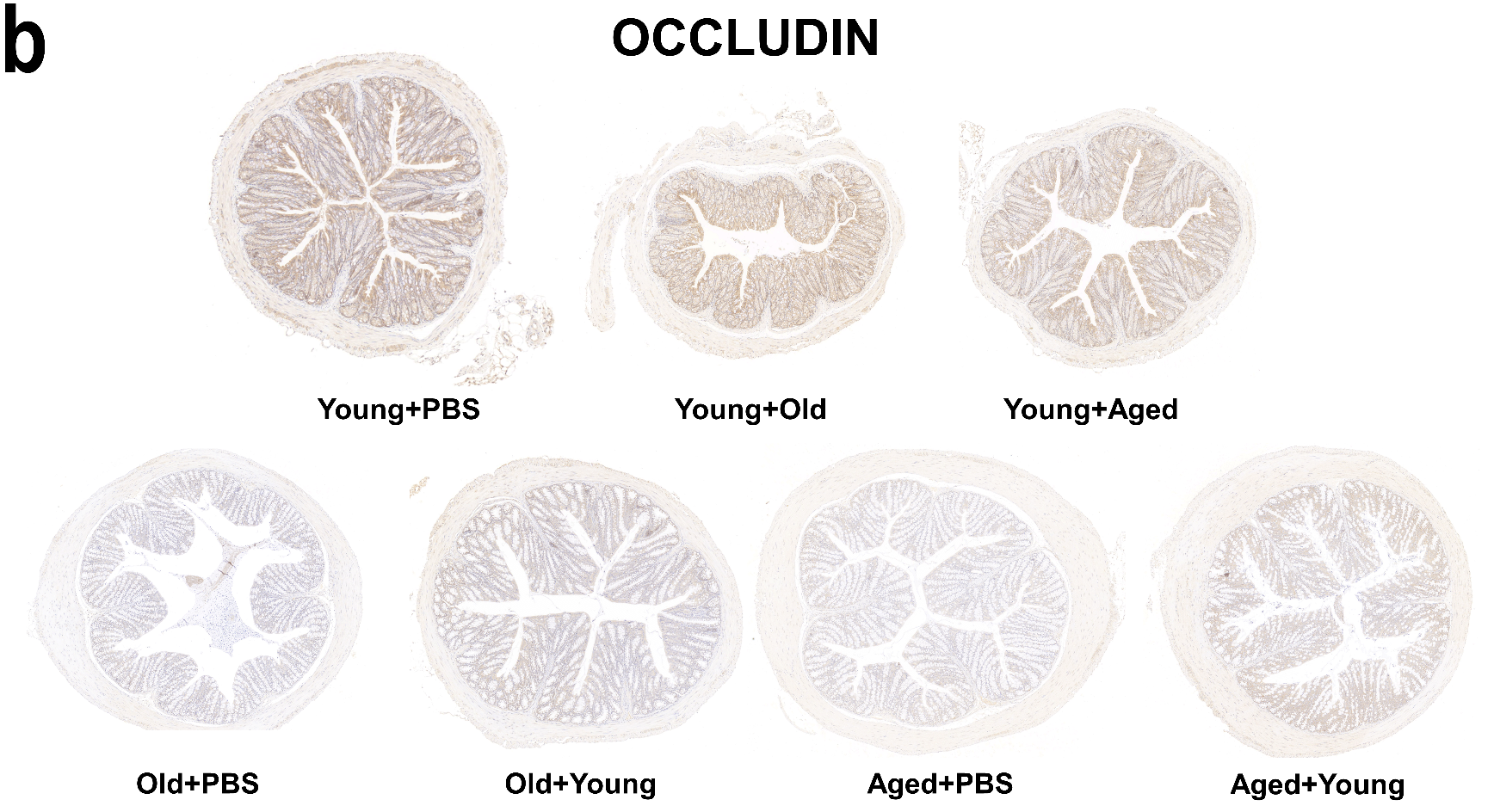

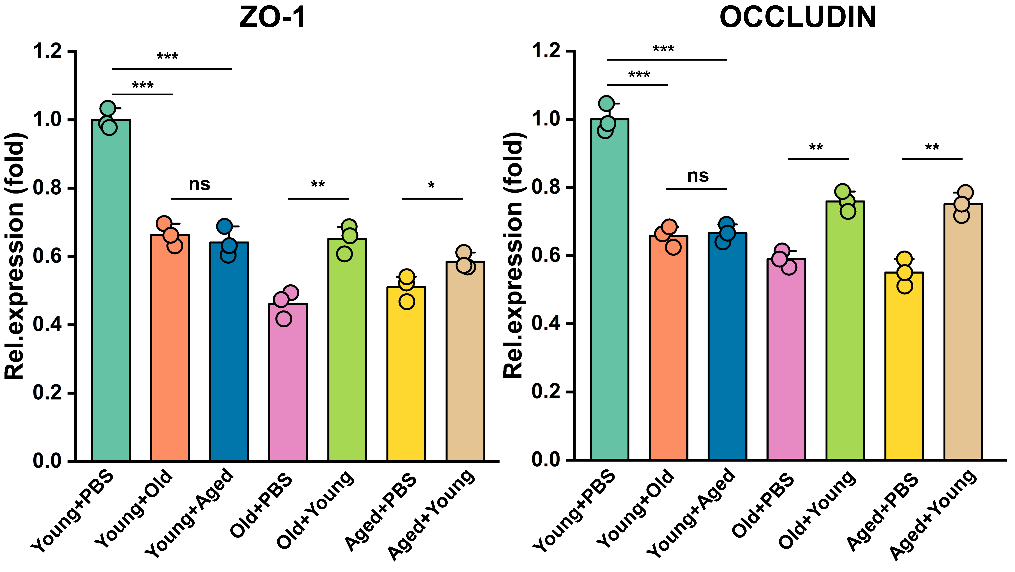

(a, b) Immunohistochemical slices of Zo-1 (a) and Occludin (b). n=3, Scale bars 1000 μm and 5x magnification. Values are presented as the mean ± SEM. Differences were assessed by t-test or One-Way ANOVA and denoted as follows: *p < 0.05, **p < 0.01, and ***p < 0.001, “ns” indicates no significant difference between groups.

4. **Table 2 with a more intuitive visual format**

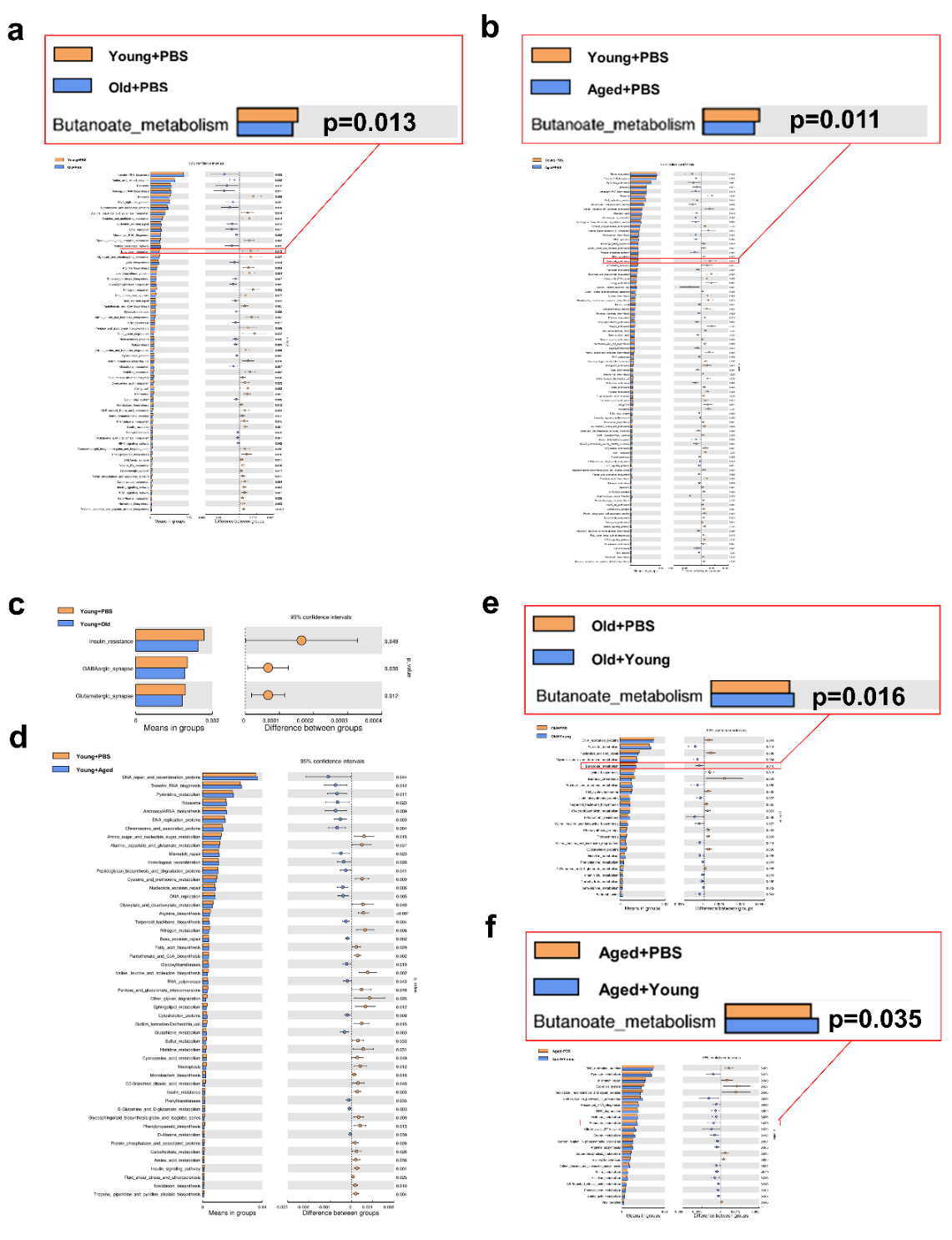

(a and b) Significant differences in gene categories between the Young+PBS group and the

Old+PBS group(a), and between the Young+PBS and Aged+PBS (b). Distinct gene categories were selected based on significant differences in gene categories at level 3 (t-test, p < 0.05).

(c and d) Significant differences in gene categories between the Young+PBS group and the

Young+Old group(c), and between the Young+PBS and Young+Aged (d). Distinct gene categories were selected based on significant differences in gene categories at level 3 (t-test, p < 0.05).

(e and f) Significant differences in gene categories between the Old+PBS group and the

Old+Young group(e), and between the Aged+PBS and Aged+Young (f). Distinct gene categories were selected based on significant differences in gene categories at level 3 (t-test, p < 0.05).

5. **The expression of uric acid-producing enzymes activity and uric acid transporters at the mRNA level across different age groups.**

The activity of GDA in the liver significantly differ from those of the Young group (b). Aged group demonstrates a higher level of KIM-1 mRNA expression relative to Young group (e). The Aged and Old groups showed lower mRNA expression levels of the uric acid excretion proteins OAT1 compared with the Young group (h). These are the results observed in mice across different age groups.

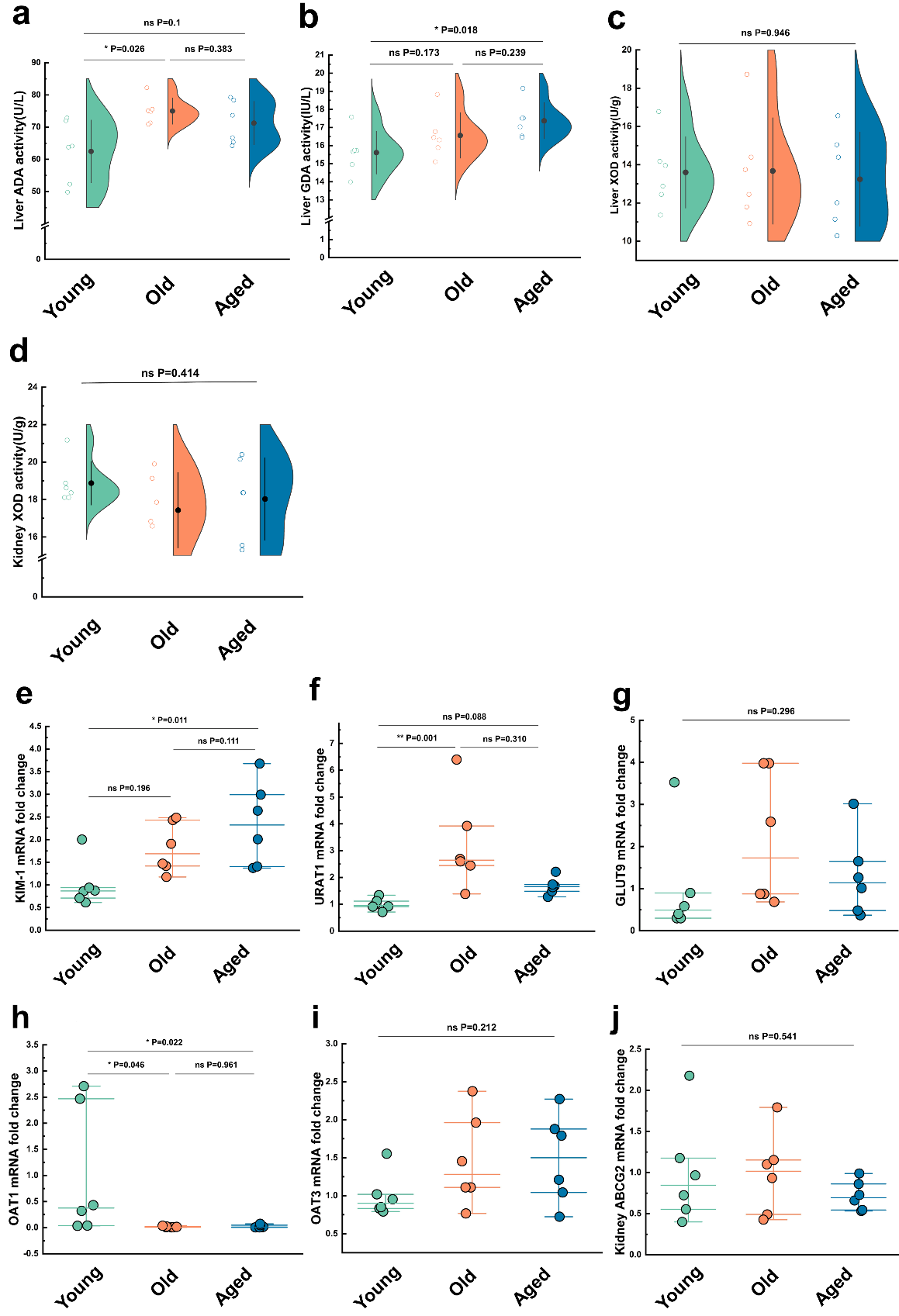

(a-c) The activity of uric acid-producing enzymes of liver in the all age groups. (n=6), including ADA (a), GDA (b) and XOD (c).

(d) The activity of XOD of kidney in the cross-age fecal microbiota transplantation group and its control group (n=6).

(e) Relative kidney injury molecule-1(KIM-1) expression in the indicated groups by qPCR (n = 6).

(f and g) Relative renal genes for uric acid reabsorption expression in the indicated groups by qPCR (n = 6), including URAT1 (f) and GLUT9 (g).

(h-j) Relative renal genes for uric acid excretion expression in the indicated groups by qPCR (n = 6), including OAT1 (h), OAT3 (i) and ABCG2 (j).

Values are presented as the mean ± SEM. Differences were assessed by t-test or One-Way ANOVA and denoted as follows: *p < 0.05, **p < 0.01, and ***p < 0.001, “ns” indicates no significant difference between groups.
